# Brain Substrates of Episodic Memory for Identity, Location, and Action Information: A Lesion-Behavior Mapping Study

**DOI:** 10.1101/2022.10.21.512938

**Authors:** Shir Ben-Zvi Feldman, Nachum Soroker, Daniel A. Levy

## Abstract

Brain networks supporting visual memory include extrastriate and other cortical regions associated with visual perception, which manifest domain-specific processing of “where,” “how,” and various aspects of “what” information. However, whether and how such specialization affects memory for these types of information is still a matter of debate. Functional neuroimaging studies point to dissociable as well as common network components supporting the perception and memory of different aspects of visual information. In the current neuropsychological study, we assess the impact of stroke lesion topography on recall of identity, location, and action of event participants, as assessed by the WMS-III Family Pictures subtest. We used voxel-based lesion-behavior mapping (VLBM) to identify brain lesions specifically implicated in memory deficits for each dimension. Behavioral analysis disclosed impaired performance by both right- and left-hemisphere damage patients, with lesions on each side yielding distinct effects. VLBM analysis revealed a bi-hemispheric network supporting these various aspects of visual memory. In the right hemisphere, the network includes frontal, parietal, and temporal cortical regions and the basal ganglia. In the left hemisphere, the network is more restricted, including visual association areas and medial temporal lobe regions. We further observed that a subset of these regions - those included in the ventral (“what”) stream, and in the putative core recollection network - is implicated in multiple aspects of visual memory, whereas other areas are specifically implicated in memory for specific aspects of the visual scene.

## 1. Introduction

Our visual experience of the world is an ongoing stream of distinct features of events unfolding simultaneously, such as the identity of the persons who take part in an episode, their relative positions, their intentions, and their actions, converging into a coherent and seemingly unified representation. Certain aspects of such information, encoded in our episodic memory, are crucial for everyday orientation in the world, social interactions, and even our ability to simulate future scenarios and plan ahead (Schacter, Benoit, & Szpunar, 2017). During episodic remembering, we recollect multiple perceptual features tied to an event that occurred in a specific spatio-temporal context (Clewett, DuBrow, & Davachi, 2019; Ranganath, 2010). One kind of episode which is arguably especially important to remember in ecological contexts is situations in which other people in our vicinity are engaged in meaningful activities and interactions. Regarding those kinds of events, we would like to be able to remember the identity of the participants, the actions they executed, and their locations, among other possible details.

What are the substrates of our memory abilities for such observed events? As remembering involves recapitulation of neural activity that occurred at the time of the original event (Johnson, McDuff, Rugg, & Norman, 2009; Staresina, Henson, Kriegeskorte, & Alink, 2012; Wing, Ritchey, & Cabeza, 2015), we might expect that remembering the identity of items or persons forming part of an experience would require reactivation and network integrity of ventral visual stream areas implicated in object and person representation, while remembering the locations of objects would be based on dorsal visual stream substrates of spatial representation (Gallivan & Goodale, 2018). Research efforts have indeed confirmed the reactivation of ventral stream representations in the context of episodic memory (Xue, 2018), with the most consistent activity across studies for experienced objects occurring in the fusiform gyrus (Schacter, Reimanti, Uecker, Polster, Yun & Cooper, 1995; Vaidya, Zhao, Desmond, & Gabrieli, 2002; Wheeler et al., 2000; Wheeler & Buckner, 2003) and parahippocampal gyrus (Schacter et al.,1995, Slotnick et al., 2003). Other regions activated during item identity memory were frontal regions, precuneus (Slotnick et al., 2003; Wheeler et al., 2000), lateral temporal regions (Slotnick et al., 2003), and middle occipital and lateral parietal regions (Wheeler et al., 2000). However, some studies have demonstrated activity related to visual item memory in both dorsal and ventral visual streams (Bone, Ahmad, & Buchsbaum, 2020; Buchsbaum, Lemire-Rodger, Fang, & Abdi, 2012; Moscovitch et al., 1995; Slotnick et al., 2003; Wheeler et al., 2000). In contrast to these functional imaging studies, there is a paucity of lesion studies of cortical regions implicated in object identity memory, with the exception of the report of Kessels, Postma, Kappelle, & de Haan (2000) of left parietal and occipital as well as right medial parietal and posterior parietal involvement.

Many studies have focused on memory for objects’ spatial location. A typical spatial-location paradigm entails the presentation of objects in various locations in a grid, in which participants are tested subsequently for their memory for the objects and their original locations. Early PET studies reported location memory-related activity in the right supramarginal gyrus, frontal, and lateral temporal regions (Köhler et al., 1998; Moscovitch et al., 1995), middle occipital gyrus, bilateral insula (Kohler et al., 1998) and inferior temporal and fusiform gyri (Moscovitch et al., 1995). Subsequently, fMRI studies testing source memory of spatial location have shown activity in the frontal cortex, lateral temporal regions (Slotnick et al., 2003; Ross & Slotnick, 2008), hippocampus, and superior parietal lobe (Ross & Slotnick, 2008). Another fMRI study has shown frontal and superior parietal activity during an object-location memory task (Hales & Brewer, 2013). Furthermore, the posterior parietal cortex was also implicated in object-location memory by lesion studies (Kessels et al., 2000; van Asselen et al., 2009). Two reviews of the literature of lesion studies (Postma, Kessels & van Asselen, 2008; Zimmermann & Eschen, 2017) and neuroimaging studies (Zimmermann & Eschen, 2017) have indicated that the cingulate cortex, the left superior parietal lobe, the right hippocampus and the posterior parietal cortex are involved in object-location perceptual processing; it remains to be determined whether those areas are equally involved in object location memory as well.

Actions in the context of events are represented and interpreted in the context of conceptual schemas, which can have a profound impact on our ability to remember those events (Alba & Hasher, 1983; Gilboa & Marlatte, 2017). Knowledge structures that specify the sequence of expected actions in an event are called scripts (Abelson, 1981; Abbott, Black, & Smith, 1985; Hudson et al., 1992). Such scripts may guide comprehension whenever events are experienced or referred to from memory. For example, in a restaurant script, there is a sequence of constant activities that are generally involved: being seated, getting a menu, ordering food, eating, and paying (Hudson et al., 1992). The neural substrates of such scripts might be found in the temporoparietal junction and medial prefrontal cortex. Neuroimaging literature is consistent with the role of a temporoparietal junction mirror system for inferring temporary goals and intentions at a perceptual level of representation. In contrast, the medial prefrontal cortex is related to inferring more stable characteristics of others and the self or interpersonal norms and scripts at a more abstract cognitive level (van Overwalle, 2009). Recent neuroimaging studies have noted networks of areas with medial prefrontal or hippocampal hubs for encoding or retrieval of schematic or specific aspects of event memories (e.g., Bonasia et al., 2018; Massis-Obando, Norman, & Baldassano, 2022). However, to the best of our knowledge no imaging or lesion studies have directly examined whether there are specific brain substrates of the action components of events (as opposed to overall memory including objects, participants, locations, etc.). As described below, the current study aims to address this lacuna.

Beyond using no more than two memory features simultaneously, studies of memory features generally use non-ecological paradigms to study each feature separately; little work has been conducted to understand the infrastructure of representation and retrieval of memory features in more natural visual scenes. Such scenes are composed of numerous spatially congruent objects to produce a semantically coherent view of a real-world environment (Henderson & Hollingworth, 1999). Investigating memory for such complex stimulus arrays may enable not only decomposing the memory into more basic elements and comparing them in an ecologically valid method but also better understanding the degree of overlap or independence of the neural substrates of the component aspects.

One such study by Burgess and colleagues (2001) used event-related fMRI to explore the neural substrates associated with retrieval of visual memory of rich environments. During the encoding phase, participants explored a virtual reality town and received various objects from different people in different places. They were instructed to remember each object and its context. Participants were tested for their memory for the objects, persons, and locations. Results indicated activation in bilateral anterior prefrontal cortices during retrieval of both types of context information (the person giving the item and the location of the receipt). In addition, they reported a network of regions that included the right posterior parietal lobe, precuneus, and bilateral parahippocampal gyri, specifically related to location memory.

Two studies which investigated different categories of episodic memory in an autobiographical memory context are also worth noting. An fMRI study by Gilmore et al. (2021) demonstrated separable systems in the memory network supporting people, places, and objects. The people system included the medial prefrontal cortex, posterior cingulate cortex, anterior temporal lobes, and posterior superior temporal sulcus/angular gyrus; the place system included the parahippocampal cortex, posterior angular gyrus, and retrosplenial cortex/parieto-occipital sulcus; and the objects system included the lateral occipital complex, dorsal parietal cortex, and somatosensory cortex. A study by Memel, Wank, Ryan & Grilli (2020) investigated the relationship between autobiographical retrievals of people/objects (event) details and spatiotemporal details and integrity of the anterotemporal (uncinate fasciculus) and posteromedial (cingulum bundle) cortical pathways, using DTI. They found that the anterotemporal pathway was strongly implicated in memory for event element details, while the posteromedial pathway was similarly implicated in both event element and spatiotemporal details.

As opposed to functional imaging studies, which can only show correlation between regional brain activity and performance of a behavioral task, neuropsychological studies of the behavioral consequences of brain lesions enable the determination of necessity and causality in structure-function relationships, thus complementing and aiding the interpretation of regional activations in fMRI/PET studies (Siddiqi et al., 2022; Rosenbaum, Gilboa, & Moscovitch, 2014). Unfortunately, the handful of neuropsychological studies related to this issue generally included only a few patients; they did not apply quantitative voxel-wise analyses that provide high spatial resolution; and furthermore, they were limited by a hypothesis-driven focus on predefined regions-of-interest (ROIs), leaving the involvement of other potentially relevant brain regions unassessed (Pigott & Milner, 1993; Holdstock et al., 2002, 2005; Crane & Milner 2005; Stepankova et al., 2004). Furthermore, to the best of our knowledge, with the exception of Memel et al. (2020), no studies have attempted to assess the contribution of white matter integrity to mnemonic performance for the various aspects of episodic experience elements. Hence, systematic data-driven group analyses of lesion effects on multiple aspects of visual memory with high spatial precision are warranted.

In the current neuropsychological study, we aimed to examine the causal impact of stroke lesion topography of grey and white matter structures of the brain on episodic memory for complex visual scenes. We examine several aspects of visual memory concurrently within the same study; focusing specifically on identity, location, and action memory, as assessed by the WMS-III Family Pictures subtest. In this task, participants view pictures presenting different episodes in family life. The pictures include persons, each in a certain location, performing a specific action. Participants are tested for their memory of each aspect, immediately after encoding and again after a 30-minute delay. We decomposed the standard single neuropsychological scoring of picture memory recall into separate factors of endorsement of each memory aspect. Furthermore, the present study aimed to differentiate the brain substrates of episodic memory processes that are distinct for each type of information from those supporting non-specific memory processes, using systematic group analyses of lesion effects by voxel-based lesion-behavior mapping (VLBM; Bates et al., 2003). Moreover, we aimed to determine the degree to which visual-memory-related lesions overlap regions that have been associated with visual perception of the same scene aspects. From a clinical perspective, the current study aims to assess more process-specific aspects of episodic visual memory, to improve the clinical utility of neuropsychological diagnostics and help characterize specific difficulties, as stroke patients with different damage patterns may experience difficulty remembering various types of information.

## 2. Methods

### 2.1 Participants

Ninety-three patients with first-incident ischemic or hemorrhagic hemispheric stroke in the subacute phase were recruited for the study during their hospitalization at the Loewenstein Rehabilitation Medical Center (LRMC), Raanana, Israel, or under treatment in the neurology department at Wolfson Medical Center (WMC), Holon, Israel. In the right-hemisphere-damage group (RHD; n = 54), the mean age was 60.1 years (*SD* = 12.6), 19 patients were females, eight patients were left-handed, and the group’s mean educational level was 13.4 years of formal schooling (*SD* = 3.2). In the left-hemisphere-damage group (LHD; n = 39), the mean age was 59.2 years (*SD* = 13.8), 17 patients were females, two were left-handed, and the group’s mean educational level was 12.7 years of formal schooling (*SD* = 2.8). All the patients provided informed consent to participate in the study, which was performed according to a protocol approved by the human subjects’ research committees of the LRMC and WMC, following the ethical standards prescribed in the 1964 Declaration of Helsinki. Patients were included in the study only if they did not suffer from psychiatric or prior neurological disorders, had not received psychotropic drugs regularly prior to the precipitating stroke incident, were in a stable clinical and metabolic state, had unilateral hemispheric damage, and their language and cognitive status enabled full comprehension of the task requirements. Table S1 (see Supplementary Materials) details each patient’s demographic and clinical data.

Seventy-three healthy individuals of an age range of 21-84 years, with no history of neurological or psychiatric disorders, served as comparison group participants in return for payment. Their mean age was 57.9 years (*SD* = 16.9), and they had a mean educational level of 14.0 years (*SD* = 2.8); 33 were females; 6 were left-handed; handedness data was missing for 8 participants. Ten age-matched healthy participants were selected for each stroke patient to construct individually matched comparison groups of comparable age.

### 2.2 Test procedure

The Family Pictures subtest, a recall memory test for complex pictures, was administered as part of the Wechsler Memory Scale-III (WMS-III; Wechsler, 1997) according to the standard procedures. Briefly, four scene pictures are presented to the participants for 10 seconds each, with participants instructed to remember each picture. Immediately after all scenes are presented, participants are asked to serially recall the characters, their locations, and their activities for each scene, and then asked to do so again after a 30-minute delay during which other tests are administered. Some patients (3 LHD, 5 RHD) did not perform the delayed part of the test due to technical reasons. In those cases, only the immediate test phase was analyzed.

### 2.3 Scoring method

The traditional scoring method of the WMS-III separately scores correct identifications of characters, locations, and actions; However, it then yields a single performance score, summing up all these sub-measures. To examine those different aspects of memory separately, we summed the scores of each sub-measure individually and used it as an independent measure for visual memory of character identity, location, and action, respectively. Identity memory was scored according to the WMS-III record form and scoring manual. However, in the current study, location memory was scored as correct only if either identity or action of the character were correctly recalled, but action was scored independently of the correctness of the other parameters. In the original scoring method, correct location and action memory are scored as correct only when identity is correctly recalled. The change in location and action scoring method aimed to assess those parameters as independently as possible from the other measures. However, since location memory is chosen out of four possible alternatives, and there are only four persons in a picture, generally occupying all quadrants, merely guessing that each of the test pictures included events in all four quadrants would yield high scores in the absence of any veridical memory. Hence, under the new scoring method we implemented, correctness of location memory requires indication of what person and/or action occurred in that location – i.e., the location of a meaningful element. This scoring method was not necessary for action or identity memory, as not all range of possibilities for those measures appeared in each picture. The range of possible actors for the identity measure is constrained to seven characters, but only four appear in each picture possibilities. For the action measure, possibilities are partially constrained by the script presented in the scene; e.g., participants asked to remember the picture of the family dinner are unlikely to consider activities unrelated to that context, but have several options of possible appropriate actions that might have been presented.

### 2.4 Lesion analysis

Patients’ brain damage was assessed using follow-up CT/MRI scans performed during the rehabilitation period, dating on average 41 days post-stroke onset (*SD* = 33). In some cases, where the boundaries of the damaged area were clearly defined in a scan performed at an earlier stage, lesion analysis was based on the early scan (those latencies are included in the post-onset average time provided). Lesion analyses were performed with the Analysis of Brain Lesions (ABLe) module implemented in MEDx software (Medical Numerics, Sterling, VA, USA). ABLe characterizes brain lesions in MRI and CT scans of the adult human brain by spatially normalizing the lesioned brain into MNI space. It reports anatomical structures in the normalized brain by using an interface to the Automated Anatomical Labeling (AAL) atlas, and the White Matter (WM) atlas (Lancaster et al., 2000; Solomon, Raymont, Braun, Butman, & Grafman, 2007; Tzourio-Mazoyer et al., 2002). The method we used for lesion analysis was described in detail in Haramati, Soroker, Dudai, & Levy (2008). Lesions were manually outlined on the digitized CT/MRI scans using the MEDx software and were independently confirmed and adjusted as required by a physician experienced in the analysis of neuroimaging data (author NS).

Registration accuracy of the scans to the MNI template (calculated by MEDx software using the ABLe module implemented in it) across subjects ranged from 89% to 95.6% (94.2 ± 1.2, 94.1 ± 1.1 in RHD and LHD subjects, respectively; registration accuracy information from 7 RHD patients and 5 LHD patients was not extant due to technical problems). Table S2 shows the percent of damage for each brain region/structure in each patient. In addition, a comparison of lesion distribution was made between the RHD and LHD groups concerning (1) the proportion of subjects affected in each brain region/structure of the AAL and WM atlases (Lancaster et al., 2000; Solomon, Raymont, Braun, Butman, & Grafman, 2007; Tzourio-Mazoyer et al., 2002; Mori et al., 2008), using Fisher’s exact test, and (2) the extent of damage in each region, using the Mann-Whitney test, applying the Holm-Bonferroni correction method for multiple comparisons.

### 2.5 Voxel-based lesion-behavior mapping (VLBM)

VLBM (also termed VLSM - voxel-based lesion-symptom mapping; Bates et al., 2003) analyzes structure-function relationship in the brain in a voxel-by-voxel manner. For each voxel, ABLe determines which patients have and which lack damage to that voxel (lesions having been normalized to a common space prior to this analysis). Then, a t-test is computed between the behavioral results of these two groups of patients (with potentially different patients for each voxel). The output of this procedure is a map where each voxel is assigned a t-value. Each hemispheric group (RHD, LHD) was analyzed separately. A voxel was included in the analysis if at least 10% of the subjects in the relevant hemispheric group had damage to that voxel. To further correct for multiple comparisons, only voxels implicated in deficits with values exceeding a false discovery rate (FDR) threshold (Benjamini & Hochberg, 1995) of *p* < .05 were considered significant (Genovese, Lazar, & Nichols, 2002; Frenkel-Toledo et al., 2019). In addition, at least twenty-five adjacent voxels had to show a statistically significant impact on performance for a cluster of voxels to be reported (for a similar method see Ben-Zvi, Soroker & Levy, 2015).

Additionally, cases in which the results passed permutation correction (Nichols & Holmes, 2002) are indicated. Similar to FDR, permutation correction for multiple comparisons aims to reduce Type I error; however, while FDR uses adjusted *p*-value threshold levels, the permutation method re-samples the total number of observations to calculate all possible rearrangements of the data and compute an estimated null distribution, from which the *p*-value is calculated.

Taking into account that insufficient statistical power might have characterized some of the analyses, we also report anatomical regions associated with deficits that did not survive FDR correction for multiple comparisons but were implicated using a more lenient criterion of p < .005, reflecting z > 2.6 (for a similar approach see Wu et al., 2015; Schoch, Dimitrova, Gizewski, & Timmann, 2006; Moon, Pyun, Tae, & Kwon, 2016; Lo, Gitelman, Levy, Hulvershorn, & Parrish, 2010; Ben-Zvi Feldman, Soroker, & Levy, 2021). This information is provided under the assumption that in such cases, VLBM indicates areas likely implicated in the relevant mnemonic process and which require further investigation. The maximum z-score reflecting differences between the performance of patients with damage to the voxels in question vs. those without damage to those voxels is reported for each cluster of contiguous above-threshold voxels. Since there may be multiple voxels with this maximum z-score in the cluster, we report the coordinate of the voxel that is most superior, posterior, and left in its location within the cluster (the centroid of the cluster is not reported, as it may not have the highest z-score value and it may not be an above-threshold voxel). The AAL atlas for gray matter and the White Matter Atlas (Lancaster et al., 2000; Solomon et al., 2007; Tzourio-Mazoyer et al., 2002; Mori et al., 2008) were used to identify the locations of the significant clusters. Conjunction analysis was used to characterize voxels surpassing the VLBM thresholds for identity vs. location vs. action on both the immediate and delayed testing phases by overlaying significant voxels from each analysis on the same template. Since these comparisons were made between regions implicated in behavioral measures that survived different criteria, we used the lenient criterion (*p* < .005 reflecting *z* > 2.6).

## 3. Results

### 3.1 Memory test performance

Data were analyzed separately for RHD and LHD patients and compared with healthy participants, both on a group basis and for each patient relative to her/his individually matched comparison group. Group average percent correct raw scores and average z-scores for identity, location, and action memory are presented in Table 1.

**Table 1.** Identity, location, and action memory on immediate and delayed test phases in RHD and LHD patient groups and the control group.

| Test Phase | Group | identity raw | identity Z-score | location raw | location Z-score | action raw | action Z-score |
| --- | --- | --- | --- | --- | --- | --- | --- |
| Immediate | RHD (n=54) | 0.69 (0.20) | -0.83 (1.65) | 0.50 (0.27) | -0.68 (1.30) | 0.37 (0.24) | -0.66 (1.54) |
|  | LHD (n=39) | 0.70 (0.19) | -0.79 (1.39) | 0.47 (0.27) | -0.87 (1.37) | 0.32 (0.23) | -1.13 (1.44) |
|  | Control (n=73) | 0.80 (0.16) | - | 0.64 (0.23) | - | 0.49 (0.19) | - |
| Delayed | RHD (n=49) | 0.68 (0.22) | -0.83 (2.10) | 0.46 (0.30) | -0.57 (1.28) | 0.35 (0.27) | -0.55 (1.38) |
|  | LHD (n=36) | 0.69 (0.21) | -0.80 (1.63) | 0.52 (0.30) | -0.54 (1.50) | 0.34 (0.23) | -0.75 (1.33) |
|  | Control (n=73) | 0.77 (0.19) | - | 0.62 (0.26) | - | 0.46 (0.20) | - |

Raw and Z-score matched identity, location, and action recall memory measures in the family pictures subtest of the WMS-III (see Methods section for explanation); SDs appear in parentheses. RHD/LHD = right/left hemisphere damage.

We examined differences in memory performance both between the RHD and LHD groups and between the experimental groups and comparison group, for each information types in immediate and delayed testing phases (Fig. 1) by conducting a mixed-model repeated-measures ANOVA on participants’ raw scores, with test phase (immediate, delayed) and stimulus type (identity of actors, location, action) as within-subjects factors and group (RHD, LHD, healthy participants) as between-subjects factor. The *p* values for this and all subsequent analyses are reported after Greenhouse-Geisser correction when required. Raw scores analysis disclosed significant main effects of group, *F*(2,155) = 6.47, *p* < .005. To evaluate which groups were significantly different, Bonferroni post-hoc comparisons were conducted. The results indicated that the control group performed significantly higher than the LHD and RHD groups (*p* = .018 and *p* = .006, respectively). The performance of the LHD and RHD groups did not differ significantly (*p* = 1.0). In addition, a significant main effect of test phase was found, *F*(1,155) = 6.10, *p* < .025, due to lower performance in the delayed test phase compared to the immediate test phase; and of stimulus type, *F*(1.90,293.96) = 372.38, *p* < .001, stemming from significant difference between each stimulus type and the two other stimulus types, with highest performance for identity, medium for location and lowest for action. The test phase x group interaction was not significant, *F*(2,155) = 2.02, *p* = .14, nor were the stimulus type x group interaction, *F*(4,310) = 1.53, *p* = .20, or the test phase x stimulus type x group interaction, *F*(4,310) = 1.52, *p* = .20. Overall scores for each group (RHD, LHD, control), for each test phase (immediate, delayed), and stimulus type (identity, location, action) are shown in Table 2.

**Table 2.** Overall score of WMS III Family Pictures subtest for each group (RHD, LHD, healthy comparison group), test phase (immediate, delayed), and stimulus type (identity, location, action) factors.

| Factor | Group | Overall score |
| --- | --- | --- |
| Group | RHD | 51.12<br>(23.16) |
|  | LHD | 51.37<br>(22.0) |
|  | Healthy | 63.02<br>(17.79) |
| Test phase | Immediate | 56.83<br>(20.83) |
|  | Delayed | 55.70<br>(22.43) |
| Stimulus type | Identity | 72.97 |
|  | Location | 54.59<br>(26.34) |
|  | Action | 40.16<br>(22.75) |

**Figure 1.**
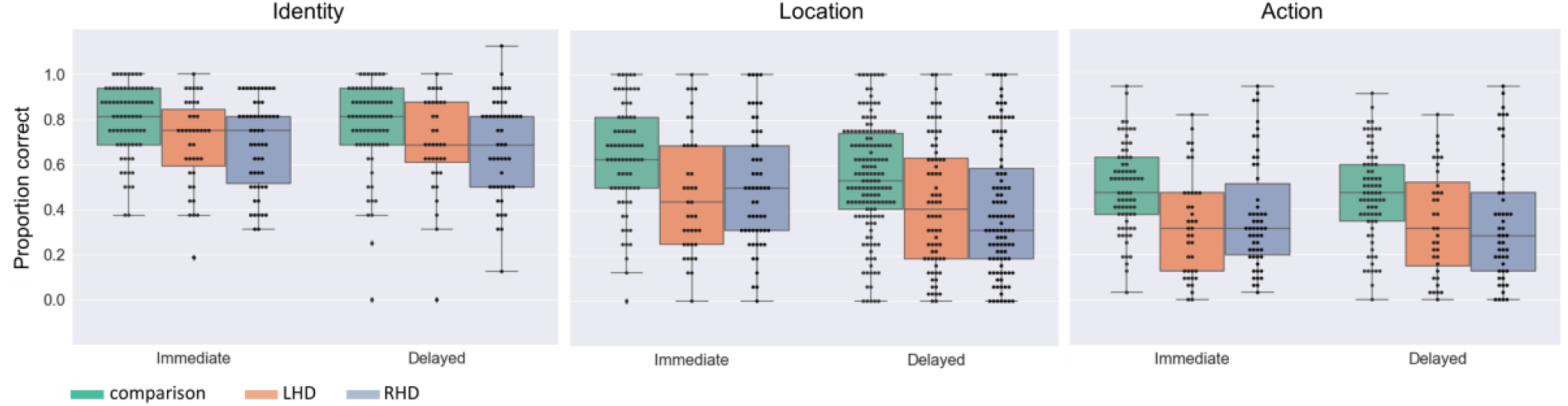
Immediate and delayed recall memory for memoranda identity, location, and action. Raw scores of healthy comparison group (n=73), LHD (n=39), and RHD (n=54) participants for identity, location, and action memory in the immediate and delayed test phases. For each group, a boxplot with individual data points illustrates the distribution of that group’s raw scores overall participants. The box marks the first and third quartiles, the continuous line marks the median, and the whiskers represent 1.5 times the interquartile range (i.e., the range between the first and third quartiles) above the third quartile and below the first quartile.

As a result of the heterogeneity in patients’ age, which can affect memory performance, for the lesion-behavior analysis, we used the more individually focused age-matched Z scores as the behavioral measure for each patient (for a similar method, see Ben-Zvi Feldman et al., 2021; Ben-Zvi et al., 2015).

### 3.2 Lesion analysis

Table S2 shows the extent of damage to each region of the AAL and WM atlases for each patient. In the following brain regions, the percent of subjects who had at least 1% of the region damaged by the stroke was larger in the RHD compared to the LHD patient group: superior and polar regions of the temporal lobe; orbital, inferior, and middle frontal gyri; supramarginal and angular gyri; insula, precentral and postcentral gyri, Heschl gyrus, the Rolandic operculum and the superior longitudinal fasciculus.

Additionally, the extent of damage was significantly larger in the RHD compared to the LHD patient group in the following regions: middle, superior and polar regions of the temporal lobe; orbital, inferior, and middle frontal gyri; supramarginal and angular gyri; insula, precentral and postcentral gyri, Heschl gyrus, the Rolandic operculum and white matter fibers of the superior longitudinal fasciculus, external capsule and superior corona radiata (detailed descriptive statistics are presented in Table S3). The above group differences could affect VLBM results, therefore hemispheric differences in lesion effects of homologous regions should be interpreted with caution.

Hemispheric volume loss ranged from 0.39 to 278 cm^3^ (*M* = 75 cm^3^, *SD* = 63 cm^3^) in the RHD group, and from 0.41 to 123.19 cm^3^ (*M* = 28 cm3, *SD* = 29 cm^3^) in the LHD group. Lesion size information from 7 RHD patients and one LHD patient was not extant due to subsequent data loss (however, those data were available when the functional analyses reported below were conducted). Figure 2 shows the overlap of lesions in the RHD and LHD patient groups. As can be seen in this figure, most patients in both groups had lesions within the middle cerebral artery (MCA) territory, as is usually encountered in a cohort of stroke patients. Inferences from the current lesion analysis are confined to brain voxels affected in a sufficient number of subjects, as explained in the Methods section.

**Figure 2:**
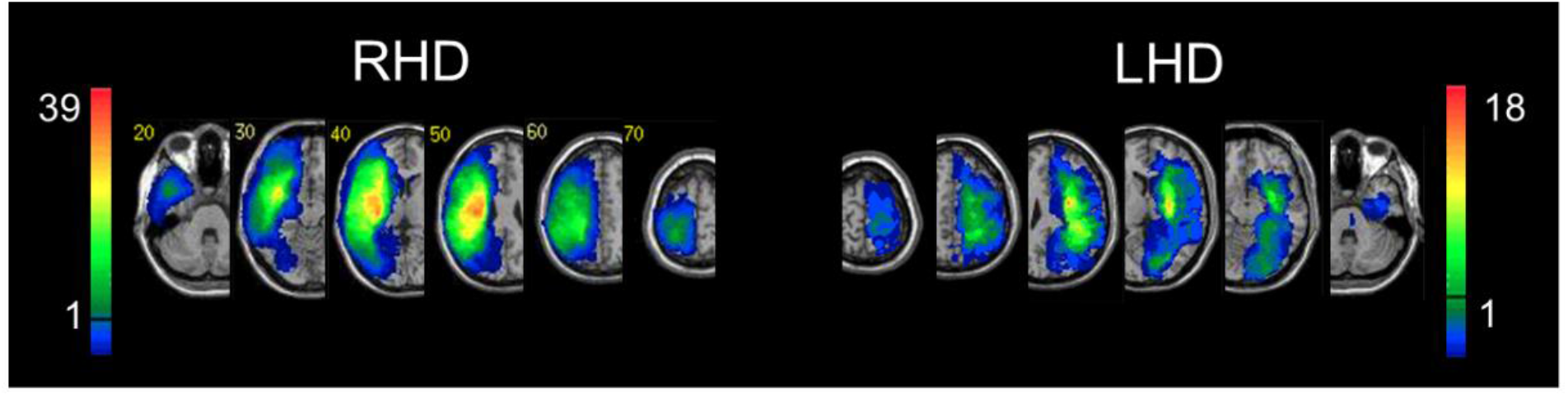
Lesion overlay. Representative normalized slices from 54 RHD patients (left) and 39 LHD patients (right). A voxel was included in the VLBM analysis if damaged in at least 10% of the cohort (5 and 4 patients in RHD and LHD groups, respectively). This threshold is denoted by the horizontal bar in the color code column representing the number of patients damaged in each region of the brain. Corner numbers indicate normalized slice numbers out of 90 total slices. The left side of the image corresponds to the right side of the brain (i.e., neurological convention).

### 3.3 Voxel-based lesion-behavior mapping (VLBM)

VLBM analysis identified clusters of voxels in which damage was associated with impaired performance. As explained in the Methods section, patients’ performance was expressed as Z-scores computed for each patient in reference to 10 age-matched healthy controls. VLBM analysis was done in 54 RHD patients in the immediate testing phase and 49 in the delayed testing phase. In the LHD group, VLBM analysis was done in 39 patients in the immediate testing phase and 36 patients in the delayed testing phase. The lists of brain regions containing clusters of 25 contiguous ‘significant’ voxels, or more are presented in Tables S4-9, which depict the VLBM results corresponding to the identity, location, and action recall memory, in the RHD and LHD groups.

Memory of actor identity in RHD patients (Figure 3; Table S4) was affected in the immediate testing phase by damage to voxel clusters in the superior and middle temporal gyri (STG, MTG), supramarginal and angular gyri (SMG, ANG) and the hippocampus, as well as in the following white matter tracts: superior longitudinal fasciculus (SLF), posterior thalamic radiation (TR_P_), posterior part of the corona radiata (CR_P_), sagittal stratum (SS) and retrolenticular limb of the internal capsule (IC_RL_). The delayed testing phase showed smaller clusters in the same regions with the exception of IC_RL_, SS, TR_P_, SMG and SLF, where damage was related to poorer performance in the immediate but not in the delayed testing phase. Clusters were significant at a threshold of *z* > 2.6 (*p* < .005). It is possible that some areas implicated in immediate test performance decrement did not emerge as being involved in delayed testing decrement because five fewer RHD patients completed the delayed tests; those regions were found to be related to identity memory in the VLSM analysis for the delayed testing phase using a lower threshold of z > 2.3 (*p* < 0.1).

**Figure 3.**
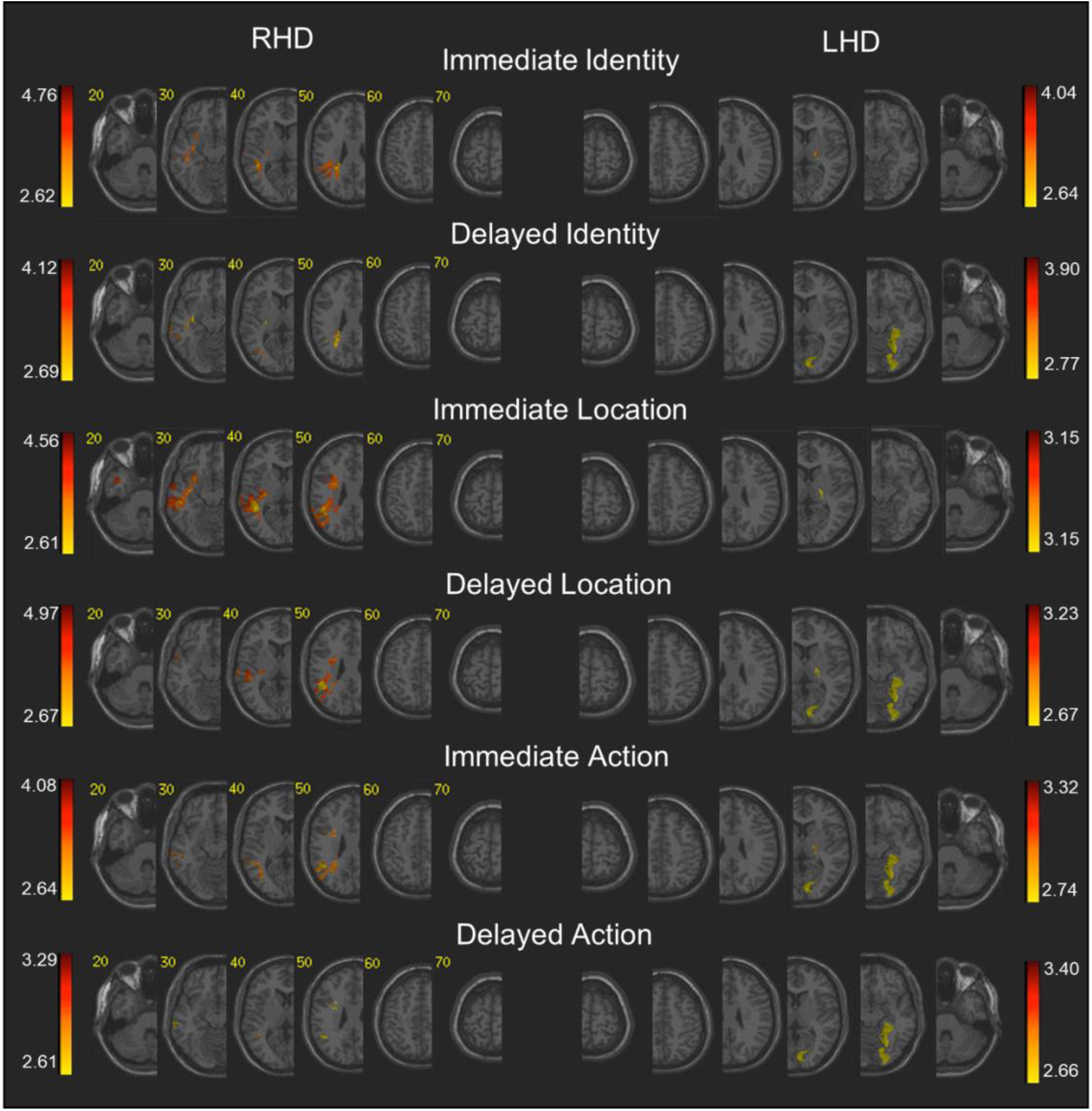
Voxel-based lesion-behavior mapping (VLBM) analysis in the RHD and LHD groups. Colored regions denote brain voxels in which the existence of damage significantly impacted the following measures of immediate and delayed memory for an object’s identity, location, and action for the RHD (left side) and LHD (right side) patients’ groups. The colored voxels passed threshold of z ≥ 2.6, (*p*≤ 0.005). Voxels were analyzed only if they were damaged in at least 10% of the patients. Regions in red correspond to higher z scores. Corner numbers indicate normalized slice numbers out of 90 total slices. The left side of the image corresponds to the right side of the brain (i.e., neurological convention).

Memory of actor identity in LHD patients (Figure 3; Table S7) was affected in the immediate testing phase by damage only to a small cluster in the anterior limb of the internal capsule (IC_AL_), which was significant at a threshold of *z* > 2.6 (*p* < .005). In contrast, actor identity memory in the delayed testing phase was affected in LHD patients by damage to occipito-temporal regions encompassing the primary visual cortex (calcarine region) and adjacent lingual, fusiform, parahippocampal and inferior occipital regions. Significant voxel clusters in these regions survived FDR correction for multiple comparisons.

Location memory for meaningful scene elements was affected in RHD patients (Figure 3; Table S5) in the immediate testing phase by damage to voxel clusters in temporo-parietal, posterior frontal, opercular, and middle occipital cortical regions, as well as by damage to medial temporal-lobe regions (hippocampus, amygdala), the basal ganglia and white matter association and projection tracts (see detailed list of significant anatomical structures in Table S5). Part of these areas were also involved in the mediation of actor identity memory; however, there was a greater extent of areas implicated in location memory in the temporal cortex (including also temporo-polar region and Heschl and inferior temporal gyri), larger clusters in inferior parietal regions, with the addition of areas not implicated in identity memory - somatosensory cortex, inferior frontal, insular and opercular cortical regions, the amygdala, putamen and pallidum, and some association and projection white matter tracts, including the uncinate fasciculus, posterior limb of the internal capsule (IC_PL_), anterior corona radiata (CR_A_), and the inferior portion of the fronto-occipital fasciculus (FO_I_), that were not involved in the network mediating identity memory. In most of these regions, damage affected location memory only in the immediate testing phase. As mentioned above regarding actor identity memory, it is possible that some areas implicated in immediate test performance decrement did not emerge as being involved in delayed testing decrement because five fewer RHD patients completed the delayed tests; some of the above-mentioned regions were found to be related to location memory in the VLSM analysis for the delayed testing phase using a lower threshold of z > 2.3 (*p* < 0.1).

These effects survived the FDR correction for multiple comparisons. The effects of damage to the superior temporal gyrus on the immediate and delayed testing phases also survived permutation correction. The magnitude of lesion effects observed for location memory presumably also result from the characteristics of the location measure used in the current study, which depends on the identity and/or action knowledge together with location information.

Location of meaningful element memory was affected in the LHD group (Figure 3; Table S8) in a manner quite similar to identity memory, with ‘significant’ voxel clusters in occipito-temporal regions encompassing the primary visual cortex (calcarine region) and adjacent lingual, fusiform, parahippocampal, hippocampal and inferior occipital regions, all showing significant effect of damage on performance in the delayed but not in the immediate testing phase, which survived FDR correction for multiple comparison.

Action memory in the RHD group (Figure 3; Table S6) was affected quite similarly to identity and location memory, by damage to voxel clusters mainly in a temporo-parietal network of cortical structures, with the addition of damage to association and projection white matter tracts and small areas of damage within the inferior frontal gyrus and the middle occipital gyrus. Lesion effects were more robust in the immediate than in the delayed testing phase for action memory. Clusters were significant at a threshold of *z* > 2.6 (*p* < .005).

Action memory in the LHD group (Figure 3; Table S9) was affected, both in the immediate and delayed testing phases, by damage to large voxel clusters in occipito-temporal regions encompassing the primary visual cortex (calcarine region) and adjacent lingual, fusiform, parahippocampal and inferior occipital regions, i.e., quite similarly to the anatomical pattern of damage revealed in VLBM of identity and location memory in the delayed testing phase. The insula and the anterior portion of the corona radiata were implicated in action memory but not in identity and location memory in this group. These effects survived the FDR correction for multiple comparisons.

### 3.4 VLBM conjunction analysis

Figure 4 and Table S10 show the anatomical structures where VLBM conjunction analysis revealed voxel clusters on the right and left hemispheres in which damage exerts a significantly specific impact on identity vs. location vs. action memory (or alternatively, an impact on all types) in the immediate and delayed testing phases. All anatomical comparisons are presented using the lower threshold of z ≥ 2.6 (*p* ≤ .005) to accommodate measures that passed different threshold levels.

**Figure 4.**
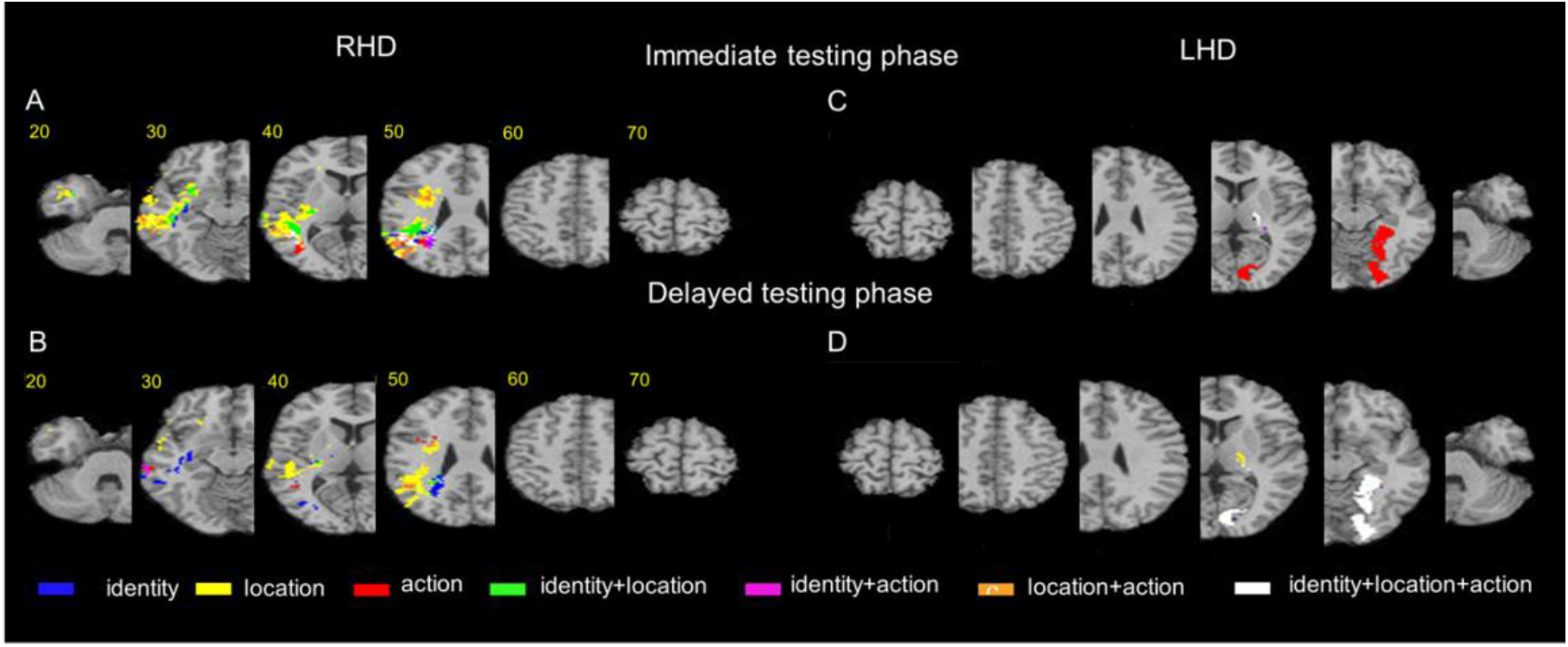
Voxel-based lesion-behavior mapping (VLBM) conjunction analysis. Conjunction VLBM analysis comparing: (A) (top left panel) identity vs. location vs. action memory measures on the immediate testing phase for the RHD group; (B) (lower left panel) the delayed testing phase for the RHD group; (C) (top right panel) the immediate testing phase for the LHD group; (D) (lower right panel) the delayed testing phase for the LHD group. The colored voxels passed the threshold of z > 2.6, (p < 0.005). Representative slices from VLBM maps were computed for each behavioral measure and overlaid on an MNI template brain. Colored pixels represent voxels in which the existence of damage exerted a significant impact on the tested behavior. Displays follow radiological convention, i.e., right hemisphere displayed on the left side. For a verbal description of the brain regions in which significant voxel clusters have been found in each conjunction analysis, see Table 10.

Conjunction VLBM analysis comparing identity vs. location vs. action memory measures on the immediate testing phase in the RHD group (Figure 4, top left panel; Table S10-A) reveals non-specific effects on all stimulus types by damage within the middle and superior temporal gyri, the angular gyrus, and the superior longitudinal fasciculus. However, damage to other portions of the same anatomical regions were found to affect each of the three stimulus types in a specific manner. Damage to other cortical and subcortical regions of the right hemisphere was found to present mostly specific effects. Specificity for identity memory was found in the immediate testing phase for voxel clusters within the hippocampus, middle temporal gyrus and the corona radiata. Specificity for location memory was found in the immediate testing phase for large voxel clusters within the lateral temporal cortex and smaller clusters in temporo-polar, insular, inferior parietal and middle occipital cortices, the putamen and a series of white matter projection and association tracts. VLBM conjunction analysis in the RHD group did not reveal specificity for action memory in the immediate testing phase (i.e., the effect of voxels’ damage restricted to action memory only). Damage to several voxel clusters, notably in temporo-parietal regions and some white matter tracts (detailed in Table S10-A) was found to affect two types of stimuli (i.e., identity plus location, identity plus action, or location plus action).

Conjunction VLBM analysis comparing identity vs. location vs. action memory measures on the delayed testing phase in the RHD group (Figure 4, top left panel; Table S10-A) did not reveal non-specific effects (i.e., damage to voxel clusters affecting all three stimulus types). Voxels’ damage specifically affecting identity memory at the delayed testing phase was found within the middle and superior temporal gyri, the hippocampus and a few white matter tracts. Damage specifically affecting location memory at the delayed testing phase was found in the RHD group in large voxel clusters within the superior, polar and middle temporal cortex, and in smaller clusters within the angular and supramarginal gyri, the superior longitudinal fasciculus, and the corona radiata. In small voxel clusters within the right middle temporal gyrus and inferior frontal gyrus, the existence of damage had a specific effect on action memory only. Damage to several, relatively small, voxel clusters in temporo-parietal regions was found to affect two types of stimuli (i.e., identity plus location, identity plus action or location plus action) in the delayed testing phase.

Conjunction VLBM analysis comparing identity vs. location vs. action memory measures on the immediate testing phase in the LHD group (Figure 4, top right panel; Table S10-B) did not reveal non-specific effects (i.e., effects on all the three stimulus types), and also did not reveal specific effects on location memory. Damage to small voxel clusters within the anterior and posterior limbs of the internal capsule had a specific effect on identity memory only. Damage to large occipito-temporal clusters in the lingual, fusiform, calcarine, parahippocampal and inferior occipital gyri, together with portions of the insula and anterior corona radiata, specifically affected action memory.

Conjunction VLBM analysis comparing identity vs. location vs. action memory measure on the delayed testing phase in the LHD group (Figure 4, lower right panel; Table S10-B) revealed non-specific effects on all the three stimulus types for damage to large voxel clusters within the temporo-occipital cortex, in the lingual, fusiform, calcarine, parahippocampal and inferior occipital gyri. Damage to a small voxel cluster within the anterior limb of the internal capsule had a specific effect on identity memory only, and damage to a small voxel cluster within the anterior limb of the internal capsule had a specific effect on action memory only. There were no specific effects on location memory in the LHD group in the delayed testing phase.

## 4. Discussion

The purpose of the present study was to investigate, using lesion-behavior mapping in patients after stroke, the shared and differential anatomical substrates of memory for identity, location and action of persons in a scene, which are key features of visual episodic memory. We identified right and left-hemisphere anatomical loci of cognitive processes underlying visual episodic memory recall. We found that both RHD and LHD patients’ performance was lower than that of healthy participants on all measures (as documented in Table 1). However, we found differences between patterns of lesions that affect those measures. Our lesion-behavior analyses (Figures 3-4 and Tables S4-10) revealed non-selective effects on all three dimensions (actor identity, location and action) associated with damage to large voxel clusters in the left occipitotemporal cortex in the delayed testing phase and in the right lateral temporoparietal cortex in the immediate testing phase. In contradistinction, damage to other clusters in this analysis was specifically implicated in only one or two of the measures. In the following sections, we will discuss possible interpretations of this pattern of lesion-behavior mapping.

### 4.1 Remembering actor identity

Actor identity memory was most prominently affected, in the immediate testing phase, by damage to the temporo-parietal cortex (superior and middle temporal gyri, angular and supramarginal gyri of the parietal lobe) and the superior longitudinal fasciculus, as shown in the RHD group lesion-behavior analysis. In the delayed testing phase, it was most prominently affected by lesions in occipitotemporal regions (calcarine region, lingual, fusiform, and inferior occipital gyri), as shown in the LHD group lesion-behavior analysis. However, those regions were mostly non-specific to identity processing.

Memory for characters’ identity was also associated with integrity of identity-specific clusters in the right hippocampus.

### 4.2 Remembering the locations of meaningful scene elements

Like identity memory, location memory was prominently affected by lesions in the temporoparietal regions and SLF in the immediate testing phase, as shown in the RHD group lesion-behavior analysis. Location memory in the delayed testing phase was prominently and non-specifically affected by left occipitotemporal lesions. Right temporoparietal regions and SLF were found to contain very high-specificity clusters related only to location memory, and less specific clusters related also to other memory features.

In accord with this high specificity of location-memory clusters, a TMS study by Chambers and colleagues in healthy participants demonstrated that AnG has a crucial role in orienting of spatial attention (Chambers, Payne, Stokes, & Mattingley, 2004). It is further notable that lesion studies have shown that posterior parietal damage impaired object location memory performance (Kessels et al., 2000; van Asselen et al., 2009).

The right temporal pole was implicated in the current VLBM analysis as location-memory specific. In accordance with this finding, the temporal pole’s role in spatial processing was demonstrated in a case of a patient with a right temporal pole focal lesion who failed in tasks requiring recall of spatial position or spatially based imagery operations (Luzzatti, Vecchi, Agazzi, Cesa-Bianchi & Vergani, 1998). with a study Right insula specificity corresponds with a lesion study showing evidence for the involvement of the right insula in spatial attention processes (Marin et al., 2017). A meta-analysis of neuroimaging studies on spatial cognition by Cona and Scarpazza (2019) has identified a core network of spatial processing, including the anterior insula.

The current VLBM results identified a location-related voxel cluster in the right hippocampus, an area the integrity of which was also identified with identity memory. This is in accordance with the notion that the hippocampus is the binding site of object location and identity information, creating unified representations (Gaffan, 1998; Manns & Eichenbaum, 2006; Knierim, Lee, & Hargreaves, 2006; Diana, Yonelinas & Ranganath, 2007). In the current paradigm, location memory is not an independent function, rather requiring identity (and action) information. It thus more greatly represents binding of location information with the other features. Lesion studies have also shown that right hippocampal lesions cause deficits in various tasks of object-location and spatial memory (Kopelman, Stanhope & Kingsley, 1997; Pigott & Milner, 1993; Smith & Milner, 1981, 1984; Holdstock et al., 2002, 2005; Crane & Milner 2005; Nunn, Graydon, Polkey & Morris, 1999; Stepankova et al., 2004; van Asselen et al., 2009). Patients with hippocampal pathology due to perinatal anoxia (Vargha-Khadem et al., 1997) show impaired object-location memory. According to two reviews of the literature of lesion studies (Postma, Kessels & van Asselen, 2008; Zimmermann & Eschen, 2017) and neuroimaging studies (Zimmermann & Eschen, 2017), the right hippocampus plays a critical role in object-location processing.

The right middle occipital gyrus is also implicated as a location-specific region in the current VLBM analysis. This is in accordance with fMRI activations found in the contralateral middle occipital gyrus during a spatial attention task (Mangun, Buonocore, Girelli, & Jha, 1998), and increased rCBF observed in a PET study during the processing of spatial location in the right middle occipital gyrus (Kohler et al., 1998).

Right white matter tracts IC_RL_, external capsule, SS, and inferior fronto-occipital fasciculus were also implicated in the VLBM analysis as location-specific. According to lesion study by Marin and colleagues, lesions in those regions in the right hemisphere impair visuospatial attention (Marin et al., 2017).

### 4.3 Remembering actions

As for identity and location, action memory was prominently affected by lesions in the right lateral temporoparietal regions and SLF in the immediate testing phase. However, in contrast with identity and location memory, it was also prominently affected by lesions in left occipitotemporal areas both on the immediate and delayed testing phases. Some action-related clusters were action-specific, whereas others were implicated in performance on the other memory features.

In accord with the finding of lateral temporoparietal voxel clusters with high specificity for action memory, a meta-analysis of functional imaging studies revealed action processing performed by MTG and STG (Caspers, Zilles, Laird, & Eickhoff, 2010). fMRI studies have demonstrated action processing related activations in MTG (Wurm & Lingnau, 2015; Wurm, Ariani, Greenlee, & Lingnau, 2016; Wurm, Caramazza, & Lingnau, 2017).

Moreover, right pSTS has been shown to play a crucial role in representing actions (Saxe, Xiao, Kovacs, Perrett, Kanwisher, 2004; Cattaneo & Rizzolatti, 2009; Iacoboni et al., 2005). The impact on action memory of damage to MTG and STG which we found may also reflect the contribution of the STS to action processing. Additionally, in accordance with SLF action-specific clusters, SLF lesions were associated with poor gesture comprehension (Binder et al., 2017).

The IFG pars opercularis was identified in the current VLBM analysis as being specifically important for action memory. This region is presumably a human homolog of the macaque mirror area F5, and hence, a core region of the putative human mirror neurons system, which is believed to constitute a fronto-parietal circuitry (Iacoboni et al., 1999; Rizzolatti, Fogassi & Gallese, 2001; Rizzolatti and Craighero, 2004; Fabbri-Destro & Rizzolatti, 2008; Rizzolatti & Fabbri-Destro, 2010; Rizzolatti and Sinigaglia, 2010).

The left insula was specifically related to action memory in the current VLBM results. A few functional imaging studies have demonstrated the importance of insula to action processing. In a study by Paulus and colleagues, the insula was specifically engaged during assessment and action selection (Paulus, Feinstein, Leland, & Simmons, 2005). Additionally, a lesion study by Binder and colleagues has shown that lesions within the left insula caused deficits in gesture comprehension (Binder et al., 2017).

Left CR_A_ was also implicated in the current VLBM analysis as action-specific. Previous lesion studies have shown that lesions in left CR_A_ caused impairments in action verb comprehension (Riccardi, Yourganov, Rorden, Fridriksson, & Desai, 2019) and gesture comprehension (Binder et al., 2017).

Left occipitotemporal regions have shown high specificity to action memory on the current VLBM analysis on the early testing phase, while they were non-specific on the delayed testing phase. The strong link of those regions with action processing is corroborated with a meta-analysis of fMRI studies of mirror neurons activity, showing consistent activation across action observation experiments in the extrastriate cortex (Caspers et al., 2010). Lesions in those regions may cause deficits in processes of visual recognition and perception and working memory, which are integral components of action observation.

### 4.4 Non-specific substrates

As noted, non-selective clusters on the right lateral temporoparietal cortices (AnG, STG, and MTG) were implicated by the VLBM analysis as important for all three memory functions assessed in this paradigm. Those regions are vital parts of the core recollection network (CRN), an episodic memory network that is specifically associated with recollection, according to functional imaging studies (Rugg & King, 2018; Vilberg & Rugg, 2012; Johnson & Rugg, 2007). The CRN, in accordance with the present findings, supports consciously accessible representations of prior experiences, regardless of the content of the information recollected. (Hayama, Vilberg, & Rugg, 2012; Johnson & Rugg, 2007); Rugg, & Vilberg, 2013; Vilberg, & Rugg, 2012). Voxel clusters supporting all memory features may perform essential memory functions, free of category or task segregation.

What kinds of processes may cortical CRN areas contribute to remembering? The lateral parietal cortex is the most studied among those regions, and hence, numerous suggestions regarding its memory functions were proposed, among which, attentional processes (e.g., Cabeza et al., 2008, 2011; Ciaramelli et al., 2008, 2010), expectation, and salience (Buchsbaum et al., 2011; O’Connor et al., 2010), retrieval buffering (Vilberg & Rugg, 2008, 2009a,b), or subjective report (Hower et al., 2014; Simons et al., 2010; reviewed by Levy, 2012). It has also been suggested that parietal activity reflects stimulus-specific representations (Lee et al., 2017), memory precision (Richter et al., 2016), semantic representations (Binder et al., 2009; Price et al., 2015), and multimodal integration of features into a unified episodic representation (Shimamura, 2011). However, late-retrieval roles such as feature integration were not directly tested in the current study, as no condition assesses the binding of features together as opposed to single features’ memory.

Furthermore, as noted, a large cluster of left occipitotemporal regions was associated non-specifically with all tasks in the delayed testing phase. This cluster constitutes part of the visual cortical processing stream associated with item identification, suggesting a perceptual processing role for those regions across all memory aspects. This is in accordance with the notion that episodic visual memory retrieval entails the reactivation of some of the same brain regions engaged when encoding the corresponding information, thus activating the visual cortices in the current task (Wheeler et al., 2000). However, since striate and extrastriate cortical lesions are often confounded with deficits in visual perception, it is difficult to know whether memory deficits after such lesions are due to damage of the storage representations or damage to the sites producing the percept that is to be stored.

The present study found that ventral perceptual stream regions were involved in persons’ identity, location, and action memory. This evidence seemingly contrasts with prior PET findings showing differentiation between identity and location memory related to ventral and dorsal streams, respectively (Moscovitch et al., 1995; Kohler et al., 1998). However, only contrasting between those tasks disclosed a dissociation between the streams; when contrasted separately to a baseline task, identity and location memory activity was found in both perceptual streams, demonstrating the importance of ventral and dorsal stream regions for both tasks. Furthermore, in accordance with our findings, increasing evidence shows that dorsal and ventral streams carry both ‘what’ and ‘where’ information (albeit to different degrees). There is evidence that spatial information is also processed in the ventral stream (Lehky et al., 2008; Sereno & Lehky, 2011), while object shape information has been found in the dorsal stream (Sereno & Maunsell, 1998; Murata, Gallese, Luppino, Kaseda, & Sakata, 2000; Sereno & Amador, 2006; Konen & Kastner, 2008).

The white matter tract SLF is another brain structure where damage to voxel clusters non-specifically affected all the three memory features. It is a large bundle of fibers connecting the parietal, occipital, and frontal cortices (Schmahmann, Smith, Eichler, & Filley, 2008). In accordance with our findings, DTI studies have demonstrated that aberrations in the right SLF significantly impair visual perception (Kim, Jeon & Park, 2020; Hoeft et al., 2007).

### 4.5 Laterality

The current VLBM results manifest lateralization of lesion effects shared by actor identity, location and action memory processes within the right CRN and the left ventral stream region. The left ventral stream visual regions manifested in the current VLBM results are in accordance with reported fMRI activations, mainly in the fusiform gyrus, seen during picture memory recall (Wheeler et al., 2000). Additionally, both lesion (Farah, Levine, Calvanio, 1988; Grossi, Modafferi, Pelosi, & Trojano, 1989; Thorudottir et al., 2020) and neuroimaging work (Spagna, Hajhajate, Liu, & Bartolomeo, 2021; D’Esposito et al., 1997) has suggested that left visual regions are related to mental image generation. These results suggest that left visual cortical areas are involved in retrieving visual information from long-term memory. However, since there are significant differences in lesion distributions between the RHD and LHD groups, we cannot draw any firm conclusions regarding the lack of involvement of the right occipito-temporal cortex and the left temporo-parietal cortex in visual memory recall. Differences in the involvement of lateral temporo-parietal regions between RHD and LHD groups can be seen in Tables S3A and S3B. Less involvement of left lateral temporo-parietal regions in the current study stems from exclusion of LHD patients with extensive lesions in these regions, who had severe aphasia precluding comprehension of task demands. Differences in occipito-temporal lesion distribution between the two groups are probably related to a low number of participants with posterior cerebral artery infarction in the cohort, which comprised mainly (in both groups) of patients with middle cerebral artery stroke. The different number of patients in LHD and RHD groups, created different thresholds for the number of patients damaged in these regions who could be included in the VLBM analysis. Hence, in cases of PCA infarction, damaged voxels in some regions which did not pass the threshold for inclusion in the analysis due to low representation could still affect visual memory without being detected.

### 4.6 Limitations

Beyond the current study’s inability to assess laterality effects, another limitation arising from our use of the Family Pictures subtest of the Wechsler Memory Scale as an evaluative instrument is that this task does not provide optimal location measure characteristics. As noted, this measure is defined for the purpose of the current study as memory for the location of a meaningful scene element, and hence, is not a pure measure of location memory. Furthermore, the identity, location, and action measures are not fully equivalent in the number of possible response choices. However, since our study uses ecological situations as stimuli, and participants presumably rely on their pre-existing schemas that create certain expectations for actions that can happen in the situation. The reliance on schemas makes the possible actions more limited and homogenous, and therefore the degrees of freedom for action memory are arguably of the same order of magnitude as the participant-identity and participant-location memory (albeit not having the same specific number of options). Beyond those shortcomings, there are also several advantages to using this test on clinical populations. Among them are: (1) the test is short and easily performed by stroke patients, who generally have short attention spans; (2) the test is widely accessible and is in common use worldwide, (3) it is well standardized, using a broad and representative sample of different age groups, (4) it is validated on various clinical samples, and (5) it has relatively good reliability and internal validity (Kent, 2013). Those advantages provide a reasonable basis for applying this test to clinical populations and make the current suggested analysis easily used.

## 5. Conclusions

The current study identifies a distributed set of cortical and sub-cortical regions supporting visual memory for persons, actions, and locations portrayed in a narrative scene. Those regions include extrastriate and other cortical areas associated with visual perception and information processing. Our finding of differential processing for each visual element are consistent with findings from functional imaging and lesion studies. Additionally, non-specific processing of those features was also demonstrated in other clusters in left temporo-occipital and right lateral temporo-parietal networks, presumably related to generic visual episodic memory processes. Importantly, the current study uses a more ecological visual episodic memory test to decompose complex visual memory into more basic neurocognitive components. In a diagnostic context, this may lead to better understanding of the subjective memory problems many patients experience in daily life.

## References

Abbott, V., Black, J. B., & Smith, E. E. (1985). The representation of scripts in memory. Journal of Memory and Language, 24(2), 179-199. https://doi.org/10.1016/0749-596X(85)90023-3

Abelson, R. P. (1981). Psychological status of the script concept. American Psychologist, 36(7), 715–729. https://doi.org/10.1037/0003-066X.36.7.715

Alba, J. W., & Hasher, L. (1983). Is memory schematic?. Psychological Bulletin, 93(2), 203–231. https://psycnet.apa.org/doi/10.1037/0033-2909.93.2.203

Bates, E., Wilson, S. M., Saygin, A. P., Dick, F., Sereno, M. I., Knight, R. T., & Dronker, N. F. (2003). Voxel-based lesion-symptom mapping. Nature Neuroscience, 6, 448–450. https://doi.org/10.1038/nn1050

Benjamini, Y., & Hochberg, Y. (1995). Controlling the false discovery rate: a practical and powerful approach to multiple testing. Journal of the Royal Statistical Society: Series B (Methodological), 57(1), 289–300. https://doi.org/10.1111/j.2517-6161.1995.tb02031.x

Ben-Zvi, S., Soroker, N., & Levy, D. A. (2015). Parietal lesion effects on cued recall following pair associate learning. Neuropsychologia, 73, 176–194. https://doi.org/10.1016/j.neuropsychologia.2015.05.009

Ben-Zvi Feldman, S., Soroker, N., & Levy, D. A. (2021). Lesion-behaviour mapping reveals multifactorial neurocognitive processes in recognition memory for unfamiliar faces. Neuropsychologia, 163, 108078. https://doi.org/10.1016/j.neuropsychologia.2021.108078

Binder, J. R., Desai, R. H., Graves, W. W., & Conant, L. L. (2009) Where is the semantic system? A critical review and meta-analysis of 120 functional neuroimaging studies. Cerebral Cortex, 19, 2767–2796. https://doi.org/10.1093/cercor/bhp055

Binder, E., Dovern, A., Hesse, M. D., Ebke, M., Karbe, H., Saliger, J., Fink, G. R., & Weiss, P. H. (2017). Lesion evidence for a human mirror neuron system. Cortex, 90, 125–137. https://doi.org/10.1016/j.cortex.2017.02.008

Bonasia, K., Sekeres, M. J., Gilboa, A., Grady, C. L., Winocur, G., & Moscovitch, M. (2018). Prior knowledge modulates the neural substrates of encoding and retrieving naturalistic events at short and long delays. Neurobiology of Learning and Memory, 153, 26–39. https://doi.org/10.1016/j.nlm.2018.02.017

Bone, M.B., Ahmad, F. & Buchsbaum, B.R. (2020). Feature-specific neural reactivation during episodic memory. Nature Communications, 11, 1945. https://doi.org/10.1038/s41467-020-15763-2

Buchsbaum, B. R., Ye, D., & D’Esposito, M. (2011). Recency effects in the inferior parietal lobe during verbal recognition memory. Frontiers in Human Neuroscience, 5, 59. https://doi.org/10.3389/fnhum.2011.00059

Buchsbaum, B. R., Lemire-Rodger, S., Fang, C., & Abdi, H. (2012). The neural basis of vivid memory is patterned on perception. Journal of Cognitive Neuroscience, 24(9), 1867–1883. https://doi.org/10.1162/jocn_a_00253

Burgess, N., Maguire, E. A., Spiers, H. J., & O’Keefe, J. (2001). A temporoparietal and prefrontal network for retrieving the spatial context of lifelike events. Neuroimage, 14(2), 439–453. https://doi.org/10.1006/nimg.2001.0806

Cabeza, R., Ciaramelli, E., Olson, I. R., & Moscovitch, M. (2008). The parietal cortex and episodic memory: an attentional account. Nature Reviews Neuroscience, 9(8), 613–625. https://doi.org/10.1038/nrn2459

Cabeza, R., Mazuz, Y. S., Stokes, J., Kragel, J. E., Woldorff, M. G., Ciaramelli, E., Olson, I. R., & Moscovitch, M. (2011). Overlapping parietal activity in memory and perception: Evidence for the attention to memory model. Journal of Cognitive Neuroscience, 23(11), 3209–3217. https://doi.org/10.1162/jocn_a_00065

Caspers, S., Zilles, K., Laird, A. R., & Eickhoff, S. B. (2010). ALE meta-analysis of action observation and imitation in the human brain. Neuroimage, 50(3), 1148–1167. https://doi.org/10.1016/j.neuroimage.2009.12.112

Cattaneo, L., & Rizzolatti, G. (2009). The mirror neuron system. Archives of Neurology, 66(5), 557–560. https://doi.org/10.1001/archneurol.2009.41

Chambers, C. D., Payne, J. M., Stokes, M. G., & Mattingley, J. B. (2004). Fast and slow parietal pathways mediate spatial attention. Nature Neuroscience, 7(3), 217–218. https://doi.org/10.1038/nn1203

Ciaramelli, E., Grady, C., Levine, B., Ween, J., & Moscovitch, M. (2010). Top-down and bottom-up attention to memory are dissociated in posterior parietal cortex: neuroimaging and neuropsychological evidence. Journal of Neuroscience, 30(14), 4943–4956. https://doi.org/10.1523/JNEUROSCI.1209-09.2010

Ciaramelli, E., Grady, C. L., & Moscovitch, M. (2008). Top-down and bottom-up attention to memory: a hypothesis (AtoM) on the role of the posterior parietal cortex in memory retrieval. Neuropsychologia, 46(7), 1828–1851. https://doi.org/10.1016/j.neuropsychologia.2008.03.022

Clewett, D., DuBrow, S., & Davachi, L. (2019). Transcending time in the brain: How event memories are constructed from experience. Hippocampus, 29(3), 162–183. https://doi.org/10.1002/hipo.23074

Cona, G., & Scarpazza, C. (2019). Where is the “where” in the brain? A meta-analysis of neuroimaging studies on spatial cognition. Human Brain Mapping, 40(6), 1867–1886. https://doi.org/10.1002/hbm.24496

Crane, J., & Milner, B. (2005). What went where? Impaired object-location learning in patients with right hippocampal lesions. Hippocampus, 15(2), 216–231. https://doi.org/10.1002/hipo.20043

D’Esposito, M., Detre, J. A., Aguirre, G. K., Stallcup, M., Alsop, D. C., Tippet, L. J., & Farah, M. J. (1997). A functional MRI study of mental image generation. Neuropsychologia, 35(5), 725–730. https://doi.org/10.1016/S0028-3932(96)00121-2

Diana, R. A., Yonelinas, A. P., & Ranganath, C. (2007). Imaging recollection and familiarity in the medial temporal lobe: a three-component model. Trends in Cognitive Sciences, 11(9), 379–386. https://doi.org/10.1016/j.tics.2007.08.001

Fabbri-Destro, M., & Rizzolatti, G. (2008). Mirror neurons and mirror systems in monkeys and humans. Physiology, 23(3), 171–179. https://doi.org/10.1152/physiol.00004.2008

Farah, M. J., Hammond, K. M., Levine, D. N., & Calvanio, R. (1988). Visual and spatial mental imagery: Dissociable systems of representation. Cognitive Psychology, 20(4), 439–462. https://doi.org/10.1016/0010-0285(88)90012-6

Frenkel-Toledo, S., Fridberg, G., Ofir, S., Bartur, G., Lowenthal-Raz, J., Granot, O., Handelzalts, S., & Soroker, N. (2019). Lesion location impact on functional recovery of the hemiparetic upper limb. PLoS ONE 14(7), e0219738. https://doi.org/10.1371/journal.pone.0219738

Gaffan, D. (1998). Idiothetic input into object-place configuration as the contribution to memory of the monkey and human hippocampus: a review. Experimental Brain Research, 123(1), 201–209. https://doi.org/10.1007/s002210050562

Gallivan, J. P., & Goodale, M. A. (2018). The dorsal “action” pathway. Handbook of Clinical Neurology, 151, 449–466. https://doi.org/10.1016/B978-0-444-63622-5.00023-1

Genovese, C. R., Lazar, N. A., & Nichols, T. (2002). Thresholding of statistical maps in functional neuroimaging using the false discovery rate. Neuroimage, 15(4), 870–878. https://doi.org/10.1006/nimg.2001.1037

Gilboa, A, & Marlatte H. (2017). Neurobiology of schemas and schema-mediated memory. Trends in Cognitive Sciences, 21, 618–631. http://doi:10.1016/j.tics.2017.04.013

Gilmore, A. W., Quach, A., Kalinowski, S. E., Gotts, S. J., Schacter, D. L., & Martin, A. (2021). Dynamic content reactivation supports naturalistic autobiographical recall in humans. Journal of Neuroscience, 41(1), 153–166. https://doi.org/10.1523/JNEUROSCI.1490-20.2020

Grossi, D., Modafferi, A., Pelosi, L., & Trojano, L. (1989). On the different roles of the cerebral hemispheres in mental imagery: The “O’Clock Test” in two clinical cases. Brain and Cognition, 10(1), 18–27. https://doi.org/10.1016/0278-2626(89)90072-9

Hales, J. B., & Brewer, J. B. (2013). Parietal and frontal contributions to episodic encoding of location. Behavioural Brain Research, 243, 16–20. https://doi.org/10.1016/j.bbr.2012.12.048

Haramati, S., Soroker, N., Dudai, Y., & Levy, D. A. (2008). The posterior parietal cortex in recognition memory: A neuropsychological study. Neuropsychologia, 46, 1756–1766. https://doi.org/10.1016/j.neuropsychologia.2007.11.015

Hayama, H. R., Vilberg, K. L., & Rugg, M. D. (2012). Overlap between the neural correlates of cued recall and source memory: evidence for a generic recollection network?. Journal of Cognitive Neuroscience, 24(5), 1127–1137. https://doi.org/10.1162/jocn_a_00202

Henderson, J. M., & Hollingworth, A. (1999). High-level scene perception. Annual Review of Psychology, 50(1), 243–271. https://doi.org/10.1146/annurev.psych.50.1.243

Hoeft, F., Barnea-Goraly, N., Haas, B. W., Golarai, G., Ng, D., Mills, D., Korenberg, J., Bellugi, U., Galaburda A., & Reiss, A. L. (2007). More is not always better: increased fractional anisotropy of superior longitudinal fasciculus associated with poor visuospatial abilities in Williams syndrome. Journal of Neuroscience, 27(44), 11960–11965. https://doi.org/10.1523/JNEUROSCI.3591-07.2007

Holdstock, J. S., Mayes, A. R., Gong, Q. Y., Roberts, N., & Kapur, N. (2005). Item recognition is less impaired than recall and associative recognition in a patient with selective hippocampal damage. Hippocampus, 15(2), 203–215. https://doi.org/10.1002/hipo.20046

Holdstock, J. S., Mayes, A. R., Roberts, N., Cezayirli, E., Isaac, C. L., O’reilly, R. C., & Norman, K. A. (2002). Under what conditions is recognition spared relative to recall after selective hippocampal damage in humans?. Hippocampus, 12(3), 341–351. https://doi.org/10.1002/hipo.10011

Hower, K. H., Wixted, J., Berryhill, M. E., & Olson, I. R. (2014). Impaired perception of mnemonic oldness, but not mnemonic newness, after parietal lobe damage. Neuropsychologia, 56, 409–417. https://doi.org/10.1016/j.neuropsychologia.2014.02.014

Hudson, J. A., Fivush, R., & Kuebli, J. (1992). Scripts and episodes: The development of event memory. Applied Cognitive Psychology, 6(6), 483–505. https://doi.org/10.1002/acp.2350060604

Iacoboni, M., Molnar-Szakacs, I., Gallese, V., Buccino, G., Mazziotta, J. C., & Rizzolatti, G. (2005). Grasping the intentions of others with one’s own mirror neuron system. PLoS Biology, 3(3), e79. https://doi.org/10.1371/journal.pbio.0030079

Iacoboni, M., Woods, R. P., Brass, M., Bekkering, H., Mazziotta, J. C., & Rizzolatti, G. (1999). Cortical mechanisms of human imitation. Science, 286(5449), 2526–2528. https://doi.org/10.1126/science.286.5449.2526

Johnson, J. D., McDuff, S. G., Rugg, M. D., & Norman, K. A. (2009). Recollection, familiarity, and cortical reinstatement: A multivoxel pattern analysis. Neuron, 63(5), 697–708. https://doi.org/10.1016/j.neuron.2009.08.011

Johnson, J. D., & Rugg, M. D. (2007). Recollection and the reinstatement of encoding-related cortical activity. Cerebral Cortex, 17, 2507–2515. https://doi.org/10.1093/cercor/bhl156

Kent, P. (2013). The evolution of the Wechsler Memory Scale: A selective review. Applied Neuropsychology: Adult, 20(4), 277–291. https://doi.org/10.1080/09084282.2012.689267

Kessels, R. P., Postma, A., Kappelle, L. J., & de Haan, E. H. (2000). Spatial memory impairment in patients after tumour resection: evidence for a double dissociation. Journal of Neurology, Neurosurgery & Psychiatry, 69(3), 389–391. http://dx.doi.org/10.1136/jnnp.69.3.389

Kim, S. H., Jeon, H. E., & Park, C. H. (2020). Relationship between visual perception and microstructural change of the superior longitudinal fasciculus in patients with brain injury in the right hemisphere: a preliminary diffusion tensor tractography study. Diagnostics, 10(9), 641. https://doi.org/10.3390/diagnostics10090641

Köhler, S., Moscovitch, M., Winocur, G., Houle, S., & McIntosh, A. R. (1998). Networks of domain-specific and general regions involved in episodic memory for spatial location and object identity. Neuropsychologia, 36(2), 129–142. https://doi.org/10.1016/S0028-3932(97)00098-5

Konen, C. S., & Kastner, S. (2008). Two hierarchically organized neural systems for object information in human visual cortex. Nature Neuroscience, 11(2), 224–231. https://doi.org/10.1038/nn2036

Kopelman, M. D., Stanhope, N., & Kingsley, D. (1997). Temporal and spatial context memory in patients with focal frontal, temporal lobe, and diencephalic lesions. Neuropsychologia, 35(12), 1533–1545. https://doi.org/10.1016/S0028-3932(97)00076-6

Lancaster, J. L., Woldorff, M. G., Parsons, L. M., Liotti, M., Freitas, C. S., Rainey, L., Kochunov, P. V., Nickerson, D., Mikiten, S. A., & Fox, P. T. (2000). Automated Talairach atlas labels for functional brain mapping. Human Brain Mapping, 10, 120–131. https://doi.org/10.1002/1097-0193(200007)10:3<120::AID-HBM30>3.0.CO;2-8

Lee, H., Chun, M. M., & Kuhl, B. A. (2017). Lower parietal encoding activation is associated with sharper information and better memory. Cerebral Cortex, 27(4), 2486–2499. https://doi.org/10.1093/cercor/bhw097

Lehky, S. R., Peng, X., McAdams, C. J., & Sereno, A. B. (2008). Spatial modulation of primate inferotemporal responses by eye position. PLoS One, 3(10), e3492. https://doi.org/10.1371/journal.pone.0003492

Levy, D. A. (2012). Towards an understanding of parietal mnemonic processes: Some conceptual guideposts. Frontiers in Integrative Neuroscience, 6, 41. https://doi.org/10.3389/fnint.2012.00041

Lo, R., Gitelman, D., Levy, R., Hulvershorn, J., & Parrish, T. (2010). Identification of critical areas for motor function recovery in chronic stroke subjects using voxel-based lesion symptom mapping. Neuroimage, 49, 9–18. https://doi.org/10.1016/j.neuroimage.2009.08.044

Luzzatti, C., Vecchi, T., Agazzi, D., Cesa-Bianchi, M., & Vergani, C. (1998). A neurological dissociation between preserved visual and impaired spatial processing in mental imagery. Cortex, 34(3), 461–469. https://doi.org/10.1016/S0010-9452(08)70768-8

Mangun, G. R., Buonocore, M. H., Girelli, M., & Jha, A. P. (1998). ERP and fMRI measures of visual spatial selective attention. Human Brain Mapping, 6(5-6), 383–389. https://doi.org/10.1002/(SICI)1097-0193(1998)6:5/6<383::AID-HBM10>3.0.CO;2-Z

Manns, J. R., & Eichenbaum, H. (2006). Evolution of declarative memory. Hippocampus, 16(9), 795–808. https://doi.org/10.1002/hipo.20205

Marin, D., Madotto, E., Fabbro, F., Skrap, M., & Tomasino, B. (2017). Design fluency and neuroanatomical correlates in 54 neurosurgical patients with lesions to the right hemisphere. Journal of Neuro-oncology, 135(1), 141–150. https://doi.org/10.1007/s11060-017-2560-3

Masís-Obando, R., Norman, K. A., & Baldassano, C. (2022). Schema representations in distinct brain networks support narrative memory during encoding and retrieval. Elife, 11, e70445. https://doi.org/10.7554/eLife.70445

Memel, M., Wank, A. A., Ryan, L., & Grilli, M. D. (2020). The relationship between episodic detail generation and anterotemporal, posteromedial, and hippocampal white matter tracts. Cortex, 123, 124–140. https://doi.org/10.1016/j.cortex.2019.10.010

Moon, H., Pyun, S. B., Tae, W. S., & Kwon, H. K. (2016). Neural substrates of lower extremity motor, balance, and gait function after supratentorial stroke using voxel-based lesion symptom mapping. Neuroradiology, 58(7), 723–731. https://doi.org/10.1007/s00234-016-1672-3

Mori, S., Oishi, K., Jiang, H., Jiang, L., Li, X., Akhter, K., Hua, K., Faria, A. V., Mahmood, A., Woods, R., Toga, A. W., Pike, G. B., Neto, P. R., Evans, A., Zhang, J., Huang, H., Miller, M. I., van Zijl, P., & Mazziotta, J. (2008). Stereotaxic white matter atlas based on diffusion tensor imaging in an ICBM template. NeuroImage, 40(2), 570–582. https://doi.org/10.1016/j.neuroimage.2007.12.035

Moscovitch, C., Kapur, S., Köhler, S., & Houle, S. (1995). Distinct neural correlates of visual long-term memory for spatial location and object identity: a positron emission tomography study in humans. Proceedings of the National Academy of Sciences, 92(9), 3721–3725. https://doi.org/10.1073/pnas.92.9.3721

Murata, A., Gallese, V., Luppino, G., Kaseda, M., & Sakata, H. (2000). Selectivity for the shape, size, and orientation of objects for grasping in neurons of monkey parietal area AIP. Journal of Neurophysiology, 83(5), 2580–2601. https://doi.org/10.1152/jn.2000.83.5.2580

Nichols T. E., & Holmes, A. P. (2002). Nonparametric permutation tests for functional neuroimaging: A primer with examples. Human Brain Mapping, 15(1), 1–25. https://doi.org/10.1002/hbm.1058

Nunn, J. A., Graydon, F. J. X., Polkey, C. E., & Morris, R. G. (1999). Differential spatial memory impairment after right temporal lobectomy demonstrated using temporal titration. Brain, 122(1), 47–59. https://doi.org/10.1093/brain/122.1.47

O’Connor, A. R., Han, S., & Dobbins, I. G. (2010). The inferior parietal lobule and recognition memory: expectancy violation or successful retrieval?. Journal of Neuroscience, 30(8), 2924–2934. https://doi.org/10.1523/JNEUROSCI.4225-09.2010

Paulus, M. P., Feinstein, J. S., Leland, D., & Simmons, A. N. (2005). Superior temporal gyrus and insula provide response and outcome-dependent information during assessment and action selection in a decision-making situation. Neuroimage, 25(2), 607–615. https://doi.org/10.1016/j.neuroimage.2004.12.055

Pigott, S., & Milner, B. (1993). Memory for different aspects of complex visual scenes after unilateral temporal-or frontal-lobe resection. Neuropsychologia, 31(1), 1–15. https://doi.org/10.1016/0028-3932(93)90076-C

Postma, A., Kessels, R. P., & van Asselen, M. (2008). How the brain remembers and forgets where things are: The neurocognition of object–location memory. Neuroscience & Biobehavioral Reviews, 32(8), 1339–1345. https://doi.org/10.1016/j.neubiorev.2008.05.001

Price, A. R., Bonner, M. F. & Grossman, M. (2015) Semantic memory: Cognitive and neuroanatomical perspectives. In: Toga, A. W. (Ed.). Brain Mapping: An Encyclopedic Reference, vol. 3 (pp. 529–536). Academic Press: Elsevier. https://doi.org/10.1016/B978-0-12-397025-1.00280-3

Ranganath, C. (2010). Binding items and contexts: The cognitive neuroscience of episodic memory. Current Directions in Psychological Science, 19(3), 131–137. https://doi.org/10.1177/0963721410368805

Riccardi, N., Yourganov, G., Rorden, C., Fridriksson, J., & Desai, R. H. (2019). Dissociating action and abstract verb comprehension post-stroke. Cortex, 120, 131–146. https://doi.org/10.1016/j.cortex.2019.05.013

Richter, F. R., Cooper, R. A., Bays, P. M., & Simons, J. S. (2016). Distinct neural mechanisms underlie the success, precision, and vividness of episodic memory. eLife, 5, e18260. http://doi.org/10.7554/eLife.18260.

Rizzolatti, G., & Craighero, L. (2004). The mirror-neuron system. Annual Review of Neuroscience, 27, 169–192. https://doi.org/10.1146/annurev.neuro.27.070203.144230

Rizzolatti, G., & Fabbri-Destro, M. (2010). Mirror neurons: from discovery to autism. Experimental Brain Research, 200(3), 223–237. https://doi.org/10.1007/s00221-009-2002-3

Rizzolatti, G., Fogassi, L., & Gallese, V. (2001). Neurophysiological mechanisms underlying the understanding and imitation of action. Nature Reviews Neuroscience, 2(9), 661–670. https://doi.org/10.1038/35090060

Rizzolatti, G., & Sinigaglia, C. (2010). The functional role of the parieto-frontal mirror circuit: interpretations and misinterpretations. Nature Reviews Neuroscience, 11(4), 264–274. https://doi.org/10.1038/nrn2805

Rosenbaum, R. S., Gilboa, A., & Moscovitch, M. (2014). Case studies continue to illuminate the cognitive neuroscience of memory. Annals of the New York Academy of Sciences, 1316, 105–133. https://doi.org/10.1111/nyas.12467

Ross, R. S., & Slotnick, S. D. (2008). The hippocampus is preferentially associated with memory for spatial context. Journal of Cognitive Neuroscience, 20(3): 432–446. https://doi.org/10.1162/jocn.2008.20035

Rugg, M. D., & King, D. R. (2018). Ventral lateral parietal cortex and episodic memory retrieval. Cortex, 107, 238–250. https://doi.org/10.1016/j.cortex.2017.07.012

Rugg, M. D., & Vilberg, K. L. (2013). Brain networks underlying episodic memory retrieval. Current Opinion in Neurobiology, 23(2), 255–260. https://doi.org/10.1016/j.conb.2012.11.005

Saxe, R., Xiao, D. K., Kovacs, G., Perrett, D. I., & Kanwisher, N. (2004). A region of right posterior superior temporal sulcus responds to observed intentional actions. Neuropsychologia, 42(11), 1435–1446. https://doi.org/10.1016/j.neuropsychologia.2004.04.015

Schacter, D. L., Benoit, R. G., & Szpunar, K. K. (2017). Episodic future thinking: Mechanisms and functions. Current Opinion in Behavioral Sciences, 17, 41–50. https://doi.org/10.1016/j.cobeha.2017.06.002

Schacter, D. L., Reiman, E., Uecker, A., Roister, M. R., Yun, L. S., & Cooper, L. A. (1995). Brain regions associated with retrieval of structurally coherent visual information. Nature, 376(6541), 587–590. https://doi.org/10.1038/376587a0

Schmahmann, J. D., Smith, E. E., Eichler, F. S., & Filley, C. M. (2008). Cerebral white matter: Neuroanatomy, clinical neurology, and neurobehavioral correlates. Annals of the New York Academy of Sciences, 1142(1), 266–309. https://doi.org/10.1196/annals.1444.017

Schoch, B., Dimitrova, A., Gizewski, E. R., & Timmann, D. (2006). Functional localization in the human cerebellum based on voxelwise statistical analysis: A study of 90 patients. Neuroimage, 30(1), 36–51. https://doi.org/10.1016/j.neuroimage.2005.09.018

Sereno, A. B., & Amador, S. C. (2006). Attention and memory-related responses of neurons in the lateral intraparietal area during spatial and shape-delayed match-to-sample tasks. Journal of Neurophysiology, 95(2), 1078–1098. https://doi.org/10.1152/jn.00431.2005

Sereno, A. B., & Lehky, S. R. (2011). Population coding of visual space: comparison of spatial representations in dorsal and ventral pathways. Frontiers in Computational Neuroscience, 4, 159. https://doi.org/10.3389/fncom.2010.00159

Sereno, A. B., & Maunsell, J. H. (1998). Shape selectivity in primate lateral intraparietal cortex. Nature, 395(6701), 500–503. https://doi.org/10.1038/26752

Shimamura, A. P. (2011). Episodic retrieval and the cortical binding of relational activity. Cognitive, Affective, & Behavioral Neuroscience, 11, 277–291. https://doi.org/10.3758/s13415-011-0031-4

Siddiqi, S. H., Kording, K. P., Parvizi, J. & Fox, M. D. (2022). Causal mapping of human brain function. Nature Reviews Neuroscience, 23, 361–375. https://doi.org/10.1038/s41583-022-00583-8

Simons, J. S., Peers, P. V., Mazuz, Y. S., Berryhill, M. E., & Olson, I. R. (2010). Dissociation between memory accuracy and memory confidence following bilateral parietal lesions. Cerebral Cortex, 20, 479–485. https://doi.org/10.1093/cercor/bhp116

Slotnick, S. D., Moo, L. R., Segal, J. B., & Hart Jr, J. (2003). Distinct prefrontal cortex activity associated with item memory and source memory for visual shapes. Cognitive Brain Research, 17(1), 75–82. https://doi.org/10.1016/S0926-6410(03)00082-X

Smith, M. L., & Milner, B. (1981). The role of the right hippocampus in the recall of spatial location. Neuropsychologia, 19(6), 781–793. https://doi.org/10.1016/0028-3932(81)90090-7

Smith, M. L., & Milner, B. (1984). Differential effects of frontal-lobe lesions on cognitive estimation and spatial memory. Neuropsychologia, 22(6), 697–705. https://doi.org/10.1016/0028-3932(84)90096-4

Solomon, J., Raymont, V., Braun, A., Butman, J. A., & Grafman, J. (2007). User-friendly software for the analysis of brain lesions (ABLe). Computer Methods and Programs in Biomedicine, 86, 245–254. https://doi.org/10.1016/j.cmpb.2007.02.006

Spagna, A., Hajhajate, D., Liu, J., & Bartolomeo, P. (2021). Visual mental imagery engages the left fusiform gyrus, but not the early visual cortex: a meta-analysis of neuroimaging evidence. Neuroscience & Biobehavioral Reviews, 122, 201–217. https://doi.org/10.1016/j.neubiorev.2020.12.029

Stepankova, K., Fenton, A. A., Pastalkova, E., Kalina, M., & Bohbot, V. D. (2004). Object– location memory impairment in patients with thermal lesions to the right or left hippocampus. Neuropsychologia, 42(8), 1017–1028. https://doi.org/10.1016/j.neuropsychologia.2004.01.002

Thorudottir, S., Sigurdardottir, H. M., Rice, G. E., Kerry, S. J., Robotham, R. J., Leff, A. P., & Starrfelt, R. (2020). The architect who lost the ability to imagine: The cerebral basis of visual imagery. Brain Sciences, 10(2), 59. https://doi.org/10.3390/brainsci10020059

Tzourio-Mazoyer, N., Landeau, B., Papathanassiou, D., Crivello, F., Etard, O., Delcroix, N., Mazoyer, B., & Joliot, M. (2002). Automated anatomical labeling of activations in SPM using a macroscopic anatomical parcellation of the MNI MRI single-subject brain. Neuroimage, 15, 273–289. https://doi.org/10.1006/nimg.2001.0978

Vaidya, C. J., Zhao, M., Desmond, J. E., & Gabrieli, J. D. (2002). Evidence for cortical encoding specificity in episodic memory: memory-induced re-activation of picture processing areas. Neuropsychologia, 40(12), 2136–2143. https://doi.org/10.1016/S0028-3932(02)00053-2

van Asselen, M., Kessels, R. P., Frijns, C. J., Kappelle, L. J., Neggers, S. F., & Postma, A. (2009). Object-location memory: A lesion-behavior mapping study in stroke patients. Brain and Cognition, 71(3), 287–294. https://doi.org/10.1016/j.bandc.2009.07.012

van Overwalle, F. (2009), Social cognition and the brain: A meta-analysis. Human Brain Mapping, 30, 829–858. https://doi.org/10.1002/hbm.20547

Vargha-Khadem, F., Gadian, D. G., Watkins, K. E., Connelly, A., Van Paesschen, W., & Mishkin, M. (1997). Differential effects of early hippocampal pathology on episodic and semantic memory. Science, 277(5324), 376–380. https://doi.org/10.1126/science.277.5324.376

Vilberg, K. L., & Rugg, M. D. (2008). Memory retrieval and the parietal cortex: A review of evidence from a dual-process perspective. Neuropsychologia, 46, 1787–1799. https://doi.org/10.1016/j.neuropsychologia.2008.01.004

Vilberg, K. L., & Rugg, M. D. (2009a). Functional significance of retrieval-related activity in lateral parietal cortex: Evidence from fMRI and ERPs. Human Brain Mapping, 30, 1490–1501. https://doi.org/10.1002/hbm.20618

Vilberg, K. L., & Rugg, M. D. (2009b). An investigation of the effects of relative probability of old and new test items on the neural correlates of successful and unsuccessful source memory. NeuroImage, 45, 562–571. https://doi.org/10.1016/j.neuroimage.2008.12.020

Vilberg, K. L., & Rugg, M. D. (2012). The neural correlates of recollection: transient versus sustained fMRI effects. Journal of Neuroscience, 32(45), 15679–15687. https://doi.org/10.1523/JNEUROSCI.3065-12.2012

Wechsler, D. (1997). Wechsler Memory Scale (WMS-III). Psychological Corporation, San Antonio, Texas.

Wheeler, M. E., & Buckner, R. L. (2003). Functional dissociation among components of remembering: control, perceived oldness, and content. Journal of Neuroscience, 23(9), 3869–3880. https://doi.org/10.1523/JNEUROSCI.23-09-03869.2003

Wheeler, M. E., Petersen, S. E., & Buckner, R. L. (2000). Memory’s echo: vivid remembering reactivates sensory-specific cortex. Proceedings of the National Academy of Sciences, 97(20), 11125–11129. https://doi.org/10.1073/pnas.97.20.11125

Wing, E. A., Ritchey, M., Cabeza, R. (2015). Reinstatement of individual past events revealed by the similarity of distributed activation patterns during encoding and retrieval. Journal of Cognitive Neuroscience, 27(4): 679–691. https://doi.org/10.1162/jocn_a_00740

Wu, O., Cloonan, L., Mocking, S. J., Bouts, M. J., Copen, W. A., Cougo-Pinto, P. T., Fitzpatrick, K., Kanakis, A., Schaefer, P. W., Rosand, J., Furie, K. L., & Rost, N. S. (2015). Role of acute lesion topography in initial ischemic stroke severity and long-term functional outcomes. Stroke, 46(9), 2438–2444. https://doi.org/10.1161/STROKEAHA.115.009643

Wurm, M. F., Ariani, G., Greenlee, M. W., & Lingnau, A. (2016). Decoding concrete and abstract action representations during explicit and implicit conceptual processing. Cerebral Cortex, 26(8), 3390–3401. https://doi.org/10.1093/cercor/bhv169

Wurm, M. F., Caramazza, A., & Lingnau, A. (2017). Action categories in lateral occipitotemporal cortex are organized along sociality and transitivity. Journal of Neuroscience, 37(3), 562–575. https://doi.org/10.1523/JNEUROSCI.1717-16.2016

Wurm, M. F., & Lingnau, A. (2015). Decoding actions at different levels of abstraction. Journal of Neuroscience, 35(20), 7727–7735. https://doi.org/10.1523/JNEUROSCI.0188-15.2015

Xue, G. (2018). The neural representations underlying human episodic memory. Trends in Cognitive Sciences, 22(6), 544–561. https://doi.org/10.1016/j.tics.2018.03.004

Zimmermann, K., & Eschen, A. (2017). Brain regions involved in subprocesses of small-space episodic object-location memory: A systematic review of lesion and functional neuroimaging studies. Memory, 25(4), 487–519. https://doi.org/10.1080/09658211.2016.1188965

