## Supplementary files for "Brain Substrates of Episodic Memory for Identity, Location, and Action Information: A Lesion-Behavior Mapping Study"

**Supplementary Materials**

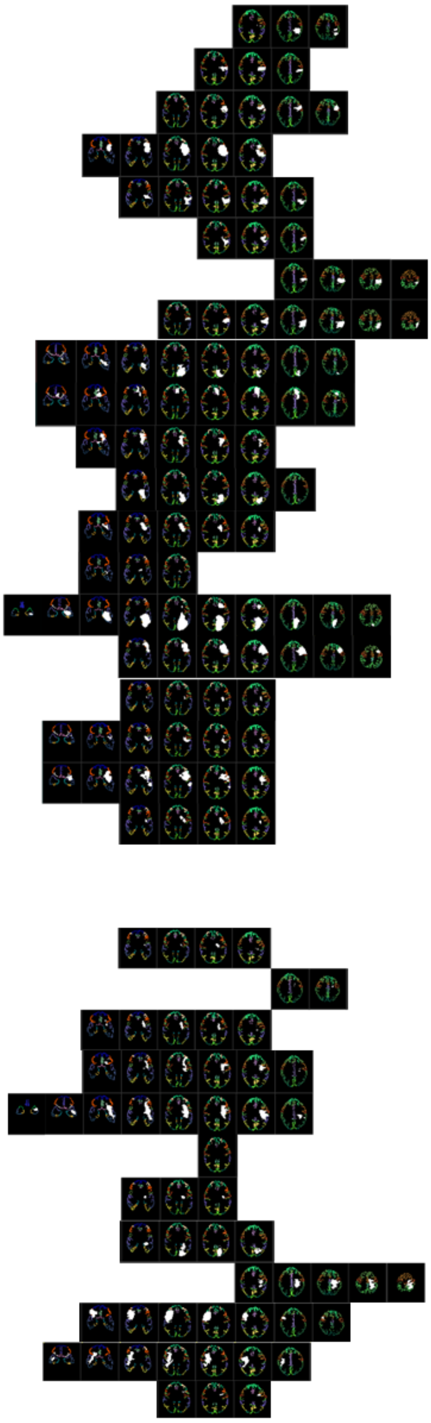

2006
2007
2008
2009
2010
2011
2012
2013
2014
2015
2016
2017
2018
2019
2020
2021
2022
2023
2024
2025
2026
2027
2028
2029
2030
2031
2032
2033
2034
2035
2036
2037
2038
2039

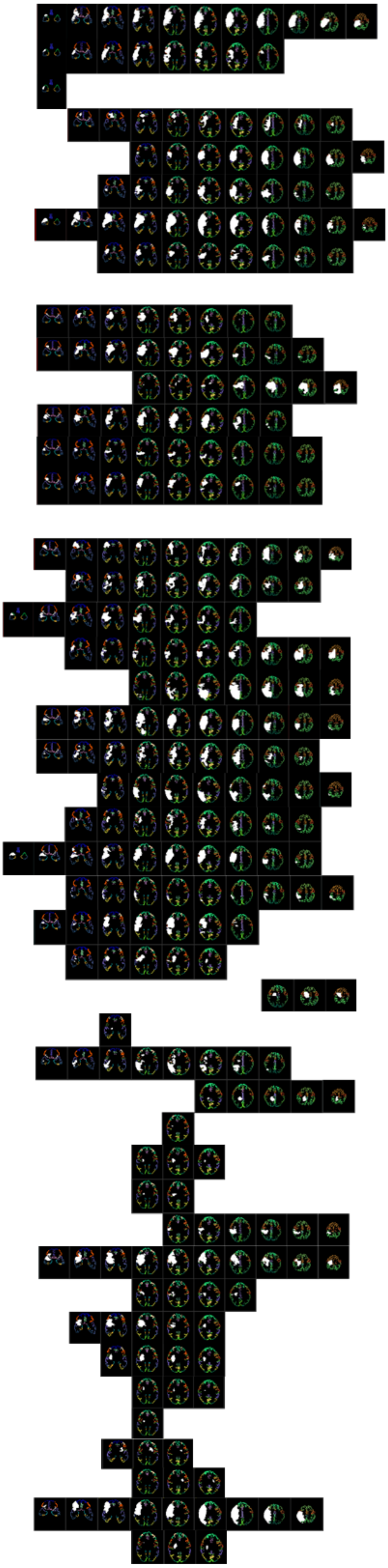

1002
1004
1008
1010
1011

**Figure S1. Individual lesion data**

Each patient's lesion is marked on an array of 11 standard templates. Displays follow neurological conventions, i.e., right-sided damage displayed on the left and left-sided on the right side. Only slices containing brain damage are shown. In the case of a minimal lesion, the MEDx system does not depict the lesion in the restricted set of standard templates used to present the structural damage. Data required for the preparation of this figure were not available for 7 RHD and 5 LHD patients.

**Table S1. Demographic and clinical characteristics of individual patients.**

| Participant | Age/ Gender | Education (years) | Lesion type | Lesion volume (cc) | Registration Accuracy (%) | TAO (weeks) | MI | SI | VFD | Neglect | Aphasia |
|---|---|---|---|---|---|---|---|---|---|---|---|
| **A. Right Hemisphere Damage (RHD)** | | | | | | | | | | | |
| 1001 | 40/F | 14 | H | NA | NA | 10 | ++ | +/e | −/e | + | - |
| 1002* | 61/F | 17 | I | 229.72 | 91.47 | 13 | ++ | ++ | −/e | + | - |
| 1003 | 29/F | 17 | I | NA | NA | 4 | + | + | -/e | + | - |
| 1004 | 63/M | 8 | I | 88.37 | 95.22 | 4 | + | +/e | -/e | + | - |
| 1005* | 63/F | 12 | H | NA | NA | 3 | + | +/e | - | + | - |
| 1006* | 57/M | 15 | I | NA | NA | 6 | ++ | + | -/e | - | - |
| 1007 | 76/M | 16 | I | NA | NA | 7 | NA | NA | NA | + | - |
| 1008 | 72/M | 16 | I/H | 138.82 | 94.96 | 3 | + | +/e | -/e | + | - |
| 1009 | 82/F | 15 | NA | NA | NA | 5 | + | + | - | + | - |
| 1010 | 66/F | 9 | I | 60.3 | 95.21 | 13 | + | + | -/e | + | - |
| 1011 | 60/F | 11 | I | 107 | 94.4 | 5 | ++ | + | -/e | + | - |
| 1012* | 26/F | 12 | AVM-H | NA | NA | 5 | + | + | - | - | - |
| 1013 | 58/M | 12 | I | 93.62 | 94.64 | 12 | + | + | + | - | - |
| 1014 | 60/M | 12 | I | 278 | 91.29 | 9 | + | + | -/e | - | - |
| 1015 | 62/F | 12 | I | 53.6 | 95.29 | 10 | + | + | - | + | - |
| 1016 | 63/M | 12 | I | 61.3 | 95.15 | 3 | ++ | - | -/e | + | - |
| 1017 | 64/M | 10 | I | 47.8 | 94.51 | 5 | + | - | -/e | + | - |
| 1018 | 44/F | 17 | I/H | 117 | 94.66 | 9 | + | + | -/e | + | - |
| 1019 | 61/M | 15 | I | 72.3 | 95 | 12 | + | + | - | + | - |
| 1020 | 60/M | 19 | I | 139 | 94.49 | 5 | - | + | - | + | - |
| 1021 | 66/M | 15 | H | 24 | 95.18 | 14 | + | - | - | + | - |
| 1022 | 76/F | 8 | I | 43.5 | 94.58 | 12 | + | + | - | + | - |
| 1023 | 75/M | 10 | I | 46.4 | 95.33 | 8 | + | - | - | - | - |
| 1024 | 73/M | 16 | I | 137 | 93.98 | 10 | + | - | - | + | - |
| 1025 | 79/M | 12 | H | 118 | 94.88 | 8 | + | + | -/e | + | - |
| 1026* | 40/M | 16 | H | 69.6 | 95.04 | 16 | + | - | - | + | - |
| 1027 | 79/F | 16 | I | 117 | 94.41 | 12 | + | + | - | + | - |
| 1028 | 65/M | 10 | H | 137 | 94.16 | 11 | + | - | - | + | - |
| 1029 | 71/M | 12 | I | 170 | 95.03 | 15 | + | ++ | + | + | - |
| 1030 | 63/M | 8 | H | 96.6 | 94.19 | 8 | + | + | - | + | - |
| 1031 | 65/M | 14 | I | 92.4 | 94.31 | 10 | - | - | -/e | - | - |
| 1032 | 60/M | 18 | I | 52.4 | 94.44 | 9 | + | + | -/e | + | - |
| 1033* | 57/M | 20 | I/H | 113 | 88.95 | 13 | ++ | + | -/e | + | - |
| 1034 | 56/F | 18 | I | 24.3 | 94.53 | 6 | + | + | -/e | + | - |
| 1035 | 66/M | 16 | I | 90.57 | 94.01 | 9 | + | - | - | + | - |
| 1036 | 28/F | 16 | I | 22.69 | 94.71 | 9 | + | - | - | - | - |
| 1037 | 58/M | 12 | H | 26.5 | 94.63 | 6 | + | - | - | - | - |
| 1038 | 43/M | 11 | I | 0.67 | 95.02 | 13 | + | - | - | - | - |
| 1039 | 57/F | 12 | I | 47.13 | 94.12 | 25 | + | - | - | - | - |
| 1040 | 60/F | 18 | H | 17.86 | 92.57 | 4 | + | + | - | - | - |
| 1041* | 72/M | 9 | I | 0.39 | 95.08 | 5 | + | + | - | - | - |
| 1042* | 55/M | 20+ | I | 5.44 | 94.8 | 7 | + | - | - | - | - |
| 1043 | 65/M | 17 | I | 3.41 | 93.83 | 3 | + | - | - | - | - |
| 1044 | 66/F | 14 | H | 25.76 | 94.33 | 4 | + | + | + | - | - |
| 1045 | 45/F | 14 | I | 125.09 | 94.19 | 8 | + | + | - | - | - |
| 1046 | 53/M | 12 | H | 10.03 | 94.49 | 6 | + | + | - | - | - |
| 1047 | 46/M | 12 | I | 41.56 | 95.05 | 4 | - | - | - | - | - |
| 1048 | 46/M | 16 | H | 22.27 | 93.96 | 5 | + | + | - | - | - |
| 1049 | 66/M | 11 | I | 2.07 | 94.28 | 8 | + | - | - | - | - |
| 1050 | 59/M | 9.5 | I | 0.93 | 92.53 | 23 | - | - | - | - | - |
| 1051 | 74/M | 15 | I | 86.8 | 94.63 | 8 | + | + | - | + | - |
| 1052 | 60/F | 9 | H | 68.23 | 92.38 | 9 | + | + | + | + | - |
| 1053 | 60/M | 9 | I | 202.12 | 94.13 | 7.1 | + | + | - | + | - |
| 1054 | 74/M | 16 | I | 5.99 | 94.47 | 13 | + | + | - | - | - |

**B. Left Hemisphere Damage (LHD)**

| | | | | | | | | | | |
|---|---|---|---|---|---|---|---|---|---|---|
| 2001 | 43/M | 15 | H | 58.1 | NA | 12 | + | + | - | - | - |
| 2002 | 77/F | 13 | I | 45.5 | NA | 12 | + | - | - | - | + |
| 2003 | 32/F | 12 | AVM-H | NA | NA | 4 | - | + | - | - | - |
| 2004 | 32/F | 16 | CVST-H | 6.1 | NA | 5 | - | - | - | - | - |
| 2005 | 27/F | 13 | CVST-H | 112 | NA | 15 | ++ | + | - | - | - |
| 2006 | 75/F | 12 | I | 9.08 | 94.44 | 12 | + | + | - | - | + |
| 2007 | 57/M | 12 | I | 10.4 | 94.45 | 4 | - | + | - | - | + |
| 2008* | 65/M | 20 | I | 22.9 | 94.42 | 3 | - | + | - | - | + |
| 2009 | 58/M | 15 | I | 67.7 | 95.21 | 11 | + | + | - | - | + |
| 2010 | 70/F | 16 | I/H | 34.3 | 93.9 | 9 | + | - | - | - | + |
| 2011 | 81/F | 8 | I | 12.5 | 94.67 | 7 | - | - | - | - | + |
| 2012 | 67/F | 16 | I | 14.2 | 94.5 | 5 | + | - | - | - | - |
| 2013 | 67/M | 19 | I | 57.8 | 93.96 | 8 | + | + | - | - | + |
| 2014 | 54/M | 12 | I | 41.75 | 95.13 | 3 | + | - | + | - | - |
| 2015 | 62/F | 12 | I | 28.45 | 95.16 | 6 | + | - | - | - | - |
| 2016 | 52/F | 12 | I | 21.8 | 94.35 | 10 | + | + | - | - | + |
| 2017 | 54/M | 10 | I | 35.5 | 95.12 | 6 | + | + | + | - | - |
| 2018 | 48/M | 12 | I | 21.19 | 94.59 | 5 | + | - | - | - | + |
| 2019 | 70/F | 12 | I | 1.89 | 94.17 | 9 | + | - | - | - | + |
| 2020 | 65/M | 12 | I | 123.19 | 94.68 | 6 | + | + | - | - | - |
| 2021 | 75/M | 11 | I | 52.12 | 93.17 | 7 | - | - | - | - | + |
| 2022 | 60/M | 16 | I | 4.02 | 90.42 | 4 | + | - | - | - | + |
| 2023 | 46/M | 11 | H | 11.61 | 94.25 | 6 | + | - | - | - | - |
| 2024* | 70/F | 15 | I | 49.61 | 92.42 | 11.5 | + | - | - | - | - |
| 2025 | 59/F | 10 | I | 8.45 | 94.86 | 8.6 | + | + | - | - | - |
| 2026 | 49/M | 14 | H | 9.13 | 93.57 | 5.6 | + | + | - | - | + |
| 2027 | 49/M | 9 | I | 12.29 | 93.86 | 8.4 | + | + | - | - | + |
| 2028 | 60/M | 12 | I | 3.25 | 94.38 | 6 | + | + | + | - | - |
| 2029 | 60/F | 12 | I | 0.84 | 91.08 | 10 | + | - | - | - | - |
| 2030 | 65/M | 16 | I | 11.82 | 94.19 | 10.6 | + | - | - | - | - |
| 2031 | 39/M | 14 | H | 21.72 | 95.57 | 13 | + | + | - | - | + |
| 2032 | 74/M | 15 | H | 55.91 | 93.16 | 8.2 | + | + | + | - | + |
| 2033 | 58/M | 12 | I | 0.41 | 94.32 | 6.4 | + | - | - | - | - |
| 2034 | 66/F | 8 | I | 2.95 | 93.86 | 4.6 | + | + | + | - | - |
| 2035 | 70/M | 8 | I | 25.98 | 95.17 | 2.4 | + | - | + | - | - |
| 2036 | 79/M | 10 | H | 42.17 | 94.09 | 2.3 | + | + | + | - | + |
| 2037 | 35/F | 12 | H | 8.78 | 94.52 | 9 | + | + | + | - | - |
| 2038 | 67/M | 12 | I | 3.28 | 94.1 | 8 | + | + | - | - | - |
| 2039 | 72/F | 10 | I | 1.18 | 94.2 | 20 | - | - | - | - | - |

* = left handedness (all other patients were right handers); I/H = ischemic/hemorrhagic stroke; CVST = cerebral venous sinus thrombosis; AVM = arterial venous malformation; TAO = time of memory testing after stroke onset; MI/SI = motor/sensory impairment (- = no impairment, + = mild impairment, ++ = moderate/severe impairment); VFD = visual field defect (- = no, -/e = extinction upon bilateral simultaneous stimulation but no VFD); NA = data not available.

**Table S2**. **Extent of damage (%) in brain regions defined by the AAL and WM atlases for individual RHD and LHD patients.**

**A. Right hemisphere damage (RHD) group**

| Areas | 1001 | 1002 | 1003 | 1004 | 1005 | 1006 | 1007 | 1008 | 1009 | 1010 | 1011 | 1012 | 1013 | 1014 | 1015 | 1016 | 1017 | 1018 | 1019 |
|---|---|---|---|---|---|---|---|---|---|---|---|---|---|---|---|---|---|---|---|
| Precentral g | 0 | 71.28 | 58.56 | 6.71 | 1.04 | 0.71 | 22.33 | 33.78 | 6.48 | 11.48 | 81.6 | 14.49 | 5.62 | 68.74 | 3.67 | 1.98 | 0.86 | 2.81 | 61.85 |
| Superior frontal g | 0 | 3.57 | 63.17 | 0.02 | 0 | 0 | 0 | 0.02 | 0 | 13.19 | 9.89 | 0 | 0 | 4.17 | 0 | 0 | 0 | 1.5 | 20.12 |
| Sup. frontal g, orbital | 0 | 28.89 | 38.01 | 1.5 | 0 | 0 | 0 | 19.26 | 0 | 20.46 | 0 | 0 | 0 | 56.17 | 0 | 0 | 1 | 2.11 | 0 |
| Mid. frontal g | 0 | 57.23 | 88.6 | 9.5 | 0 | 0 | 7.84 | 11.36 | 0 | 11.21 | 34.37 | 0 | 0.43 | 67.81 | 0.69 | 0 | 0.67 | 4.94 | 11.19 |
| Mid. frontal g, med. orb. | 0 | 70.94 | 59.41 | 0.1 | 0 | 0 | 0 | 62.27 | 0 | 7.98 | 0 | 0 | 0 | 81.58 | 0 | 0 | 0 | 1.67 | 0 |
| IFGOP | 2.64 | 90.42 | 22.94 | 76.13 | 0 | 3.57 | 48.96 | 88.21 | 6.22 | 20.23 | 90.85 | 0 | 25.88 | 99.07 | 34.6 | 1.86 | 40.17 | 51.47 | 0 |
| IFGTRI | 0.65 | 93.49 | 18.18 | 53.28 | 0 | 1.86 | 2.46 | 77.27 | 0 | 19.67 | 21.8 | 0 | 0.23 | 99.91 | 9.9 | 0 | 15.9 | 41.56 | 0 |
| IFGORB | 4.8 | 98.01 | 5.92 | 28.76 | 0 | 0 | 22.96 | 72.52 | 0 | 2.46 | 2.99 | 0 | 0 | 98.36 | 17.93 | 0 | 17.46 | 31.69 | 0 |
| Rolandic op. | 29.6 | 95.94 | 0 | 89.71 | 0 | 16.53 | 88.13 | 95.27 | 26.07 | 11.8 | 91.36 | 1.8 | 75.13 | 100 | 57.02 | 76.4 | 18.26 | 87.83 | 2.48 |
| Supp. motor area | 0 | 0 | 8.73 | 0 | 3.54 | 0 | 0 | 0 | 0 | 2.91 | 0 | 5.74 | 0 | 0 | 0 | 0 | 0 | 0 | 47.66 |
| Olfactory | 0 | 11.42 | 0 | 0.69 | 0 | 4.84 | 0 | 1.73 | 0 | 2.77 | 0 | 0 | 0 | 23.53 | 0 | 0 | 4.5 | 8.65 | 0 |
| Sup. frontal g, medial | 0 | 0 | 15.65 | 0 | 0 | 0 | 0 | 0 | 0 | 6.28 | 0 | 0 | 0 | 0.09 | 0 | 0 | 0 | 0.94 | 0 |
| Sup. frontal g, orbital | 0 | 0 | 10.63 | 0 | 0 | 0 | 0 | 1.29 | 0 | 26.64 | 0 | 0 | 0 | 8.18 | 0 | 0 | 0 | 0.47 | 0 |
| Rectus g | 0 | 2.68 | 0.13 | 0 | 0 | 0 | 0 | 0.13 | 0 | 2.28 | 0 | 0 | 0 | 24.3 | 0 | 0 | 0 | 2.01 | 0 |
| Insula | 58.42 | 99.89 | 1.07 | 98.53 | 0 | 17.68 | 63.5 | 99.32 | 3.95 | 6.89 | 79.32 | 1.53 | 61.53 | 100 | 85.08 | 41.5 | 69.89 | 95.76 | 0.23 |
| Anterior cingulum | 0 | 0.08 | 0 | 0 | 0 | 0 | 0 | 0 | 0 | 21.17 | 0 | 0 | 0 | 3.05 | 0 | 0 | 0 | 4.8 | 3.35 |
| Middle cingulum | 0 | 0.18 | 0.23 | 0 | 21.83 | 0 | 0 | 0 | 0 | 11.58 | 0 | 4.68 | 0.32 | 0 | 0 | 0.05 | 0 | 0 | 47.48 |
| Posterior cingulum | 0 | 0 | 0 | 0 | 0 | 0 | 0 | 0 | 0 | 0 | 0 | 0 | 2.69 | 0 | 0 | 0 | 0 | 0 | 0 |
| Hippocampus | 14.16 | 0 | 0 | 11.73 | 0 | 0.21 | 0.63 | 0.32 | 0 | 0 | 2.33 | 0 | 15.33 | 1.8 | 0 | 5.92 | 0.11 | 4.44 | 0 |
| Parahippocampal g | 0 | 1.24 | 0 | 0.71 | 0 | 0 | 4.59 | 0 | 0 | 0 | 0 | 0 | 0.18 | 6.54 | 0 | 0 | 0 | 0 | 0 |
| Amygdala | 3.23 | 6.05 | 0 | 13.31 | 0 | 26.21 | 5.65 | 5.65 | 0 | 0 | 5.24 | 0 | 1.21 | 37.9 | 0 | 0 | 13.71 | 7.26 | 0 |
| Calcarine | 0 | 0.7 | 0 | 0 | 0 | 0 | 0.05 | 0 | 0 | 0 | 0 | 0 | 2.85 | 0 | 0 | 0 | 0 | 0 | 0 |
| Cuneus | 0 | 0.07 | 0.14 | 0 | 0 | 0.63 | 0 | 0 | 0 | 0 | 1.4 | 0 | 0.21 | 0 | 0 | 0 | 0 | 0 | 0 |
| Lingual g | 0 | 0 | 0 | 0 | 0 | 0 | 0 | 0 | 0 | 0 | 0 | 0 | 2.3 | 0 | 0 | 0 | 0 | 0 | 0 |
| Superior occipital g | 1.06 | 0.07 | 1.77 | 0 | 0.28 | 1.77 | 0 | 0 | 0 | 0 | 8.7 | 0 | 1.49 | 0 | 0 | 0 | 0 | 0 | 0.57 |
| Middle occipital g | 7.29 | 14.87 | 10.1 | 0.24 | 0 | 5.2 | 2.48 | 0 | 0 | 1.76 | 5.05 | 0 | 3.24 | 27.55 | 0 | 0.71 | 0 | 0 | 0 |
| Inferior occipital g | 0 | 1.31 | 0 | 0 | 0 | 0 | 0 | 0 | 0 | 0 | 0 | 0 | 0 | 17.9 | 0 | 0 | 0 | 0 | 0 |
| Fusiform g | 0.08 | 1.07 | 0 | 4.01 | 0 | 0 | 3.02 | 0 | 0 | 0 | 0 | 0 | 2.14 | 2.1 | 0 | 0 | 0 | 0.28 | 0 |
| Postcentral g | 4.05 | 85.56 | 59.85 | 3.3 | 14.54 | 0.18 | 6.25 | 37.35 | 7.59 | 7.38 | 82.47 | 62.23 | 10.28 | 68.43 | 10.62 | 12.1 | 0 | 9.84 | 59.74 |
| Superior parietal g | 0.09 | 32.18 | 69.67 | 0 | 34.97 | 0 | 0 | 0 | 0 | 0.05 | 25.52 | 12.96 | 0 | 8.28 | 0.14 | 0 | 0 | 0 | 30.38 |
| Inferior parietal g | 20.52 | 96.88 | 86.02 | 0 | 7.06 | 0 | 0.22 | 2.97 | 0 | 1.26 | 94.35 | 17.55 | 16.21 | 64.76 | 22.45 | 0.22 | 0 | 18.74 | 1.41 |
| Supramarginal g | 34.8 | 96.61 | 15.4 | 10.28 | 0.51 | 12.82 | 61.96 | 84.14 | 2.33 | 17.98 | 96.1 | 18.24 | 73.91 | 98.18 | 68.69 | 63.6 | 0 | 59.02 | 3.55 |
| Angular g | 67.18 | 73.52 | 15.07 | 3.88 | 1.26 | 10.96 | 22.09 | 6.56 | 0 | 15.58 | 82.82 | 0 | 37.04 | 68.84 | 5.19 | 7.65 | 0 | 1.03 | 0.23 |
| Precuneus | 0.03 | 0.34 | 7.26 | 0 | 13.48 | 0 | 0 | 0 | 0 | 0 | 0.25 | 5.11 | 0.61 | 0 | 0 | 0 | 0 | 0.12 | 33.45 |
| Paracentral lobule | 0 | 0 | 37.32 | 0 | 20.22 | 0 | 0 | 0 | 0 | 1.2 | 0.24 | 52.87 | 0 | 0 | 0 | 0 | 0 | 0 | 80.38 |
| Caudate | 19.62 | 13.28 | 29.78 | 0.3 | 0 | 50.8 | 0 | 1.81 | 0 | 17 | 1.51 | 0 | 0 | 73.24 | 2.52 | 0 | 42.86 | 66.2 | 0.4 |
| Putamen | 56.77 | 68.7 | 47.09 | 30.36 | 0 | 84.87 | 21.05 | 75 | 0 | 25.56 | 24.53 | 0 | 10.62 | 100 | 55.92 | 12.4 | 92.11 | 93.14 | 7.71 |
| Pallidum | 25.36 | 52.86 | 4.29 | 2.86 | 0 | 76.43 | 11.43 | 19.64 | 0 | 71.43 | 0 | 0 | 8.57 | 98.57 | 4.64 | 1.07 | 64.64 | 48.57 | 0 |
| Thalamus | 17.6 | 0.95 | 0.09 | 1.51 | 0 | 7.28 | 0 | 0 | 0 | 0.28 | 0 | 0 | 0.38 | 12.68 | 0 | 0.95 | 2.27 | 4.26 | 0 |
| Heschl g | 69.48 | 98.39 | 0 | 90.36 | 0 | 6.83 | 93.98 | 96.39 | 0 | 2.41 | 97.19 | 0 | 94.78 | 100 | 71.49 | 90.8 | 2.81 | 97.99 | 0 |

| | | | | | | | | | | | | | | | | | | | |
|---|---|---|---|---|---|---|---|---|---|---|---|---|---|---|---|---|---|---|---|
| Superior temporal g | 69.15 | 91.02 | 0.38 | 76.06 | 0 | 8.28 | 94.08 | 54.95 | 0 | 5.44 | 59.57 | 0 | 80.74 | 99.87 | 22.83 | 45.2 | 2.42 | 60.9 | 0 |
| Superior temporal pole | 28.33 | 86.55 | 0 | 62.11 | 0 | 0 | 86.32 | 56.13 | 0 | 0 | 12.71 | 0 | 6.65 | 98.8 | 16.07 | 0.07 | 3.66 | 19.36 | 0 |
| Middle temporal g | 18.19 | 75.01 | 6.53 | 60.35 | 0 | 5.15 | 78.16 | 25.65 | 0 | 1.38 | 11.54 | 0 | 37.06 | 97.85 | 0.02 | 20.7 | 0 | 3.79 | 0 |
| Middle temporal pole | 4.63 | 82.14 | 0 | 37.41 | 0 | 0 | 86.6 | 34.12 | 0 | 0 | 0 | 0 | 0 | 95.45 | 0 | 0 | 0 | 0.25 | 0 |
| Inferior temporal g | 0.39 | 11.5 | 0 | 16.02 | 0 | 0 | 41.41 | 14.39 | 0 | 0 | 0 | 0 | 1.8 | 42.11 | 0 | 0.03 | 0 | 2.45 | 0 |
| CST | 0 | 0 | 0 | 0 | 0 | 0 | 0 | 0 | 0 | 0 | 0 | 0 | 0 | 0 | 0 | 0 | 0 | 0 | 0 |
| ML | 0 | 0 | 0 | 0 | 0 | 0 | 0 | 0 | 0 | 0 | 0 | 0 | 0 | 0 | 0 | 0 | 0 | 0 | 0 |
| ICP | 0 | 0 | 0 | 0 | 0 | 0 | 0 | 0 | 0 | 0 | 0 | 0 | 0 | 0 | 0 | 0 | 0 | 0 | 0 |
| SCP | 0 | 0 | 0 | 0 | 0 | 0 | 0 | 0 | 0 | 0 | 0 | 0 | 0 | 0 | 0 | 0 | 0 | 0 | 0 |
| CP | 14.93 | 0 | 0 | 0 | 0 | 0 | 1.12 | 0 | 0 | 0 | 0 | 0 | 1.12 | 0 | 0 | 0 | 0 | 0 | 0 |
| $IC_{AL}$ | 20.64 | 65.6 | 57 | 1.97 | 0 | 84.77 | 0 | 15.97 | 0 | 26.29 | 0 | 0 | 2.21 | 93.61 | 25.55 | 0 | 86.24 | 87.71 | 2.95 |
| $IC_{PL}$ | 77.84 | 33.93 | 30.74 | 19.56 | 0 | 64.67 | 0 | 19.36 | 0 | 29.74 | 3.19 | 1.6 | 12.57 | 78.44 | 14.77 | 13 | 53.09 | 58.48 | 0.4 |
| $IC_{RL}$ | 86.08 | 35.13 | 0 | 25.95 | 0 | 12.34 | 10.76 | 11.71 | 0 | 16.14 | 0.32 | 0 | 59.81 | 31.65 | 6.96 | 77.2 | 7.91 | 65.51 | 0 |
| $CR_{A}$ | 0 | 49.77 | 27.22 | 34.58 | 0 | 10.51 | 0 | 27.69 | 0 | 90.42 | 3.5 | 0 | 4.32 | 89.95 | 0.35 | 0 | 58.29 | 73.71 | 1.87 |
| $CR_{S}$ | 46.41 | 81.2 | 22.93 | 48.7 | 14.35 | 54.24 | 6.09 | 52.61 | 28.04 | 88.15 | 37.5 | 22.17 | 34.35 | 56.96 | 34.35 | 33.8 | 55.33 | 57.17 | 38.37 |
| $CR_{P}$ | 52.21 | 56.64 | 4.87 | 25.88 | 23.67 | 11.73 | 4.87 | 5.31 | 5.97 | 64.16 | 25 | 6.64 | 62.83 | 18.58 | 30.09 | 43.1 | 19.91 | 42.04 | 16.37 |
| $TR_{P}$ | 9.86 | 12.53 | 0 | 8.42 | 0 | 0 | 3.49 | 0 | 0 | 6.57 | 0 | 0 | 55.03 | 20.53 | 0 | 36.6 | 0 | 5.34 | 0 |
| SS | 52.45 | 11.19 | 0 | 21.68 | 0 | 0 | 15.03 | 2.45 | 0 | 0 | 0 | 0 | 53.85 | 17.83 | 0 | 34.6 | 0 | 16.43 | 0 |
| External capsule | 84.12 | 96.35 | 54.72 | 68.03 | 0 | 92.7 | 31.33 | 97.42 | 1.29 | 20.82 | 18.03 | 1.29 | 37.12 | 100 | 78.33 | 46.8 | 95.92 | 100 | 18.03 |
| CGC | 0 | 0 | 0 | 0 | 6.8 | 0 | 0 | 0 | 0 | 2.04 | 0 | 0 | 0.68 | 0 | 0 | 0 | 0 | 0 | 32.31 |
| CGH | 0 | 0 | 0 | 0 | 0 | 0 | 0 | 0 | 0 | 0 | 0 | 0 | 0 | 0 | 0 | 0 | 0 | 0 | 0 |
| FX/ST | 30.66 | 2.19 | 0 | 0 | 0 | 0 | 5.11 | 0 | 0 | 0 | 0 | 0 | 32.12 | 0 | 0 | 4.38 | 0 | 0.73 | 0 |
| SLF | 50.3 | 90.42 | 18.79 | 45.33 | 1.82 | 22.91 | 44.48 | 73.45 | 33.09 | 80.73 | 80.12 | 40.61 | 97.33 | 77.33 | 66.79 | 83.3 | 12.97 | 72 | 15.76 |
| $FO_{S}$ | 76.27 | 93.22 | 62.71 | 0 | 0 | 100 | 0 | 10.17 | 0 | 47.46 | 0 | 0 | 0 | 54.24 | 27.12 | 1.69 | 100 | 100 | 5.08 |
| $FO_{I}$ | 62.74 | 77.57 | 10.27 | 37.64 | 0 | 64.26 | 49.05 | 72.24 | 0 | 0.38 | 0 | 0 | 27.38 | 98.86 | 29.66 | 15.6 | 73.76 | 83.65 | 0 |
| UNC | 44.68 | 57.45 | 0 | 36.17 | 0 | 46.81 | 80.85 | 91.49 | 0 | 0 | 0 | 0 | 6.38 | 100 | 19.15 | 17 | 48.94 | 65.96 | 0 |
| TAP | 19.23 | 11.54 | 0 | 0 | 0 | 0 | 0 | 0 | 0 | 16.67 | 0 | 0 | 5.13 | 0 | 0 | 10.3 | 0 | 8.97 | 0 |
| PCT - bilateral | 0 | 0 | 0 | 0 | 0 | 0 | 0 | 0 | 0 | 0 | 0 | 0 | 0 | 0 | 0 | 0 | 0 | 0 | 0 |
| BCC - bilateral | 0 | 1.51 | 0.06 | 0 | 0.87 | 0.12 | 0 | 0 | 0 | 14.71 | 0 | 0.35 | 0 | 3.88 | 0.06 | 0.29 | 0.23 | 2.55 | 14.07 |
| FX - bilateral | 0 | 0 | 0 | 0 | 0 | 0 | 0 | 0 | 0 | 0 | 0 | 0 | 0 | 0 | 0 | 0 | 0 | 0 | 0 |
| MCP - bilateral | 0 | 0 | 0 | 0 | 0 | 0 | 0 | 0 | 0 | 0 | 0 | 0 | 0 | 0 | 0 | 0 | 0 | 0 | 0 |
| GCC - bilateral | 0 | 3.45 | 5.13 | 0 | 0 | 0 | 0 | 0 | 0 | 20.16 | 0 | 0 | 0 | 22.37 | 0 | 0 | 1.33 | 10.43 | 3.36 |
| SCC - bilateral | 0.19 | 1.88 | 0 | 0 | 0 | 0 | 0 | 0 | 0 | 0.06 | 0 | 0 | 4.28 | 0 | 0 | 0.06 | 0 | 2.59 | 0 |

**A. Right hemisphere damage (RHD) group (cont.)**

| Areas | 1020 | 1021 | 1022 | 1023 | 1024 | 1025 | 1026 | 1027 | 1028 | 1029 | 1030 | 1031 | 1032 | 1033 | 1034 | 1035 | 1036 | 1037 | 1038 |
|---|---|---|---|---|---|---|---|---|---|---|---|---|---|---|---|---|---|---|---|
| Precentral g | 7.93 | 0 | 14.23 | 0.12 | 37.62 | 22.89 | 0 | 39.6 | 14.2 | 19.4 | 7.84 | 11.5 | 4.02 | 35.08 | 0.06 | 16.95 | 0 | 14.73 | 0 |
| Superior frontal g | 0.02 | 0 | 0.59 | 0.02 | 24.85 | 9.32 | 0 | 16.96 | 3.55 | 0.2 | 5.7 | 0 | 0 | 0.52 | 0 | 0 | 0 | 12.15 | 0 |
| Sup. frontal g, orbital | 0 | 0 | 4.81 | 0.1 | 14.64 | 4.31 | 0 | 0 | 0 | 0.7 | 0.3 | 0 | 0 | 0 | 0 | 0 | 0 | 0 | 0 |
| Mid. frontal g | 2.04 | 0 | 5.25 | 1.06 | 57.48 | 5.33 | 0 | 3.12 | 0.45 | 10.68 | 4.27 | 2.02 | 4.68 | 33.99 | 0.02 | 18.34 | 0 | 2.16 | 0 |
| Mid. frontal g, med. orb. | 0 | 0 | 8.57 | 0.89 | 40.49 | 2.66 | 0 | 0 | 0 | 12.91 | 0 | 0 | 0 | 1.97 | 0 | 0.1 | 0 | 0 | 0 |
| $IFG_{OP}$ | 44.89 | 0 | 70.34 | 10.01 | 13.8 | 23.02 | 6.58 | 2.07 | 8.72 | 77.06 | 17.23 | 16.9 | 11.58 | 95.57 | 0 | 93.14 | 0.21 | 0 | 0 |
| $IFG_{TRI}$ | 11.34 | 0 | 51.93 | 11.11 | 25.71 | 15.67 | 1.02 | 0.65 | 0 | 70.2 | 7.58 | 3.63 | 18.55 | 79.36 | 0 | 81.08 | 0.14 | 0 | 0 |
| $IFG_{ORB}$ | 12.6 | 0.06 | 38.72 | 10.37 | 36.85 | 3.34 | 12.36 | 16.93 | 0 | 36.26 | 0.47 | 0.18 | 0 | 19.27 | 0.88 | 10.95 | 0.94 | 0 | 0 |

| | | | | | | | | | | | | | | | | | | | |
|---|---|---|---|---|---|---|---|---|---|---|---|---|---|---|---|---|---|---|---|
| Rolandic op. | 89.93 | 1.43 | 77.76 | 75.36 | 1.73 | 40.57 | 8.26 | 54.32 | 12.47 | 92.11 | 28.32 | 14.4 | 9.24 | 93.91 | 0 | 83.02 | 11.87 | 0 | 0 |
| Supp. motor area | 0 | 0 | 0 | 0 | 5.19 | 21.81 | 0 | 60.35 | 16.28 | 0 | 3.29 | 0 | 0 | 0 | 0 | 0 | 0 | 32.77 | 0 |
| Olfactory | 3.46 | 0 | 0 | 0 | 0 | 2.42 | 2.08 | 0 | 0 | 5.19 | 0 | 0 | 9.34 | 0 | 0 | 0.35 | 0 | 0 | 0 |
| Sup. frontal g, medial | 0 | 0 | 0 | 0 | 0.19 | 10.45 | 0 | 0.7 | 0 | 0 | 1.31 | 0 | 0 | 0 | 0 | 0 | 0 | 0 | 0 |
| Sup. frontal g, orbital | 0 | 0 | 0 | 0 | 0.35 | 16.82 | 0 | 0 | 0 | 0 | 0 | 0 | 0 | 0 | 0 | 0 | 0 | 0 | 0 |
| Rectus g | 0 | 0 | 0 | 0 | 1.07 | 5.91 | 0 | 0 | 0 | 0 | 0 | 0 | 0.27 | 0 | 0 | 0 | 0 | 0 | 0 |
| Insula | 86.95 | 20.96 | 74.12 | 93.95 | 12.37 | 49.77 | 68.14 | 36.5 | 1.24 | 68.59 | 57.8 | 8.47 | 11.47 | 85.93 | 0 | 86.16 | 48.31 | 0 | 0 |
| Anterior cingulum | 0 | 0 | 0 | 0.08 | 0.99 | 23.08 | 0 | 2.28 | 0 | 0 | 3.27 | 0 | 0 | 0 | 0 | 0 | 0 | 0 | 0 |
| Middle cingulum | 0.32 | 0 | 1.04 | 0 | 6.08 | 14.25 | 0 | 24.88 | 10.17 | 0.41 | 3.45 | 0 | 0 | 0 | 0 | 0 | 0 | 8.76 | 0 |
| Posterior cingulum | 0 | 0 | 0 | 0 | 0 | 0 | 0 | 0 | 42.09 | 0 | 0 | 0 | 0.3 | 0 | 0 | 0 | 0 | 0 | 0 |
| Hippocampus | 15.01 | 0 | 0 | 3.07 | 0 | 14.38 | 50.95 | 0 | 1.9 | 20.93 | 21.04 | 0 | 0 | 0.63 | 0 | 0 | 2.11 | 0 | 0 |
| Parahippocampal g | 3.45 | 0 | 0 | 0 | 1.5 | 0.8 | 3.8 | 0 | 0 | 10.42 | 0 | 0 | 0 | 14.66 | 0 | 0 | 0.09 | 0 | 0 |
| Amygdala | 25.4 | 0 | 0 | 10.48 | 0 | 1.21 | 30.24 | 0 | 0 | 58.06 | 11.29 | 0 | 7.66 | 73.39 | 0 | 4.03 | 4.44 | 0 | 0 |
| Calcarine | 0.21 | 0 | 0 | 0 | 0 | 0 | 0 | 0 | 66.79 | 0 | 6.23 | 6.61 | 0 | 0 | 0 | 0 | 0 | 0 | 0 |
| Cuneus | 0 | 0 | 0 | 0 | 1.69 | 0 | 0 | 0 | 78.93 | 0 | 3.79 | 3.02 | 0.63 | 0 | 0 | 0 | 0 | 0 | 0 |
| Lingual g | 0 | 0 | 0 | 0 | 0 | 0 | 0 | 0 | 5.83 | 9.96 | 0 | 0.13 | 0 | 0 | 0 | 0 | 0 | 0 | 0 |
| Superior occipital g | 0 | 0 | 0 | 0 | 4.25 | 0 | 0 | 0 | 68.22 | 0.71 | 6.51 | 19.9 | 0.42 | 0 | 0.21 | 0 | 0 | 0 | 0 |
| Middle occipital g | 4.1 | 0 | 0 | 0 | 3.43 | 0 | 0 | 0 | 71.16 | 5.96 | 8.1 | 66 | 0.14 | 0 | 2.43 | 0 | 0 | 0 | 0 |
| Inferior occipital g | 0 | 0 | 0 | 0 | 0 | 0 | 0 | 0 | 23.86 | 18.71 | 0 | 8.9 | 0 | 0 | 0 | 0 | 0 | 0 | 0 |
| Fusiform g | 1.47 | 0 | 0 | 0 | 0 | 0.12 | 0.56 | 0 | 5.08 | 22.52 | 1.07 | 1.19 | 0 | 3.85 | 0 | 0 | 0.2 | 0 | 0 |
| Postcentral g | 7.64 | 0 | 3.48 | 0.94 | 47.58 | 22.31 | 0 | 82.5 | 25.77 | 38.29 | 10.75 | 1.36 | 2.01 | 37.82 | 6.51 | 3.24 | 0.24 | 2.85 | 0 |
| Superior parietal g | 0 | 0 | 0 | 0 | 41.49 | 0.14 | 0 | 48.83 | 41.09 | 11.39 | 0.05 | 8.64 | 0.68 | 0.05 | 22.37 | 0 | 0 | 0 | 0 |
| Inferior parietal g | 6.25 | 0 | 0 | 0 | 80.15 | 6.25 | 0.22 | 92.57 | 58.96 | 85.72 | 4.76 | 49.5 | 17.17 | 42.83 | 63.64 | 0 | 0 | 0 | 0 |
| Supramarginal g | 39.31 | 2.74 | 2.53 | 8.66 | 24.62 | 22.14 | 15.25 | 92.2 | 53.29 | 73.86 | 29.74 | 54.7 | 8.41 | 80.24 | 29.23 | 1.67 | 0 | 0 | 0 |
| Angular g | 29.11 | 8.28 | 0 | 0 | 51.83 | 3.54 | 1.37 | 29.05 | 89.67 | 62.39 | 18.44 | 94.6 | 13.3 | 0.46 | 34.02 | 0 | 0 | 0 | 0 |
| Precuneus | 0 | 0 | 0 | 0 | 15.16 | 1.68 | 0 | 12.25 | 53.54 | 0 | 3.83 | 1.56 | 2.7 | 0 | 0.28 | 0 | 0 | 0.12 | 0 |
| Paracentral lobule | 0 | 0 | 0 | 0 | 23.09 | 22.61 | 0 | 70.33 | 14.95 | 0.12 | 3.95 | 0 | 0 | 0 | 0 | 0 | 0 | 44.5 | 0 |
| Caudate | 51.91 | 0 | 0.1 | 4.12 | 0.4 | 22.23 | 5.03 | 0 | 0 | 28.97 | 21.93 | 0 | 8.35 | 14.89 | 0 | 1.01 | 61.37 | 0 | 0 |
| Putamen | 61.84 | 23.4 | 6.58 | 30.92 | 12.22 | 50.94 | 73.87 | 0.75 | 0 | 82.71 | 59.87 | 0.47 | 77.63 | 60.81 | 0 | 37.97 | 67.39 | 0 | 0 |
| Pallidum | 1.07 | 13.21 | 0 | 5 | 0 | 39.64 | 41.07 | 0 | 0 | 66.79 | 18.57 | 0 | 19.29 | 14.29 | 0 | 4.29 | 30.71 | 0 | 1.43 |
| Thalamus | 4.45 | 6.15 | 0 | 0 | 0 | 29.61 | 16.84 | 0 | 0.09 | 8.61 | 30.37 | 0 | 0.19 | 0 | 0 | 0.09 | 2.08 | 0 | 1.23 |
| Heschl g | 97.99 | 55.02 | 35.74 | 83.94 | 2.81 | 44.58 | 65.06 | 83.94 | 12.45 | 97.99 | 43.78 | 64.7 | 9.64 | 80.72 | 0 | 85.14 | 26.51 | 0 | 0 |
| Superior temporal g | 80.58 | 58.61 | 10.28 | 14.39 | 3.92 | 5.09 | 67.14 | 55.81 | 15.19 | 68.13 | 15.5 | 82.9 | 3.12 | 81.41 | 17.67 | 63.9 | 2.74 | 0 | 0 |
| Superior temporal pole | 52.77 | 21.75 | 6.28 | 17.79 | 6.8 | 0.15 | 58.37 | 26.91 | 0 | 17.64 | 2.84 | 2.09 | 0 | 45.52 | 9.27 | 29.45 | 0.22 | 0 | 0 |
| Middle temporal g | 54.21 | 2.79 | 0 | 0 | 8.69 | 5.15 | 7.08 | 25.49 | 25.49 | 47.33 | 21.52 | 66.6 | 3.67 | 17.31 | 25.47 | 34.23 | 0 | 0 | 0 |
| Middle temporal pole | 18.28 | 6.15 | 0 | 0 | 0 | 0 | 41.53 | 0.93 | 0 | 8.51 | 1.01 | 0 | 0 | 54.42 | 0 | 1.35 | 0 | 0 | 0 |
| Inferior temporal g | 3.96 | 0 | 0 | 0 | 0 | 1.04 | 2.7 | 0.31 | 2.42 | 1.69 | 7.08 | 9.81 | 0 | 5.54 | 12.85 | 0 | 0.06 | 0 | 0 |
| CST | 0 | 0 | 0 | 0 | 0 | 0 | 0 | 0 | 0 | 0 | 0 | 0 | 0 | 0 | 0 | 0 | 0 | 0 | 0 |
| ML | 0 | 0 | 0 | 0 | 0 | 0 | 0 | 0 | 0 | 0 | 0 | 0 | 0 | 0 | 0 | 0 | 0 | 0 | 0 |
| ICP | 0 | 0 | 0 | 0 | 0 | 0 | 0 | 0 | 0 | 0 | 0 | 0 | 0 | 0 | 0 | 0 | 0 | 0 | 0 |
| SCP | 0 | 0 | 0 | 0 | 0 | 0 | 0 | 0 | 0 | 0 | 0 | 0 | 0 | 0 | 0 | 0 | 0 | 0 | 0 |
| CP | 0 | 0 | 0 | 0 | 0 | 41.04 | 29.1 | 0 | 0 | 0.75 | 1.87 | 0 | 0 | 0 | 0 | 0 | 0 | 0 | 8.21 |
| $IC_{AL}$ | 62.16 | 0 | 0 | 1.72 | 1.72 | 58.23 | 19.41 | 0 | 0 | 73.71 | 47.67 | 0 | 29.73 | 43.73 | 0 | 3.44 | 81.57 | 0 | 0 |
| $IC_{PL}$ | 37.92 | 36.73 | 0 | 3.39 | 11.58 | 83.43 | 77.84 | 0 | 0 | 62.28 | 72.06 | 0 | 21.16 | 1.6 | 0 | 32.34 | 32.93 | 0 | 5.39 |
| $IC_{RL}$ | 64.87 | 34.81 | 0 | 0.32 | 0 | 82.59 | 85.44 | 0 | 0.63 | 71.2 | 92.41 | 0 | 18.99 | 0 | 0 | 11.08 | 0.63 | 0 | 0 |

| | | | | | | | | | | | | | | | | | | | |
|---|---|---|---|---|---|---|---|---|---|---|---|---|---|---|---|---|---|---|---|
| $CR_A$ | 47.9 | 0 | 31.89 | 44.04 | 72.66 | 98.01 | 3.27 | 3.27 | 0 | 52.8 | 86.1 | 0 | 18.69 | 32.71 | 0 | 36.45 | 5.96 | 0 | 0 |
| $CR_S$ | 88.15 | 1.09 | 33.8 | 25 | 66.41 | 91.52 | 48.04 | 12.39 | 44.35 | 85 | 96.52 | 0 | 59.89 | 40.33 | 0 | 60.33 | 20.22 | 17.61 | 0 |
| $CR_P$ | 60.4 | 10.4 | 0.22 | 10.18 | 27.21 | 50 | 47.79 | 1.55 | 76.55 | 65.27 | 85.18 | 42.7 | 58.41 | 0 | 0 | 10.4 | 5.97 | 0 | 0 |
| $TR_P$ | 64.48 | 3.49 | 0 | 0 | 3.49 | 6.37 | 9.24 | 0 | 64.89 | 26.49 | 68.99 | 53.2 | 15.81 | 0 | 0 | 12.11 | 0 | 0 | 0 |
| SS | 66.08 | 0.7 | 0 | 0 | 0 | 41.96 | 56.99 | 0 | 0 | 74.13 | 66.78 | 7.69 | 0 | 0 | 0 | 0.7 | 0.7 | 0 | 0 |
| External capsule | 96.14 | 44.85 | 12.88 | 65.67 | 31.12 | 60.73 | 96.14 | 0.86 | 0.21 | 99.79 | 87.12 | 0.64 | 87.98 | 78.76 | 0 | 69.53 | 84.12 | 0 | 0 |
| CGC | 0 | 0 | 0 | 0 | 0.68 | 10.54 | 0 | 0 | 17.69 | 0 | 0 | 0 | 0 | 0 | 0 | 0 | 0 | 0 | 0 |
| CGH | 0 | 0 | 0 | 0 | 0 | 1.31 | 0 | 0 | 5.23 | 0 | 0 | 0 | 0 | 0 | 0 | 0 | 0 | 0 | 0 |
| FX/ST | 4.38 | 0 | 0 | 2.19 | 0 | 35.77 | 78.1 | 0 | 0 | 27.74 | 51.82 | 0 | 0 | 0 | 0 | 0 | 0.73 | 0 | 0 |
| SLF | 94.91 | 7.15 | 25.7 | 34.42 | 75.15 | 76.48 | 48.61 | 53.21 | 94.55 | 99.76 | 100 | 36.7 | 94.06 | 51.39 | 0 | 48.48 | 1.45 | 1.82 | 0 |
| $FO_S$ | 100 | 0 | 6.78 | 38.98 | 0 | 74.58 | 27.12 | 0 | 0 | 100 | 84.75 | 0 | 11.86 | 42.37 | 0 | 33.9 | 94.92 | 0 | 0 |
| $FO_I$ | 73 | 30.8 | 1.52 | 31.94 | 1.14 | 57.41 | 80.23 | 4.56 | 0 | 92.02 | 78.71 | 3.8 | 67.3 | 54.75 | 0 | 42.59 | 47.91 | 0 | 0 |
| UNC | 100 | 14.89 | 4.26 | 55.32 | 0 | 34.04 | 80.85 | 0 | 0 | 97.87 | 74.47 | 0 | 42.55 | 100 | 0 | 44.68 | 61.7 | 0 | 0 |
| TAP | 30.77 | 1.28 | 0 | 0 | 0 | 14.1 | 6.41 | 0 | 84.62 | 19.23 | 30.77 | 25.6 | 35.9 | 0 | 0 | 3.85 | 0 | 0 | 0 |
| PCT - bilateral | 0 | 0 | 0 | 0 | 0 | 0 | 0 | 0 | 0 | 0 | 0 | 0 | 0 | 0 | 0 | 0 | 0 | 0 | 0 |
| BCC - bilateral | 2.08 | 0 | 0.52 | 1.1 | 1.51 | 15.11 | 0.06 | 0.17 | 4.86 | 2.03 | 10.65 | 0 | 4.69 | 0 | 0 | 0.17 | 0 | 0.98 | 0 |
| FX - bilateral | 0 | 0 | 0 | 0 | 0 | 0 | 0 | 0 | 0 | 0 | 0 | 0 | 0 | 0 | 0 | 0 | 0 | 0 | 0 |
| MCP - bilateral | 0 | 0 | 0 | 0 | 0 | 0 | 0 | 0 | 0 | 0 | 0 | 0 | 0 | 0 | 0 | 0 | 0 | 0 | 0 |
| GCC - bilateral | 1.33 | 0 | 0.44 | 2.21 | 2.03 | 15.56 | 0 | 0 | 0 | 1.15 | 6.37 | 0 | 0.62 | 0 | 0 | 0.35 | 0.09 | 0 | 0 |
| SCC - bilateral | 0.13 | 0 | 0 | 0 | 0.06 | 0.39 | 0.19 | 0 | 32.34 | 0 | 3.11 | 2.27 | 4.41 | 0 | 0 | 0 | 0 | 0 | 0 |

### A. Right hemisphere damage (RHD) group (cont.)

| Areas | 1039 | 1040 | 1041 | 1042 | 1043 | 1044 | 1045 | 1046 | 1047 | 1048 | 1049 | 1050 | 1051 | 1052 | 1053 | 1054 | **Average** |
|---|---|---|---|---|---|---|---|---|---|---|---|---|---|---|---|---|---|
| Precentral g | 0.21 | 1.27 | 0 | 0 | 0 | 0.59 | 55.58 | 0 | 0.03 | 0 | 0 | 0 | 17.98 | 4.79 | 61.37 | 0 | **15.59** |
| Superior frontal g | 0.12 | 0 | 0 | 0 | 0 | 0 | 10.9 | 0 | 0 | 0 | 0 | 0 | 0.2 | 0.02 | 7.62 | 0 | **3.86** |
| Sup. frontal g, orbital | 0 | 0 | 0 | 0 | 0 | 0 | 2.61 | 0 | 0.2 | 0 | 0 | 0 | 2.61 | 4.41 | 21.56 | 0 | **4.14** |
| Mid. frontal g | 0.24 | 0 | 0 | 0 | 0 | 0 | 42.36 | 0 | 0.59 | 0 | 0 | 0 | 21.96 | 0.53 | 62.42 | 0 | **10.83** |
| Mid. frontal g, orbital | 0 | 0 | 0 | 0 | 0 | 0 | 3.35 | 0 | 0 | 0 | 0 | 0 | 20.1 | 0 | 52.81 | 0 | **7.92** |
| $IFG_{OP}$ | 6.58 | 0 | 0 | 0 | 0 | 0 | 76.84 | 0 | 20.66 | 0.07 | 0 | 0 | 80.27 | 9.86 | 86.78 | 0 | **27.32** |
| $IFG_{TRI}$ | 2.09 | 0 | 0 | 0 | 0 | 0 | 54.67 | 0 | 27.15 | 0 | 0 | 0 | 70.57 | 6.46 | 72.76 | 0 | **19.77** |
| $IFG_{ORB}$ | 0 | 0 | 0 | 0 | 0 | 0 | 28.59 | 0 | 20.33 | 0 | 0 | 0 | 26.54 | 3.22 | 48.45 | 0 | **13.54** |
| Rolandic op. | 7.66 | 0 | 0 | 0.15 | 0.3 | 25.32 | 57.33 | 9.24 | 25.62 | 3.83 | 0 | 0 | 50.56 | 7.96 | 93.61 | 0 | **35.92** |
| Supp. motor area | 0 | 2.36 | 0 | 0 | 0 | 0 | 13.79 | 0 | 0 | 0 | 0 | 0 | 0 | 0 | 0 | 0 | **4.16** |
| Olfactory | 0 | 0 | 0 | 0 | 0 | 0 | 13.15 | 0 | 2.08 | 0 | 0 | 0 | 0.35 | 5.54 | 0 | 0 | **1.89** |
| Sup. frontal g, medial | 0 | 0 | 0 | 0 | 0 | 0 | 0 | 0 | 0 | 0 | 0 | 0 | 0 | 0.14 | 0.09 | 0 | **0.66** |
| Sup. frontal g, med. orb. | 0 | 0 | 0 | 0 | 0 | 0 | 0 | 0 | 0 | 0 | 0 | 0 | 3.39 | 2.45 | 0 | 0 | **1.30** |
| Rectus g | 0 | 0 | 0 | 0 | 0 | 0 | 0.94 | 0 | 0.13 | 0 | 0 | 0 | 0 | 6.44 | 0.67 | 0 | **0.87** |
| Insula | 13.62 | 0 | 0.34 | 4.41 | 4.29 | 0 | 95.54 | 10.51 | 88.47 | 38.25 | 0 | 0 | 65.99 | 65.37 | 93.39 | 2.88 | **42.18** |
| Anterior cingulum | 0 | 0 | 0 | 0 | 0 | 0 | 4.65 | 0 | 0 | 0 | 0 | 0 | 1.6 | 0 | 0 | 0 | **1.27** |
| Middle cingulum | 0 | 11.03 | 0 | 0 | 0 | 0.27 | 10.99 | 0 | 0 | 0 | 0 | 0 | 0 | 0 | 0.64 | 0 | **3.31** |
| Posterior cingulum | 0 | 1.19 | 0 | 0 | 0 | 0 | 0 | 0 | 0 | 0 | 0 | 0 | 0 | 0 | 0 | 0 | **0.86** |
| Hippocampus | 8.03 | 0 | 0 | 0 | 0 | 0 | 42.6 | 0 | 5.18 | 0 | 0 | 0 | 0 | 21.67 | 0 | 0 | **4.90** |
| Parahippocampal g | 0.18 | 0 | 0 | 0 | 0 | 0 | 6.54 | 0 | 0 | 0 | 0 | 0 | 0 | 0.09 | 0 | 0 | **1.01** |

| | | | | | | | | | | | | | | | | | |
|---|---|---|---|---|---|---|---|---|---|---|---|---|---|---|---|---|---|
| Amygdala | 1.61 | 0 | 0 | 0 | 0 | 0 | 81.45 | 0 | 38.71 | 2.82 | 0 | 0 | 0 | 17.74 | 0 | 0 | **9.15** |
| Calcarine | 4.41 | 0 | 0 | 0 | 0 | 0 | 0 | 0 | 0 | 0 | 0 | 0 | 0 | 0 | 1.99 | 0 | **1.66** |
| Cuneus | 0.28 | 0 | 0 | 0 | 0 | 0 | 0 | 0 | 0 | 0 | 0 | 0 | 0 | 0 | 2.46 | 0 | **1.73** |
| Lingual g | 3.35 | 0 | 0 | 0 | 0 | 0 | 0 | 0 | 0 | 0 | 0 | 0 | 0 | 0 | 0.04 | 0 | **0.40** |
| Superior occipital g | 0.99 | 0 | 0 | 0 | 0 | 0 | 0 | 0 | 0 | 0 | 0 | 0 | 0 | 0 | 18.26 | 0 | **2.50** |
| Middle occipital g | 20.69 | 0 | 0 | 0 | 0 | 0 | 0 | 0 | 0 | 0 | 0 | 0 | 0 | 0 | 54.29 | 0 | **5.83** |
| Inferior occipital g | 11.83 | 0 | 0 | 0 | 0 | 0 | 0 | 0 | 0 | 0 | 0 | 0 | 0 | 0 | 0 | 0 | **1.53** |
| Fusiform g | 5.24 | 0 | 0 | 0 | 0 | 0 | 0.48 | 0 | 0 | 0 | 0 | 0 | 0 | 3.85 | 0 | 0 | **1.08** |
| Postcentral g | 0.08 | 6.25 | 0 | 0 | 0 | 30.32 | 19.59 | 1.96 | 0 | 0 | 0 | 0 | 2.33 | 4.19 | 69.34 | 0 | **17.95** |
| Superior parietal g | 0 | 4.28 | 0 | 0 | 0 | 11.79 | 0 | 0 | 0 | 0 | 0 | 0 | 0 | 0 | 54.28 | 0 | **8.51** |
| Inferior parietal g | 0 | 0.52 | 0 | 0 | 0 | 33.31 | 0.82 | 0 | 0 | 0 | 0 | 0 | 0 | 0 | 93.31 | 0 | **20.12** |
| Supramarginal g | 6.33 | 0.35 | 0 | 0 | 0 | 49.54 | 1.93 | 2.28 | 0 | 0 | 0 | 0 | 0.05 | 2.99 | 96.3 | 0 | **28.08** |
| Angular g | 18.78 | 0 | 0 | 0 | 0 | 4.22 | 0 | 0 | 0 | 0 | 0 | 0 | 0 | 1.48 | 96.92 | 0 | **18.08** |
| Precuneus | 0.37 | 11.85 | 0 | 0 | 0 | 1.07 | 0.28 | 0 | 0 | 0 | 0 | 0 | 0 | 0 | 4.53 | 0 | **3.15** |
| Paracentral lobule | 0 | 18.9 | 0 | 0 | 0 | 2.03 | 36.12 | 0 | 0 | 0 | 0 | 0 | 0 | 0 | 0 | 0 | **7.94** |
| Caudate | 2.21 | 0 | 0 | 1.21 | 0 | 0 | 43.96 | 0.4 | 16.4 | 5.43 | 1.91 | 0 | 4.53 | 24.55 | 3.02 | 10.56 | **12.11** |
| Putamen | 20.39 | 0 | 0 | 6.48 | 8.93 | 0 | 85.71 | 19.92 | 81.48 | 80.08 | 10.43 | 11.47 | 44.92 | 70.21 | 31.48 | 6.58 | **35.76** |
| Pallidum | 10 | 0 | 0 | 13.21 | 2.86 | 0 | 79.64 | 19.64 | 88.57 | 48.57 | 1.07 | 3.57 | 0 | 11.07 | 0 | 0 | **18.96** |
| Thalamus | 2.84 | 0 | 0 | 6.34 | 5.2 | 0 | 31.6 | 7.19 | 5.49 | 1.99 | 0.28 | 0 | 0 | 16.46 | 0.09 | 0.85 | **4.19** |
| Heschl g | 14.06 | 0 | 0 | 2.01 | 0.4 | 0 | 42.17 | 10.04 | 15.66 | 16.87 | 0 | 0 | 9.24 | 34.14 | 98.39 | 0 | **39.74** |
| Superior temporal g | 55.43 | 0 | 0 | 0 | 0 | 5.51 | 24.67 | 2.99 | 8.47 | 0.19 | 0 | 0 | 3.09 | 8.79 | 64.79 | 0 | **29.47** |
| Superior temporal pole | 0.9 | 0 | 0 | 0 | 0 | 0 | 64.65 | 0 | 29.82 | 0 | 0 | 0 | 1.79 | 1.2 | 5.08 | 0 | **16.26** |
| Middle temporal g | 63.01 | 0 | 0 | 0 | 0 | 0 | 7.6 | 0 | 0.07 | 0 | 0 | 0 | 0 | 8.1 | 24.04 | 0 | **16.47** |
| Middle temporal pole | 1.94 | 0 | 0 | 0 | 0 | 0 | 11.88 | 0 | 0.25 | 0 | 0 | 0 | 0 | 2.36 | 0 | 0 | **9.06** |
| Inferior temporal g | 25.7 | 0 | 0 | 0 | 0 | 0 | 2.31 | 0 | 0 | 0 | 0 | 0 | 0 | 9.08 | 0 | 0 | **3.98** |
| CST | 0 | 0 | 0 | 0 | 0 | 0 | 0 | 0 | 0 | 0 | 0 | 0 | 0 | 0 | 0 | 0 | **0.00** |
| ML | 0 | 0 | 0 | 0 | 0 | 0 | 0 | 0 | 0 | 0 | 0 | 0 | 0 | 0 | 0 | 0 | **0.00** |
| ICP | 0 | 0 | 0 | 0 | 0 | 0 | 0 | 0 | 0 | 0 | 0 | 0 | 0 | 0 | 0 | 0 | **0.00** |
| SCP | 0 | 0 | 0 | 0 | 0 | 0 | 0 | 0 | 0 | 0 | 0 | 0 | 0 | 0 | 0 | 0 | **0.00** |
| CP | 0 | 0 | 0 | 0 | 0 | 0 | 60.07 | 2.99 | 0.37 | 0 | 0 | 0 | 0 | 4.48 | 0 | 0 | **3.08** |
| IC$_{AL}$ | 2.7 | 0 | 0 | 0.25 | 3.19 | 0 | 71.01 | 0 | 51.35 | 31.94 | 16.71 | 0 | 4.42 | 26.54 | 1.23 | 27.76 | **22.79** |
| IC$_{PL}$ | 34.33 | 0 | 0 | 56.29 | 45.51 | 0 | 90.82 | 44.31 | 48.5 | 52.3 | 17.56 | 0.2 | 0 | 51.9 | 1.8 | 31.34 | **27.13** |
| IC$_{RL}$ | 52.53 | 0 | 0 | 11.71 | 3.16 | 0 | 74.68 | 27.53 | 4.43 | 31.01 | 0 | 0 | 0 | 83.23 | 47.78 | 2.53 | **23.13** |
| CR$_{A}$ | 23.71 | 0 | 0 | 0 | 0 | 0 | 67.52 | 0 | 27.45 | 3.62 | 0.82 | 0 | 60.63 | 54.67 | 65.3 | 0 | **24.25** |
| CR$_{S}$ | 34.35 | 3.26 | 3.15 | 11.85 | 1.3 | 0 | 96.09 | 34.02 | 37.5 | 23.26 | 4.13 | 0 | 3.59 | 54.02 | 75.54 | 23.91 | **37.21** |
| CR$_{P}$ | 26.99 | 47.57 | 0 | 1.77 | 0 | 0 | 46.24 | 20.58 | 0.44 | 2.88 | 0 | 0 | 0 | 27.65 | 62.83 | 6.64 | **24.36** |
| TR$_{P}$ | 66.53 | 0 | 0 | 0 | 0 | 0 | 2.05 | 0 | 0 | 0 | 0 | 0 | 0 | 15.81 | 28.54 | 0 | **11.11** |
| SS | 40.21 | 0 | 0 | 0 | 17.17 | 0 | 47.9 | 0 | 5.24 | 1.4 | 0 | 0 | 0 | 52.8 | 1.4 | 0 | **13.10** |
| External capsule | 42.49 | 0 | 1.5 | 20.17 | 0 | 0 | 97.42 | 40.77 | 89.06 | 97.64 | 3.65 | 5.79 | 66.31 | 98.5 | 76.82 | 25.75 | **48.61** |
| CGC | 0 | 0 | 0 | 0 | 0 | 0 | 14.29 | 0 | 0 | 0 | 0 | 0 | 0 | 0 | 0 | 0 | **1.57** |
| CGH | 0 | 0 | 0 | 0 | 0 | 0 | 4.58 | 0 | 0 | 0 | 0 | 0 | 0 | 0 | 0 | 0 | **0.21** |
| FX/ST | 11.68 | 0 | 0 | 0 | 0 | 0 | 51.82 | 0 | 9.49 | 1.46 | 0 | 0 | 0 | 42.34 | 0 | 0 | **7.27** |
| SLF | 39.03 | 5.94 | 1.21 | 0.73 | 0 | 45.94 | 56.12 | 18.06 | 15.64 | 8.12 | 0 | 0 | 10.06 | 77.94 | 93.94 | 0.12 | **43.04** |
| FO$_{S}$ | 6.78 | 0 | 3.39 | 1.69 | 0 | 0 | 100 | 1.69 | 55.93 | 55.93 | 0 | 0 | 0 | 100 | 8.47 | 47.46 | **31.01** |
| FO$_{I}$ | 25.1 | 0 | 0 | 0 | 0 | 0 | 92.4 | 1.52 | 74.14 | 62.36 | 0 | 0 | 39.92 | 84.41 | 31.94 | 0 | **33.01** |

| | | | | | | | | | | | | | | | | | |
|---|---|---|---|---|---|---|---|---|---|---|---|---|---|---|---|---|---|
| UNC | 10.64 | 0 | 0 | 0 | 0 | 0 | 100 | 0 | 100 | 19.15 | 0 | 0 | 12.77 | 91.49 | 34.04 | 0 | **31.36** |
| TAP | 2.56 | 2.56 | 0 | 0 | 0 | 0 | 0 | 0 | 0 | 0 | 0 | 0 | 0 | 0 | 15.38 | 0 | **6.39** |
| PCT - bilateral | 0 | 0 | 0 | 0 | 0 | 0 | 0 | 0 | 0 | 0 | 0 | 0 | 0 | 0 | 0 | 0 | **0.00** |
| BCC - bilateral | 1.16 | 1.68 | 0 | 0 | 0 | 0 | 10.31 | 0.52 | 0 | 0 | 0 | 0 | 0 | 0.52 | 0.58 | 0 | **1.80** |
| FX - bilateral | 0 | 0 | 0 | 0 | 0 | 0 | 0 | 0 | 0 | 0 | 0 | 0 | 0 | 0 | 0 | 0 | **0.00** |
| MCP - bilateral | 0 | 0 | 0 | 0 | 0 | 0 | 0 | 0 | 0 | 0 | 0 | 0 | 0 | 0 | 0 | 0 | **0.00** |
| GCC - bilateral | 1.33 | 0 | 0 | 0 | 0 | 0 | 11.49 | 0 | 0 | 0 | 0 | 0 | 1.41 | 0.8 | 0.71 | 0 | **2.08** |
| SCC - bilateral | 0.32 | 5.96 | 0 | 0 | 0 | 0 | 0 | 0 | 0 | 0 | 0 | 0 | 0 | 0 | 0.84 | 0 | **1.09** |

### B. Left hemisphere damage (LHD) group

| Area | 2001 | 2002 | 2003 | 2004 | 2005 | 2006 | 2007 | 2008 | 2009 | 2010 | 2011 | 2012 | 2013 | 2014 | 2015 | 2016 | 2017 | 2018 | 2019 |
|---|---|---|---|---|---|---|---|---|---|---|---|---|---|---|---|---|---|---|---|
| Precentral g | 25.84 | 7.88 | 0 | 7.2 | 61.32 | 0 | 0 | 35.42 | 6.01 | 0 | 0 | 0 | 0 | 0 | 0.2 | 0.34 | 0 | 0 | 0 |
| Superior frontal g | 14.39 | 16.95 | 0 | 0 | 37.45 | 0 | 0 | 0.03 | 0.56 | 0 | 0 | 0 | 0 | 0 | 27.9 | 0 | 0 | 0 | 0 |
| Sup. frontal g, orbital | 0 | 0 | 0 | 0 | 0 | 0 | 0 | 0 | 0 | 0 | 0 | 0 | 0 | 0 | 12.77 | 0 | 0 | 0 | 0 |
| Mid. frontal g | 4.19 | 14.7 | 0 | 0 | 20.15 | 0 | 0 | 0.8 | 4.46 | 0 | 0 | 0 | 0 | 0 | 1.11 | 0.41 | 0 | 0 | 0 |
| Mid. frontal g, orbital | 0 | 0 | 0 | 0 | 0 | 0 | 0 | 0 | 0 | 0 | 0 | 0 | 0 | 0 | 0 | 0.11 | 0 | 0 | 0 |
| IFG$_{OP}$ | 0 | 12.24 | 0 | 0 | 10.98 | 0 | 0 | 2.99 | 83.72 | 0 | 0 | 0 | 0 | 0 | 0 | 10.4 | 0 | 0 | 0 |
| IFG$_{TRI}$ | 0 | 12.3 | 0 | 0 | 1.19 | 0 | 0 | 2.85 | 50.49 | 0 | 0 | 0 | 0 | 0 | 0 | 17.04 | 0 | 0 | 0 |
| IFG$_{ORB}$ | 0 | 0 | 0 | 0 | 0 | 0 | 0 | 0 | 23.31 | 0 | 0 | 0 | 0 | 0 | 1.42 | 12.43 | 0 | 0 | 0 |
| Rolandic op. | 0 | 1.31 | 3.54 | 0 | 4.44 | 0 | 29.6 | 46.67 | 98.59 | 2.42 | 4.04 | 0 | 25.35 | 0 | 0 | 4.95 | 0 | 0 | 0 |
| Supp. motor area | 32.56 | 8.85 | 0 | 0.19 | 66 | 0 | 0 | 0 | 0 | 0 | 0 | 0 | 0 | 0 | 8.94 | 0 | 0 | 0 | 0 |
| Olfactory | 0 | 0 | 0 | 0 | 0 | 0 | 0 | 0 | 0.36 | 0 | 0 | 0 | 0 | 0 | 19.64 | 0.71 | 0 | 1.79 | 0 |
| Sup. frontal g, medial | 0.03 | 1.54 | 0 | 0 | 14.87 | 0 | 0 | 0 | 0.07 | 0 | 0 | 0 | 0 | 0 | 40.88 | 0 | 0 | 0 | 0 |
| Sup. frontal g, med. orb. | 0 | 0 | 0 | 0 | 0 | 0 | 0 | 0 | 0 | 0 | 0 | 0 | 0 | 0 | 0.83 | 0 | 0 | 0 | 0 |
| Rectus g | 0 | 0 | 0 | 0 | 0 | 0 | 0 | 0 | 0 | 0 | 0 | 0 | 0 | 0 | 24.77 | 0 | 0 | 0 | 0 |
| Insula | 0.05 | 9.53 | 0.11 | 0 | 2.15 | 0 | 6.84 | 4.14 | 92.09 | 0.22 | 3.55 | 0 | 2.1 | 0 | 2.05 | 48.17 | 0 | 14.75 | 0 |
| Anterior cingulum | 0 | 19 | 0 | 0 | 8.86 | 0 | 0 | 0 | 0 | 0 | 0 | 0 | 0 | 0 | 31.93 | 0 | 0 | 0 | 0 |
| Middle cingulum | 23.18 | 16.13 | 2.01 | 0.15 | 35.91 | 0 | 0 | 0 | 0.26 | 0 | 0 | 0 | 0 | 0 | 12.16 | 0 | 0 | 0 | 0 |
| Posterior cingulum | 0 | 0 | 0.86 | 0 | 1.73 | 0 | 0 | 0 | 0 | 0 | 0 | 0 | 0 | 0 | 0 | 0 | 0 | 0 | 0 |
| Hippocampus | 0 | 0 | 0 | 0 | 0 | 0 | 0 | 0 | 0.11 | 21.03 | 0 | 0 | 0 | 16.09 | 0 | 0.97 | 20.28 | 0.21 | 1.29 |
| Parahippocampal g | 0 | 0 | 0 | 0 | 0 | 0 | 0 | 0 | 0 | 2.56 | 0 | 0 | 0 | 33.64 | 0 | 1.23 | 22.29 | 0 | 0.41 |
| Amygdala | 0 | 0 | 0 | 0 | 0 | 0 | 0 | 0 | 4.09 | 0 | 0 | 0 | 0 | 0 | 0 | 20.45 | 0 | 15.45 | 0 |
| Calcarine | 0 | 0 | 6.82 | 0 | 0.09 | 0 | 0 | 0 | 0 | 0.58 | 1.68 | 0 | 0 | 38.49 | 0 | 0 | 33.88 | 0 | 0 |
| Cuneus | 0 | 0 | 9.04 | 0 | 6.75 | 0 | 0 | 0 | 0 | 0.79 | 0 | 0 | 0 | 23.79 | 0 | 0 | 12.32 | 0 | 0 |
| Lingual g | 0 | 0 | 0 | 0 | 0 | 0 | 0 | 0 | 0 | 0.43 | 0 | 0 | 0 | 72.17 | 0 | 0 | 53.7 | 0 | 0 |
| Superior occipital g | 0.22 | 0 | 32.21 | 0 | 23.06 | 0 | 0 | 0 | 0 | 0.66 | 0.81 | 0 | 0 | 24.16 | 0 | 0 | 10.91 | 0 | 0 |
| Middle occipital g | 0.67 | 0 | 38.26 | 0 | 15.96 | 3.09 | 0 | 0 | 0 | 4.25 | 0.76 | 0.43 | 1.44 | 8.56 | 0 | 0 | 7.77 | 0 | 0 |
| Inferior occipital g | 0 | 0 | 0 | 0 | 0 | 0 | 0 | 0 | 0 | 2.02 | 0 | 0 | 0 | 50.16 | 0 | 0 | 45.27 | 0 | 0 |
| Fusiform g | 0 | 0 | 0 | 0 | 0 | 0 | 0 | 0 | 0 | 0.22 | 0 | 0 | 0 | 59.13 | 0 | 0 | 38.35 | 0 | 3.81 |
| Postcentral g | 28.37 | 5.16 | 0 | 2.67 | 62.64 | 0 | 3.83 | 28.06 | 10.07 | 0.03 | 2 | 12.82 | 10.41 | 0 | 0 | 0 | 0 | 0 | 0 |
| Superior parietal g | 26.44 | 0.15 | 43.44 | 0 | 26.92 | 0 | 0 | 0 | 0 | 0 | 0 | 15.11 | 28.67 | 0 | 0 | 0 | 0 | 0 | 0 |
| Inferior parietal g | 12.46 | 1.59 | 53.13 | 0 | 47.4 | 8.87 | 0.53 | 0 | 0 | 0 | 4.33 | 64.73 | 80.34 | 0 | 0 | 0 | 0 | 0 | 0 |
| Supramarginal g | 0 | 0 | 20.94 | 0 | 50.56 | 12.1 | 36.94 | 6.21 | 11.23 | 22.13 | 16 | 25.16 | 92.91 | 0 | 0 | 0 | 0 | 0 | 0 |

| | | | | | | | | | | | | | | | | | | | |
|---|---|---|---|---|---|---|---|---|---|---|---|---|---|---|---|---|---|---|---|
| Angular g | 0.51 | 0 | 74.08 | 0 | 19.52 | 34.27 | 0 | 0 | 0 | 21.48 | 1.62 | 5.63 | 65.64 | 0 | 0 | 0 | 0 | 0 | 0 |
| Precuneus | 21.83 | 3.03 | 19.42 | 0.54 | 19.84 | 0 | 0 | 0 | 0 | 0.2 | 0.06 | 0.03 | 0 | 0 | 0 | 0 | 2.13 | 0 | 0 |
| Paracentral lobule | 66.64 | 10.67 | 0 | 25.95 | 54.19 | 0 | 0 | 0 | 0 | 0 | 0 | 0 | 0 | 0 | 0 | 0 | 0 | 0 | 0 |
| Caudate | 0.73 | 0 | 0 | 0 | 13.83 | 0 | 0 | 0 | 11.54 | 0 | 0 | 0 | 0 | 0 | 4.26 | 8.94 | 0 | 29.11 | 0 |
| Putamen | 0 | 3.27 | 0 | 0 | 0 | 0 | 0 | 0 | 59.96 | 0.2 | 0.3 | 0 | 0 | 0 | 15.06 | 68.58 | 0 | 95.94 | 0 |
| Pallidum | 0 | 0 | 0 | 0 | 0 | 0 | 0 | 0 | 4.44 | 0 | 0 | 0 | 0 | 0 | 0 | 30.38 | 0 | 67.58 | 5.12 |
| Thalamus | 0 | 0 | 0 | 0 | 0 | 0 | 0 | 0 | 0 | 0.18 | 0 | 0 | 0 | 0 | 0 | 1.64 | 0 | 7.91 | 11.18 |
| Heschl g | 0 | 0 | 0 | 0 | 0 | 0 | 24.44 | 24 | 96.89 | 0.89 | 7.56 | 0 | 50.22 | 0 | 0 | 1.78 | 0 | 8 | 0 |
| Superior temporal g | 0 | 0 | 8.62 | 0 | 0 | 0 | 12.37 | 5.36 | 38.5 | 13.55 | 7.23 | 0 | 43.9 | 0 | 0 | 0.09 | 0 | 3.7 | 0 |
| Superior temporal pole | 0 | 0 | 0 | 0 | 0 | 0 | 0 | 0 | 19.46 | 0 | 0 | 0 | 0 | 0 | 0 | 1.09 | 0 | 3.74 | 0 |
| Middle temporal g | 0 | 0 | 5.32 | 0 | 1.86 | 0 | 0 | 0 | 0.22 | 30.35 | 5.26 | 0 | 1.78 | 0 | 0 | 0 | 0 | 0.3 | 0 |
| Middle temporal pole | 0 | 0 | 0 | 0 | 0 | 0 | 0 | 0 | 0 | 0 | 0 | 0 | 0 | 0 | 0 | 0 | 0 | 0 | 0 |
| Inferior temporal g | 0 | 0 | 0 | 0 | 0 | 0 | 0 | 0 | 0 | 0.38 | 0 | 0 | 0 | 9.03 | 0 | 0 | 0.12 | 0 | 0.06 |
| CST | 0 | 0 | 0 | 0 | 0 | 0 | 0 | 0 | 0 | 0 | 0 | 0 | 0 | 0 | 0 | 0 | 0 | 0 | 0 |
| ML | 0 | 0 | 0 | 0 | 0 | 0 | 0 | 0 | 0 | 0 | 0 | 0 | 0 | 0 | 0 | 0 | 0 | 0 | 0 |
| ICP | 0 | 0 | 0 | 0 | 0 | 0 | 0 | 0 | 0 | 0 | 0 | 0 | 0 | 0 | 0 | 0 | 0 | 0 | 0 |
| SCP | 0 | 0 | 0 | 0 | 0 | 0 | 0 | 0 | 0 | 0 | 0 | 0 | 0 | 0 | 0 | 0 | 0 | 0 | 0 |
| CP | 0 | 0 | 0 | 0 | 0 | 0 | 0 | 0 | 0 | 4.18 | 0 | 0 | 0 | 0 | 0 | 0 | 0.38 | 0.38 | 9.13 |
| $IC_{AL}$ | 0 | 0 | 0 | 0 | 0 | 0 | 0 | 0 | 44.13 | 0 | 0 | 0 | 0 | 0 | 8.93 | 50 | 0 | 83.16 | 0 |
| $IC_{PL}$ | 2.31 | 0 | 0 | 0 | 0 | 0 | 2.31 | 0 | 35.64 | 0 | 11.74 | 0 | 0 | 0 | 0 | 50.1 | 0 | 68.13 | 10.06 |
| $IC_{RL}$ | 0.32 | 0 | 3.86 | 0 | 0 | 0 | 3.22 | 0 | 14.15 | 52.73 | 28.94 | 0 | 2.25 | 0 | 0 | 15.76 | 0 | 37.3 | 0.32 |
| $CR_{A}$ | 0 | 47.28 | 0 | 0 | 17.92 | 0 | 0 | 0 | 29.02 | 0 | 0 | 0 | 0 | 0 | 42.08 | 44.51 | 0 | 3.93 | 0 |
| $CR_{S}$ | 41.77 | 50.97 | 0 | 0 | 97.51 | 2.06 | 2.92 | 7.14 | 38.96 | 0 | 6.17 | 0 | 0 | 0 | 13.53 | 39.61 | 0 | 24.35 | 0 |
| $CR_{P}$ | 20.4 | 17.04 | 51.79 | 0 | 85.2 | 32.29 | 12.33 | 0 | 7.62 | 53.36 | 29.82 | 2.02 | 6.28 | 0 | 0 | 1.79 | 0 | 6.28 | 0 |
| $TR_{P}$ | 0 | 0 | 37.66 | 0 | 7.11 | 0 | 0 | 0 | 0 | 70.92 | 35.98 | 0 | 1.05 | 0.42 | 0 | 0 | 13.18 | 0 | 0 |
| SS | 0 | 0 | 0 | 0 | 0 | 0 | 0 | 0 | 0 | 63.89 | 0 | 0 | 0 | 6.94 | 0 | 0 | 15.28 | 2.08 | 0 |
| External capsule | 1.33 | 10.67 | 0 | 0 | 0 | 0 | 1.33 | 0.22 | 94.22 | 0 | 6 | 0 | 0 | 0 | 4.89 | 94.89 | 0 | 90 | 0 |
| CGC | 1.78 | 51.34 | 0 | 0 | 30.56 | 0 | 0 | 0 | 0 | 0 | 0 | 0 | 0 | 0 | 16.32 | 0 | 0 | 0 | 0 |
| CGH | 0 | 0 | 0 | 0 | 0 | 0 | 0 | 0 | 0 | 2.29 | 0 | 0 | 0 | 44.27 | 0 | 0 | 49.62 | 0 | 0 |
| FX/ST | 0 | 0 | 0 | 0 | 0 | 0 | 0 | 0 | 0 | 59.86 | 0 | 0 | 0 | 2.04 | 0 | 0.68 | 10.88 | 6.8 | 0 |
| SLF | 36.44 | 15.09 | 20.49 | 0 | 79.39 | 13.87 | 23.44 | 23.07 | 25.15 | 37.3 | 45.64 | 9.82 | 20.98 | 0 | 0 | 12.02 | 0 | 0.74 | 0 |
| $FO_{S}$ | 0 | 0 | 0 | 0 | 100 | 0 | 0 | 0 | 87.27 | 0 | 0 | 0 | 0 | 0 | 0 | 67.27 | 0 | 69.09 | 0 |
| $FO_{I}$ | 0 | 0 | 0 | 0 | 0 | 0 | 0 | 0 | 52.07 | 2.07 | 0 | 0 | 0 | 0 | 30.17 | 54.96 | 0 | 92.15 | 0 |
| UNC | 0 | 0 | 0 | 0 | 0 | 0 | 0 | 0 | 55.1 | 0 | 0 | 0 | 0 | 0 | 0 | 36.73 | 0 | 87.76 | 0 |
| TAP | 0 | 0 | 83.1 | 0 | 0 | 0 | 0 | 0 | 0 | 91.55 | 1.41 | 0 | 0 | 0 | 0 | 0 | 2.82 | 0 | 0 |
| PCT- bilateral | 0 | 0 | 0 | 0 | 0 | 0 | 0 | 0 | 0 | 0 | 0 | 0 | 0 | 0 | 0 | 0 | 0 | 0 | 0 |
| BCC- bilateral | 1.51 | 11.81 | 0 | 0 | 25.19 | 0 | 0 | 0 | 0 | 0 | 0 | 0 | 0 | 0 | 6.43 | 0.81 | 0 | 0 | 0 |
| FX- bilateral | 0 | 0 | 0 | 0 | 0 | 0 | 0 | 0 | 0 | 0 | 0 | 0 | 0 | 0 | 0 | 0 | 0 | 0 | 0 |
| MCP- bilateral | 0 | 0 | 0 | 0 | 0 | 0 | 0 | 0 | 0 | 0 | 0 | 0 | 0 | 0 | 0 | 0 | 0 | 0 | 0 |
| GCC- bilateral | 0 | 18.3 | 0 | 0 | 2.48 | 0 | 0 | 0 | 0 | 0 | 0 | 0 | 0 | 0 | 16.53 | 0.62 | 0 | 0.35 | 0 |
| SCC- bilateral | 0 | 0.32 | 9.66 | 0 | 7.71 | 0.45 | 0 | 0 | 0 | 5.06 | 0.71 | 0 | 0 | 0.19 | 0 | 0 | 3.63 | 0 | 0 |

**B. Left hemisphere damage (LHD) group (cont.)**

| Areas | 2020 | 2021 | 2022 | 2023 | 2024 | 2025 | 2026 | 2027 | 2028 | 2029 | 2030 | 2031 | 2032 | 2033 | 2033 | 2035 | 2036 |
|---|---|---|---|---|---|---|---|---|---|---|---|---|---|---|---|---|---|
| Precentral g | 0 | 55.16 | 0 | 0 | 7.23 | 0 | 0 | 0 | 0 | 0.09 | 0 | 2.5 | 2.3 | 0 | 0 | 0 | 4.88 |
| Superior frontal g | 0.69 | 3.08 | 0 | 0 | 0 | 0 | 0 | 0 | 0 | 0 | 0 | 0 | 0 | 0 | 0 | 0 | 0 |
| Sup. frontal g, orbital | 0 | 0 | 0 | 0 | 0.52 | 0 | 0 | 0 | 0 | 0 | 0 | 0.83 | 2.6 | 0 | 0 | 0 | 0 |
| Mid. frontal g | 2.12 | 36.79 | 0 | 0 | 0.41 | 0 | 0 | 0 | 0 | 0.02 | 0 | 2.76 | 0.12 | 0 | 0 | 0 | 0 |
| Mid. frontal g, orbital | 0 | 0 | 0 | 0 | 0 | 0 | 0 | 0 | 0 | 0 | 0 | 0 | 0 | 0 | 0 | 0 | 0 |
| IFG$_{OP}$ | 1.73 | 95.57 | 0 | 0 | 51.06 | 0 | 0 | 0 | 0 | 0 | 0 | 5.49 | 4.05 | 0 | 0 | 0 | 0 |
| IFG$_{TRI}$ | 2.25 | 79.72 | 0 | 0 | 31.47 | 0 | 0 | 0 | 0 | 0 | 0 | 2.37 | 2.61 | 0 | 0 | 0 | 0 |
| IFG$_{ORB}$ | 0 | 0.47 | 0 | 0 | 14.14 | 0 | 0 | 0 | 0 | 0 | 0.3 | 1.72 | 2.31 | 0 | 0 | 0 | 0 |
| Rolandic op. | 0 | 42.63 | 0 | 2.22 | 41.82 | 0 | 1.31 | 14.04 | 0 | 0 | 0 | 0 | 26.36 | 0 | 0 | 0 | 0 |
| Supp. motor area | 0 | 0 | 0 | 0 | 0 | 0 | 0 | 0 | 0 | 0 | 0 | 0 | 0 | 0 | 0 | 0 | 1.07 |
| Olfactory | 0 | 0 | 0 | 0 | 2.86 | 0 | 0 | 0 | 0 | 0 | 0.71 | 0 | 3.57 | 0 | 0 | 0 | 0 |
| Sup. frontal g, medial | 0.07 | 0 | 0 | 0 | 0 | 0 | 0 | 0 | 0 | 0 | 0 | 0 | 0 | 0 | 0 | 0 | 0 |
| Sup. frontal g, med. orb. | 0 | 0 | 0 | 0 | 0 | 0 | 0 | 0 | 0 | 0 | 0 | 0 | 0 | 0 | 0 | 0 | 0 |
| Rectus g | 0 | 0 | 0 | 0 | 0 | 0 | 0 | 0 | 0 | 0 | 0 | 1.41 | 1.53 | 0 | 0 | 0 | 0 |
| Insula | 0.54 | 27.13 | 0 | 8.56 | 79.17 | 0.97 | 6.4 | 25.03 | 0.05 | 0 | 5.71 | 14.64 | 84.34 | 0 | 0.27 | 0 | 0 |
| Anterior cingulum | 0 | 0 | 0 | 0 | 0 | 0 | 0 | 0 | 0 | 0 | 0 | 0 | 0 | 0 | 0 | 0 | 0 |
| Middle cingulum | 4.48 | 0.46 | 0 | 0 | 0 | 0 | 0 | 0 | 0 | 0 | 0 | 0 | 0 | 0 | 0 | 0 | 9.84 |
| Posterior cingulum | 73.43 | 0 | 0 | 0 | 0 | 0 | 0 | 0 | 0 | 0 | 0 | 0 | 0 | 0 | 0 | 0 | 0 |
| Hippocampus | 91.09 | 0 | 0 | 0.64 | 6.87 | 0 | 0 | 0.32 | 0 | 0 | 2.04 | 0 | 22.75 | 0 | 0 | 8.58 | 0 |
| Parahippocampal g | 66.77 | 0 | 0 | 0 | 0 | 0 | 0 | 0 | 0 | 0 | 0 | 0 | 0.1 | 0 | 0 | 19.63 | 0 |
| Amygdala | 23.18 | 0 | 0 | 0 | 26.82 | 0 | 0 | 0 | 0 | 0 | 9.09 | 0 | 9.09 | 0 | 0 | 0 | 0 |
| Calcarine | 88.62 | 0 | 0 | 0 | 0 | 0 | 0 | 0 | 0 | 0 | 0 | 0 | 0 | 0 | 0 | 37.95 | 0 |
| Cuneus | 60.42 | 0 | 0 | 0 | 0 | 0 | 0 | 0 | 0 | 0 | 0 | 0 | 0 | 0 | 0 | 0.13 | 0 |
| Lingual g | 96.32 | 0 | 0 | 0 | 0 | 0 | 0 | 0 | 0 | 0 | 0 | 0 | 0 | 0 | 0 | 74.37 | 0 |
| Superior occipital g | 12.81 | 0 | 0 | 0 | 0 | 0 | 0 | 0 | 0 | 0 | 0 | 0 | 0 | 0 | 0 | 1.68 | 0.59 |
| Middle occipital g | 5.99 | 0 | 0 | 0 | 0 | 0 | 0 | 0 | 0 | 0 | 0 | 0 | 0 | 0 | 0 | 1.8 | 0.89 |
| Inferior occipital g | 72.79 | 0 | 0 | 0 | 0 | 0 | 0 | 0 | 0 | 0 | 0 | 0 | 0 | 0 | 0 | 19.98 | 0 |
| Fusiform g | 87.1 | 0 | 0 | 0 | 1.34 | 0 | 0 | 0 | 0 | 0 | 0 | 0 | 2.94 | 0 | 0 | 14.2 | 0 |
| Postcentral g | 0 | 17.21 | 0 | 0 | 2.8 | 0 | 0 | 0.08 | 0 | 0 | 0 | 0.08 | 1.1 | 0 | 0 | 0 | 20.4 |
| Superior parietal g | 3 | 0 | 0 | 0 | 0 | 0 | 0 | 0 | 0 | 0 | 0 | 0 | 0 | 0 | 0 | 0 | 42.03 |
| Inferior parietal g | 0 | 0 | 0 | 0 | 0 | 0 | 0 | 0 | 0 | 0 | 0 | 0 | 0.37 | 0 | 0 | 0 | 23.33 |
| Supramarginal g | 0 | 0 | 0 | 0 | 0.08 | 0 | 0 | 12.82 | 0 | 0 | 0 | 0 | 19.59 | 0 | 0 | 0 | 6.61 |
| Angular g | 0 | 0 | 0 | 0 | 0 | 0 | 0 | 4.01 | 0 | 0 | 0 | 0 | 0.68 | 0 | 0 | 0 | 4.01 |
| Precuneus | 41.52 | 0 | 0 | 0 | 0 | 0 | 0 | 0 | 0 | 0 | 0 | 0 | 0 | 0 | 0 | 1.59 | 16.07 |
| Paracentral lobule | 0 | 0 | 0 | 0 | 0 | 0 | 0 | 0 | 0 | 0 | 0 | 0 | 0 | 0 | 0 | 0 | 16.75 |
| Caudate | 0.83 | 1.25 | 5.61 | 2.18 | 44.49 | 39.19 | 0.21 | 0.52 | 6.24 | 0 | 0.31 | 10.5 | 12.27 | 0 | 0 | 0 | 0 |
| Putamen | 3.37 | 2.48 | 12.19 | 29.83 | 66.8 | 38.85 | 22.4 | 4.56 | 13.97 | 0 | 76.71 | 48.86 | 70.27 | 0 | 1.29 | 0 | 0 |
| Pallidum | 0.68 | 0 | 1.02 | 11.95 | 27.99 | 17.41 | 3.07 | 0 | 0.34 | 0 | 28.33 | 6.48 | 27.3 | 0 | 3.41 | 0 | 0 |
| Thalamus | 26.55 | 0 | 2.64 | 0.09 | 0 | 2.09 | 3.55 | 0 | 0.64 | 0 | 0.27 | 8.09 | 17 | 0 | 21.64 | 0 | 0 |
| Heschl g | 0 | 0 | 0 | 32 | 4.89 | 0 | 4.44 | 20.44 | 0 | 0 | 0 | 4.44 | 64.89 | 0 | 0 | 0 | 0 |
| Superior temporal g | 0.26 | 3.22 | 0 | 16.38 | 14.46 | 0 | 0 | 20.17 | 0 | 0 | 0 | 0.17 | 12.2 | 0 | 0 | 0 | 0 |
| Superior temporal pole | 0 | 4.82 | 0 | 1.87 | 28.87 | 0 | 0 | 0.62 | 0 | 0 | 0 | 0 | 2.8 | 0 | 0 | 0 | 0 |

| | | | | | | | | | | | | | | | | | |
|---|---|---|---|---|---|---|---|---|---|---|---|---|---|---|---|---|---|
| Middle temporal g | 3.68 | 0 | 0 | 2.79 | 18.23 | 0 | 0 | 1.72 | 0 | 0 | 0 | 0 | 4.98 | 0 | 0 | 0 | 0 |
| Middle temporal pole | 0 | 0 | 0 | 0 | 2.78 | 0 | 0 | 0 | 0 | 0 | 0 | 0 | 0.66 | 0 | 0 | 0 | 0 |
| Inferior temporal g | 36.22 | 0 | 0 | 0 | 1.78 | 0 | 0 | 0 | 0 | 0 | 0 | 0 | 20.94 | 0 | 0 | 0.44 | 0 |
| CST | 0 | 0 | 0 | 0 | 0 | 0 | 0 | 0 | 0 | 0 | 0 | 0 | 0 | 12.36 | 0 | 0 | 0 |
| ML | 0 | 0 | 0 | 0 | 0 | 0 | 0 | 0 | 0 | 0 | 0 | 0 | 0 | 3.61 | 0 | 0 | 0 |
| ICP | 0 | 0 | 0 | 0 | 0 | 0 | 0 | 0 | 0 | 0 | 0 | 0 | 0 | 0 | 0 | 0 | 0 |
| SCP | 0 | 0 | 0 | 0 | 0 | 0 | 0 | 0 | 0 | 0 | 0 | 0 | 0 | 0 | 0 | 0 | 0 |
| CP | 7.98 | 0 | 0 | 0 | 0 | 0 | 3.42 | 0 | 0 | 0 | 0 | 2.66 | 2.66 | 0 | 12.55 | 0 | 0 |
| ICAL | 5.1 | 11.99 | 0 | 0 | 44.64 | 46.68 | 7.14 | 2.55 | 3.06 | 0 | 27.3 | 40.82 | 40.82 | 0 | 0 | 0 | 0 |
| ICPL | 36.48 | 0.21 | 38.57 | 24.53 | 0 | 34.8 | 51.36 | 3.77 | 35.85 | 0 | 30.19 | 57.44 | 71.91 | 0.42 | 53.04 | 0 | 0 |
| ICRL | 69.13 | 0 | 26.37 | 34.08 | 9.32 | 0.64 | 25.08 | 24.12 | 2.25 | 0 | 8.68 | 42.77 | 92,28 | 0 | 7.07 | 0 | 0 |
| CRA | 29.13 | 24.05 | 0 | 0 | 37.23 | 0 | 0.35 | 0.23 | 0 | 0 | 2.66 | 27.51 | 40.92 | 0 | 0 | 0 | 0 |
| CRS | 15.x91 | 47.19 | 12.88 | 20.35 | 47.4 | 11.26 | 38.1 | 14.5 | 4.98 | 4 | 14.07 | 59.9 | 53.35 | 0 | 0 | 0 | 26.52 |
| CRP | 15.02 | 0 | 8.07 | 15.92 | 12.78 | 0.22 | 9.87 | 36.55 | 2.69 | 0 | 4.93 | 8.7 | 45.29 | 0 | 0 | 0 | 65.02 |
| TRP | 89.12 | 0 | 0 | 0 | 7.53 | 0 | 0 | 4.81 | 0 | 0 | 0 | 0 | 10.04 | 0 | 0 | 6.9 | 0 |
| SS | 100 | 0 | 0 | 14.24 | 28.82 | 0 | 0 | 4.17 | 0 | 0 | 1.04 | 3.82 | 90.28 | 0 | 0 | 15.28 | 0 |
| External capsule | 4.22 | 19.11 | 8.22 | 39.11 | 55.56 | 36.67 | 47.33 | 22.44 | 8.89 | 0 | 88.22 | 82.67 | 100 | 0 | 5.11 | 0 | 0 |
| CGC | 13.06 | 0 | 0 | 0 | 0 | 0 | 0 | 0 | 0 | 0 | 0 | 0 | 0 | 0 | 0 | 0 | 2.37 |
| CGH | 100 | 0 | 0 | 0 | 0 | 0 | 0 | 0 | 0 | 0 | 0 | 0 | 0 | 0 | 0 | 19.85 | 0 |
| FX/ST | 100 | 0 | 0 | 0.68 | 12.24 | 0 | 0 | 0.68 | 0 | 0 | 6.8 | 0 | 54.42 | 0 | 0 | 0 | 0 |
| SLF | 1.6 | 37.42 | 0 | 0.98 | 28.83 | 0 | 10.43 | 39.14 | 0 | 0 | 0 | 26.13 | 71.66 | 0 | 0 | 0 | 50.55 |
| FOS | 10.91 | 58.18 | 5.45 | 0 | 81.82 | 58.18 | 25.45 | 0 | 16.36 | 0 | 14.55 | 92.73 | 96.36 | 0 | 0 | 0 | 0 |
| FOI | 6.61 | 0 | 2.89 | 46.28 | 66.94 | 4.96 | 9.09 | 22.73 | 6.2 | 0 | 54.55 | 76.45 | 90.5 | 0 | 0 | 0 | 0 |
| UNC | 6.12 | 0 | 0 | 42.86 | 100 | 0 | 0 | 6.12 | 0 | 0 | 34.69 | 14.29 | 81.63 | 0 | 0 | 0 | 0 |
| TAP | 95.77 | 0 | 0 | 0 | 0 | 0 | 0 | 11.27 | 0 | 0 | 0 | 0 | 0 | 0 | 0 | 0 | 4.23 |
| PCT - bilateral | 0 | 0 | 0 | 0 | 0 | 0 | 0 | 0 | 0 | 0 | 0 | 0 | 0 | 6.56 | 0 | 0 | 0 |
| BCC - bilateral | 3.01 | 0.87 | 0.52 | 0 | 0.06 | 0.06 | 0 | 0 | 0 | 0 | 0 | 0.17 | 0.64 | 0 | 0 | 0 | 3.42 |
| FX - bilateral | 0 | 0 | 0 | 0 | 0 | 0 | 0 | 0 | 0 | 0 | 0 | 0 | 0 | 0 | 0 | 0 | 0 |
| MCP - bilateral | 0 | 0 | 0 | 0 | 0 | 0 | 0 | 0 | 0 | 0 | 0 | 0 | 0 | 0.21 | 0 | 0 | 0 |
| GCC - bilateral | 0.97 | 0.18 | 0 | 0 | 1.33 | 0 | 0 | 0 | 0 | 0 | 0 | 0.53 | 0.09 | 0 | 0 | 0 | 0 |
| SCC - bilateral | 42.32 | 0 | 0 | 0 | 0 | 0 | 0 | 0 | 0 | 0 | 0 | 0 | 0 | 0 | 0 | 3.11 | 3.18 |

### B. Left hemisphere damage (LHD) group (cont.)

| Areas | 2037 | 2038 | 2039 | **Average** |
|---|---|---|---|---|
| Precentral g | 0 | 0 | 0 | **5.55** |
| Superior frontal g | 0 | 0 | 0 | **2.59** |
| Sup. frontal g, orbital | 0 | 0 | 0 | **0.43** |
| Mid. frontal g | 0 | 0 | 0 | **2.26** |
| Mid. frontal g, orbital | 0 | 0 | 0 | **0.00** |
| IFGOP | 0 | 0 | 0 | **7.13** |
| IFGTRI | 0 | 0 | 0 | **5.19** |
| IFGORB | 0 | 0 | 0 | **1.44** |
| Rolandic op. | 0 | 0 | 0 | **8.96** |
| Supp. motor area | 0 | 0 | 0 | **3.02** |

| | | | | |
|---|---|---|---|---|
| Olfactory | 0 | 0 | 0 | **0.76** |
| Sup. frontal g, medial | 0 | 0 | 0 | **1.47** |
| Sup. frontal g, med. orb. | 0 | 0 | 0 | **0.02** |
| Rectus g | 0 | 0 | 0 | **0.71** |
| Insula | 0.81 | 0 | 0.91 | **11.29** |
| Anterior cingulum | 0 | 0 | 0 | **1.53** |
| Middle cingulum | 0 | 0 | 0 | **2.68** |
| Posterior cingulum | 0 | 0 | 0 | **1.95** |
| Hippocampus | 0 | 0 | 0 | **4.93** |
| Parahippocampal g | 0 | 0 | 0 | **3.76** |
| Amygdala | 0 | 0 | 0 | **2.77** |
| Calcarine | 0 | 0 | 0 | **5.34** |
| Cuneus | 0 | 0 | 0 | **2.90** |
| Lingual g | 0 | 0 | 0 | **7.62** |
| Superior occipital g | 0 | 0 | 0 | **2.75** |
| Middle occipital g | 0 | 0 | 0 | **2.30** |
| Inferior occipital g | 0 | 0 | 0 | **4.88** |
| Fusiform g | 0 | 0 | 0 | **5.31** |
| Postcentral g | 0 | 0 | 0 | **5.33** |
| Superior parietal g | 0 | 0 | 0 | **4.76** |
| Inferior parietal g | 0 | 0 | 0 | **7.62** |
| Supramarginal g | 0 | 0 | 0 | **8.55** |
| Angular g | 0 | 0 | 0 | **5.93** |
| Precuneus | 0 | 0 | 0 | **3.24** |
| Paracentral lobule | 0 | 0 | 0 | **4.47** |
| Caudate | 23.49 | 2.91 | 0 | **5.60** |
| Putamen | 62.83 | 9.91 | 0 | **18.14** |
| Pallidum | 54.27 | 0 | 0 | **7.43** |
| Thalamus | 0.27 | 0 | 0 | **2.66** |
| Heschl g | 0 | 0 | 0 | **8.84** |
| Superior temporal g | 0 | 0 | 0 | **5.13** |
| Superior temporal pole | 0 | 0 | 0 | **1.62** |
| Middle temporal g | 0 | 0 | 0 | **1.96** |
| Middle temporal pole | 0 | 0 | 0 | **0.09** |
| Inferior temporal g | 0 | 0 | 0 | **1.77** |
| CST | 0 | 0 | 0 | **0.32** |
| ML | 0 | 0 | 0 | **0.09** |
| ICP | 0 | 0 | 0 | **0.00** |
| SCP | 0 | 0 | 0 | **0.00** |
| CP | 0 | 0 | 0 | **1.11** |
| $IC_{AL}$ | 78.06 | 7.91 | 0.26 | **12.89** |
| $IC_{PL}$ | 9.22 | 22.64 | 0.42 | **16.70** |
| $IC_{RL}$ | 0 | 0 | 0 | **10.75** |
| $CR_{A}$ | 2.2 | 0 | 0 | **8.95** |
| $CR_{S}$ | 0 | 21.86 | 12.45 | **18.71** |

| | | | | |
|---|---|---|---|---|
| $CR_P$ | 0 | 0.45 | 0 | **14.15** |
| $TR_P$ | 0 | 0 | 0 | **7.30** |
| SS | 0 | 0 | 0 | **8.87** |
| External capsule | 32 | 17.33 | 4.67 | **22.44** |
| CGC | 0 | 0 | 0 | **2.96** |
| CGH | 0 | 0 | 0 | **5.54** |
| FX/ST | 0 | 0 | 0 | **6.54** |
| SLF | 0 | 0 | 6.13 | **16.32** |
| $FO_S$ | 12.73 | 47.27 | 5.45 | **21.77** |
| $FO_I$ | 36.36 | 0 | 0 | **16.79** |
| UNC | 14.29 | 0 | 0 | **12.30** |
| TAP | 0 | 0 | 0 | **7.44** |
| PCT - bilateral | 0 | 0 | 0 | **0.17** |
| BCC - bilateral | 0 | 0 | 0 | **1.40** |
| FX - bilateral | 0 | 0 | 0 | **0.00** |
| MCP - bilateral | 0 | 0 | 0 | **0.01** |
| GCC - bilateral | 0 | 0 | 0 | **1.06** |
| SCC - bilateral | 0 | 0 | 0 | **1.96** |

g = gyrus; $TP_{S/M}$ = superior/middle parts of the temporal pole; $CR_{S/A/P}$ = superior/anterior/posterior part of the corona radiata; $FO_{S/I}$ = superior/inferior portion of the fronto-occipital fasciculus; $IFG_{TRI/ORB/OP}$ = triangular/orbital/opercular parts of the inferior frontal gyrus; SLF = superior longitudinal fasciculus; UNC = uncinate fasciculus; $TR_P$ = posterior thalamic radiation; $IC_{AL/PL/RL}$ = anterior/posterior/retrolenticular limb of the internal capsule; Rolandic op. = Rolandic operculum; CST = corticospinal tract, ML = medial lemniscus, ICP/SCP/MCP = inferior/superior/middle cerebellar peduncle; CP = cerebral peduncle; SS = sagittal stratum; CGC = cingulum (in the cingulate gyrus); CGH = cingulum (in the hippocampal region); FX/ST = fornix and stria terminalis; TAP = tapetum; PCT = pontine crossing tract; FX = fornix; GCC/BCC/SCC =genu/body/splenium of the corpus callosum.

**Table S3:**

**A. Brain regions in which the percent of subjects who had at least 1% of the region damaged by the stroke was significantly larger in the RHD group compared to the LHD group:**

Inferior frontal gyrus, opercular part (67% vs. 26%, $p < .001$)
Middle frontal gyrus, orbital part (26% vs. 0%, $p < .001$)
Heschl gyrus (72% vs. 33%, $p < .001$)
Superior temporal gyrus (72% vs. 33%, $p < .001$)
Supramarginal gyrus (72% vs. 33%, $p < .001$)
Rolandic operculum (78% vs. 41%, $p < .001$)
Superior temporal pole (54% vs. 18%, $p < .001$)
Insula (81% vs. 46%, $p < .001$)
Middle temporal pole (30% vs. 3%, $p < .001$)
Inferior frontal gyrus, orbital part (50% vs. 15%, $p < .001$)
Angular gyrus (57% vs. 23%, $p = .001$)
Postcentral gyrus (70% vs. 36%, $p = .001$)
Precentral gyrus (63% vs. 28%, $p = .001$)
Superior longitudinal fasciculus (87% vs. 56%, $p = .001$)

**B. Brain regions in which the average extent of damage (%) was significantly larger in the RHD group compared to the LHD group:**

Rolandic operculum ($\mathrm{U} = 508.5$, $p < .001$)
Heschl gyrus ($U = 526$, $p < .001$)
Inferior frontal gyrus, opercular part ($U = 540.5$, $p < .001$)
Superior temporal pole (Mann-Whitney $U = 572.5$, $p < .001$)
Superior temporal gyrus (Mann-Whitney $U = 545.5$, $p < .001$)
Insula ($\mathrm{U} = 537$, $p < .001$)
Inferior frontal gyrus, orbital part ($U = 586.5$, $p < .001$)
Supramarginal gyrus ($U = 577.5$, $p < .001$)
Precentral gyrus ($\mathrm{U} = 586.5$, $p < .001$)
Middle temporal gyrus ($U = 618$, $p < .001$)
Postcentral gyrus ($\mathrm{U} = 605$, $p < .001$)
Inferior frontal gyrus, triangular part ($U = 631$, $p < .001$)
Middle frontal gyrus, orbital part ($U = 742$, $p < .001$)
Middle temporal pole ($U = 726.5$, $p < .001$)
Middle frontal gyrus ($U = 669.5$, $p = .001$)
Angular gyrus ($U = 677$, $p = .001$)
Superior longitudinal fasciculus ($\mathrm{U} = 537.5$, $p < .001$)
External capsule ($\mathrm{U} = 619$, $p < .001$)
Superior corona radiata ($\mathrm{U} = 635$, $p = .001$)

**Table S4. VLBM analysis, RHD group: Voxel clusters where damage significantly affected identity memory.**

| Structure | Test phase | Z-value | X | Y | Z | Voxels | % Area |
|---|---|---|---|---|---|---|---|
| Superior temporal g | Immediate | 4.25 | 46 | -42 | 16 | 423 | 13.47 |
| | Delayed | 3.43 | 66 | -44 | 18 | 42 | 1.34 |
| SLF | Immediate | 4.43 | 42 | -46 | 6 | 204 | 24.73 |
| | Delayed | - | - | - | - | - | - |
| Middle temporal g | Immediate | 3.64 | 48 | -44 | 16 | 232 | 5.26 |
| | Delayed | 3.34 | 68 | -22 | -18 | 52 | 1.18 |
| | | 3.03 | 54 | -46 | -6 | 31 | 0.7 |
| | | 3.27 | 42 | -64 | -2 | 37 | 0.84 |
| Supramarginal g | Immediate | 3.54 | 44 | -36 | 22 | 94 | 4.76 |
| | Delayed | - | - | - | - | - | - |
| Angular g | Immediate | 4.22 | 44 | -46 | 22 | 79 | 4.51 |
| | Delayed | 3.17 | 42 | -50 | 28 | 35 | 2 |
| Hippocampus | Immediate | 3.75 | 40 | -26 | -14 | 78 | 8.25 |
| | Delayed | 3.63 | 32 | -12 | -12 | 36 | 3.81 |
| $TR_P$ | Immediate | -4.76 | 40 | -38 | 2 | 70 | 14.37 |
| | Delayed | - | - | - | - | - | - |
| $CR_P$ | Immediate | 4.32 | 26 | -44 | 30 | 68 | 15.04 |
| | Delayed | 4.12 | 26 | -44 | 30 | 52 | 11.5 |
| SS | Immediate | 4.08 | 42 | -24 | -14 | 54 | 18.88 |
| | Delayed | - | - | - | - | - | - |
| $IC_{RL}$ | Immediate | 3.74 | 38 | -34 | 4 | 51 | 16.14 |
| | Delayed | - | - | - | - | - | - |

Brain regions defined by the AAL and WM atlases with 'significant' clusters of ≥ 25 voxels. Results presented correspond to z scores of 2.6, ($p \leq .005$). Regions are sorted by the number of 'significant' voxels. X, Y, Z coordinates indicate the voxel that is most superior, posterior, and left in its location within the cluster. RHD = right hemisphere damage; g = gyrus; $CR_P$ = posterior part of the corona radiata; SLF = superior longitudinal fasciculus; $TR_P$ = posterior thalamic radiation; $IC_{RL}$ = retrolenticular limb of the internal capsule; SS = sagittal stratum.

**Table S5. VLBM analysis, RHD group: Voxel clusters where damage significantly affected location memory.**

| Structure | Test phase | Z-value | X | Y | Z | Voxels | % Area |
|---|---|---|---|---|---|---|---|
| Middle temporal g | Immediate | 4.54 | 44 | -44 | 18 | 2046 (1233) | 46.41 (27.97) |
| | Delayed | 3.92 | 48 | 4 | -30 | 43 (108) | 0.98 (2.45) |
| Superior temporal g | Immediate | 4.88 | 48 | -36 | 14 | 1725 (1248) [29] | 54.92 (39.73) [0.92] |
| | Delayed | 3.59 | 68 | -14 | 0 | 74 | 2.36 |
| | | 4.85 | 40 | -32 | 18 | 212 (785) [28] | 6.75 (24.99) [0.89] |
| Angular g | Immediate | 5.07 | 44 | -46 | 22 | 474 (273) | 27.05 (15.58) |
| | Delayed | 4.97 | 44 | -46 | 22 | 85 (151) | 4.85 (8.92) |
| SLF | Immediate | 4.56 | 40 | -38 | 18 | 385 (230) | 46.67 (27.88) |
| | | (3.64) | (34) | (-2) | (28) | (67) | (8.12) |
| | Delayed | 4.59 | 38 | -38 | 22 | 64 (114) | 7.76 (13.82) |
| Supramarginal g | Immediate | 4.19 | 48 | -40 | 22 | 219 (138) | 11.09 (6.99) |
| | Delayed | 4.67 | 46 | -36 | 22 | 31 (85) | 1.57 (4.31) |
| CR$_S$ | Immediate | 4.26 | 26 | -8 | 30 | 209 (147) | 22.72 (15.98) |
| | Delayed | (3.38) | (26) | (-2) | (28) | (48) | (5.22) |
| Putamen | Immediate | 3.87 | 34 | -8 | -4 | 178 (120) | 16.73 (11.28) |
| | Delayed | - | - | - | - | - | - |
| Insula | Immediate | 3.58 | 28 | 14 | -18 | 177 (100) | 10 (5.65) |
| | Delayed | - | - | - | - | - | - |
| Inferior temporal | Immediate | 3.71 | 50 | -44 | -6 | 174 (78) | 4.89 (2.19) |
| | Delayed | - | - | - | - | - | - |
| TP$_S$ | Immediate | 3.55 | 46 | 4 | -20 | 159 (74) | 11.88 (5.53) |
| | Delayed | 3.69 | 42 | 14 | -30 | 80 (141) | 5.98 (10.54) |
| TR$_P$ | Immediate | 4.5 | 38 | -42 | 6 | 121 (98) | 24.85 (20.12) |
| | Delayed | - | - | - | - | - | - |
| FO$_I$ | Immediate | 3.87 | 34 | -8 | -4 | 120 (80) | 45.63 (30.42) |
| | Delayed | - | - | - | - | - | - |
| CR$_P$ | Immediate | 4.25 | 30 | -34 | 24 | 119 (77) | 26.33 (17.04) |
| | Delayed | (4.23) | (32) | (-42) | (24) | (40) | (8.85) |
| IC$_{RL}$ | Immediate | 4.17 | 34 | -36 | 16 | 118 (87) | 37.34 (27.53) |
| | Delayed | - | - | - | - | - | - |
| SS | Immediate | 4.2 | 38 | -18 | -6 | 118 (83) | 41.26 (29.02) |
| | Delayed | - | - | - | - | - | - |
| External capsule | Immediate | 3.62 | 32 | -20 | 6 | 118 (79) | 25.32 (16.95) |
| | Delayed | - | - | - | - | - | - |
| TP$_M$ | Immediate | 3.54 | 44 | 8 | -30 | 116 (88) | 9.77 (7.41) |
| | Delayed | 3.73 | 44 | 14 | -30 | 53 (68) | 4.47 (5.73) |
| Middle occipital g | Immediate | 3.4 | 46 | -64 | 28 | 100 (51) | 4.77 (2.43) |
| | Delayed | - | - | - | - | - | - |
| CR$_A$ | Immediate | 3.72 | 22 | 20 | 16 | 66 | 7.71 |
| | Delayed | - | - | - | - | - | - |
| Heschl g | Immediate | 4.52 | 34 | -28 | 14 | 60 (34) | 24.1 (13.65) |
| | Delayed | (4.69) | (34) | (-28) | (14) | (33) | (13.25) |
| Hippocampus | Immediate | 3.75 | 42 | -16 | -14 | 57 (34) | 6.03 (3.59) |
| | Delayed | - | - | - | - | - | - |
| IFG$_{OP}$ | Immediate | 3 | 42 | 4 | 26 | 46 | 3.29 |
| | Delayed | - | - | - | - | - | - |
| Amygdala | Immediate | 3.91 | 30 | 0 | -14 | 39 (26) | 15.73 (10.48) |
| | Delayed | - | - | - | - | - | - |
| Rolandic op. | Immediate | 3.12 | 40 | -8 | 22 | 39 | 2.93 |
| | Delayed | - | - | - | - | - | - |
| Pallidum | Immediate | 4.42 | 30 | -8 | -4 | 32 | 11.43 |
| | Delayed | - | - | - | - | - | - |
| IC$_{PL}$ | Immediate | 3.81 | 22 | -22 | 16 | 29 | 5.79 |
| | Delayed | - | - | - | - | - | - |

| | | | | | | | |
|---|---|---|---|---|---|---|---|
| Precentral g | Immediate | 3.19 | 34 | 4 | 30 | 27 | 0.8 |
| | Delayed | - | - | - | - | - | - |
| UNC | Immediate | 3.72 | 32 | 0 | -14 | 25 | 53.19 |
| | Delayed | - | - | - | - | - | - |

Brain regions defined by the AAL and WM atlases with ‘significant’ clusters of ≥ 25 voxels. Presented results passed FDR correction for multiple comparisons (corresponding in this analysis to z scores of 2.36 and 3.06, for Immediate and delayed location memory, respectively). Results presented in brackets correspond to z scores of 2.6, ($p \leq .005$). Results presented in square brackets passed permutation correction (corresponding to z scores of 4.29 and 4.32, for Immediate and delayed location memory, respectively). Regions are sorted by the number of ‘significant’ voxels. X, Y, Z coordinates indicate the voxel that is most superior, posterior, and left in its location within the cluster. RHD = right hemisphere damage; g = gyrus; Rolandic op. = Rolandic operculum; SS = sagittal stratum; $CR_{S/P/A}$ = superior/posterior/anterior part of the corona radiata; $FO_{I}$ = inferior portion of the fronto-occipital fasciculus; SLF = superior longitudinal fasciculus; $TR_{P}$ = posterior thalamic radiation; $IC_{RL/PL}$ = retrolenticular/posterior limb of the internal capsule; UNC = uncinate fasciculus; $TP_{S/M}$ = superior/middle part of the temporal pole.

**Table S6. VLBM analysis, RHD group: Voxel clusters where damage significantly affected action memory.**

| Structure | Test phase | Z-value | X | Y | Z | Voxels | % Area |
|---|---|---|---|---|---|---|---|
| Middle temporal g | Immediate | 3.59 | 42 | -52 | 18 | 191 | 4.33 |
| | | 3.12 | 68 | -22 | -18 | 115 | 2.61 |
| | Delayed | 3.2 | 66 | -24 | -12 | 90 | 2.04 |
| Angular g | Immediate | 3.9 | 42 | -50 | 28 | 177 | 10.1 |
| | Delayed | - | - | - | - | - | - |
| SLF | Immediate | 3.86 | 36 | -48 | 18 | 98 | 11.88 |
| | Delayed | - | - | - | - | - | - |
| superior temporal g | Immediate | 3.52 | 44 | -42 | 18 | 87 | 2.77 |
| | Delayed | 3.13 | 48 | -32 | 18 | 43 | 1.37 |
| $CR_S$ | Immediate | 3.41 | 28 | 10 | 26 | 42 | 4.57 |
| | Delayed | - | - | - | - | - | - |
| $TR_P$ | Immediate | 4.08 | 34 | -46 | 18 | 38 | 7.8 |
| | Delayed | - | - | - | - | - | - |
| $CR_P$ | Immediate | 3.21 | 26 | -48 | 26 | 33 | 7.3 |
| | Delayed | - | - | - | - | - | - |
| Middle occipital g | Immediate | 3.08 | 46 | -64 | 28 | 30 | 1.43 |
| | Delayed | - | - | - | - | - | - |
| $IFG_{OP}$ | Immediate | 3.22 | 46 | 6 | 24 | 23 | 1.64 |
| | Delayed | 3.17 | 42 | 4 | 26 | 20 | 1.43 |

Brain regions defined by the AAL and WM atlases with 'significant' clusters of $\geq$ 25 voxels. Presented results correspond in this analysis to z scores of 2.6, ($p \leq .005$). Regions are sorted by the number of 'significant' voxels. X, Y, Z coordinates indicate the voxel that is most superior, posterior, and left in its location within the cluster. RHD = right hemisphere damage; g = gyrus; $CR_{S/P}$ = superior/posterior part of the corona radiata; $IFG_{OP}$ = opercular part of the inferior frontal gyrus; $TR_P$ = posterior part of the thalamic radiation; SLF = superior longitudinal fasciculus.

**Table S7. VLBM analysis, LHD group: Voxel clusters where damage significantly affected identity memory.**

| Structure | Test phase | Z-value | X | Y | Z | Voxels | % Area |
|---|---|---|---|---|---|---|---|
| $IC_{AL}$ | Immediate | (3.88) | (-22) | (18) | (12) | (25) | (6.38) |
| | Delayed | - | - | - | - | - | - |
| Lingual | Immediate | - | - | - | - | - | - |
| | Delayed | 3.21 | -26 | -86 | -18 | 729 | 34.8 |
| Calcarine | Immediate | - | - | - | - | - | - |
| | Delayed | 3.21 | -8 | -86 | -6 | 339 | 15.01 |
| Fusiform | Immediate | - | - | - | - | - | - |
| | Delayed | 3.21 | -22 | -78 | -18 | 208 | 9 |
| Parahippocampal g | Immediate | - | - | - | - | - | - |
| | Delayed | 3.21 | -24 | -28 | -14 | 109 | 11.15 |
| Inferior occipital g | Immediate | - | - | - | - | - | - |
| | Delayed | 3.21 | -26 | -80 | -12 | 92 | 9.78 |
| Hippocampus | Immediate | - | - | - | - | - | - |
| | Delayed | 3.21 | -28 | -28 | -12 | 27 | 2.9 |

Brain regions defined by the AAL and WM atlases with ‘significant’ clusters of ≥ 25 voxels. Results presented passed FDR correction for multiple comparisons (corresponding in this analysis to z scores of 2.59 for Delayed identity memory). The same values correspond also to z scores of 2.6, ($p \leq .005$). Results presented in brackets correspond only to z scores of 2.6. Regions are sorted by the number of ‘significant’ voxels. X, Y, Z coordinates indicate the voxel that is most superior, posterior, and left in its location within the cluster. LHD = right hemisphere damage; g = gyrus; $IC_{AL}$ = anterior limb of the internal capsule.

**Table S8. VLBM analysis, LHD group: Voxel clusters where damage significantly affected location memory.**

| Structure | Test phase | Z-value | X | Y | Z | Voxels | % Area |
|---|---|---|---|---|---|---|---|
| Lingual | Immediate | - | - | - | - | - | - |
| | Delayed | 2.72 | -26 | -86 | -18 | 729 | 34.8 |
| Calcarine | Immediate | - | - | - | - | - | - |
| | Delayed | 2.72 | -8 | -86 | -6 | 333 | 14.75 |
| Fusiform | Immediate | - | - | - | - | - | - |
| | Delayed | 2.72 | -22 | -78 | -18 | 205 | 8.87 |
| Parahippocampal g | Immediate | - | - | - | - | - | - |
| | Delayed | 2.72 | -24 | -28 | -14 | 109 | 11.15 |
| Inferior occipital g | Immediate | - | - | - | - | - | - |
| | Delayed | 2.72 | -26 | -80 | -12 | 92 | 9.78 |
| Hippocampus | Immediate | - | - | - | - | - | - |
| | Delayed | 2.72 | -34 | -30 | -10 | 26 | 2.79 |

Brain regions defined by the AAL and WM atlases with 'significant' clusters of ≥25 voxels. Results presented passed FDR correction for multiple comparisons (corresponding in this analysis to z scores of 2.58, for the Delayed location memory). The same values correspond also to z scores of 2.6, ($p \leq .005$). Regions are sorted by the number of 'significant' voxels. X, Y, Z coordinates indicate the voxel that is most superior, posterior, and left in its location within the cluster. LHD = right hemisphere damage; g = gyrus.

**Table S9. VLBM analysis, LHD group: Voxel clusters where damage significantly affected action memory.**

| Structure | Test phase | Z-value | X | Y | Z | Voxels | % Area |
|---|---|---|---|---|---|---|---|
| Lingual | Immediate | 2.74 | -26 | -86 | -18 | 726 | 34.65 |
| | Delayed | 3.02 | -28 | -62 | -4 | 729 | 34.8 |
| Calcarine | Immediate | 2.74 | -8 | -86 | -6 | 333 | 14.75 |
| | Delayed | 2.94 | -8 | -86 | -6 | 333 | 14.75 |
| Fusiform g | Immediate | 2.74 | -22 | -78 | -18 | 203 | 8.79 |
| | Delayed | 3.02 | -30 | -62 | -4 | 205 | 8.87 |
| Parahippocampal g | Immediate | 2.74 | -24 | -28 | -14 | 107 | 10.94 |
| | Delayed | 3.02 | -34 | -44 | -8 | 109 | 11.15 |
| Inferior occipital g | Immediate | 2.74 | -26 | -80 | -12 | 92 | 9.78 |
| | Delayed | 2.94 | -26 | -80 | -12 | 92 | 9.78 |
| Insula* | Immediate | 3.01 | -30 | 28 | -4 | 83 (82) | 4.47 (4.41) |
| | Delayed | 3.36 | -30 | 28 | -4 | 82 (78) | 4.41 (4.2) |
| $CR_A$* | Immediate | 3.32 | -24 | 24 | 12 | 46 (36) | 5.32 (4.16) |
| | Delayed | 3.4 | -24 | 24 | 12 | 64 (57) | 7.4 (6.59) |

Brain regions defined by the AAL and WM atlases with 'significant' clusters of ≥ 25 voxels. Results presented passed FDR correction for multiple comparisons (corresponding in this analysis to z scores of 2.59 and 2.53 for immediate and delayed identity memory, respectively). The same results also correspond to z scores of 2.6, ($p \leq .005$).* Indicates that results without brackets passed FDR correction, while results presented in brackets correspond to z scores of 2.6. Regions are sorted by the number of 'significant' voxels. X, Y, Z coordinates indicate the voxel that is most superior, posterior, and left in its location within the cluster. LHD = right hemisphere damage; g = gyrus; $CR_A$ = anterior part of the corona radiata.

**Table S10. VLBM conjunction analysis in the Immediate and Delayed testing phases for the RHD and LHD groups.**

### A. RHD group

| | | Identity vs. Location vs. Action | | | | | | |
|---|---|---|---|---|---|---|---|---|
| Region | Test phase | I | L | A | I+L | I+A | L+A | I+L+A |
| IFG$_{OP}$ | Immediate | - | - | - | - | - | - | - |
| | Delayed | - | - | 28 | - | - | - | - |
| Insula | Immediate | - | 106 | - | - | - | - | - |
| | Delayed | - | - | - | - | - | - | - |
| Hippocampus | Immediate | 52 | - | - | 30 | - | - | - |
| | Delayed | 35 | - | - | - | - | - | - |
| Middle occipital g | Immediate | - | 37 | - | - | - | - | - |
| | Delayed | - | - | - | - | - | - | - |
| Supramarginal g | Immediate | - | 55 | - | 81 | - | - | - |
| | Delayed | - | 92 | - | - | - | - | - |
| Angular g | Immediate | - | 99 | - | 18 | - | 99 | 63 |
| | Delayed | - | 114 | - | - | - | 40 | - |
| Putamen | Immediate | - | 113 | - | - | - | - | - |
| | Delayed | - | - | - | - | - | - | - |
| Heschl g | Immediate | - | 36 | - | - | - | - | - |
| | Delayed | - | 34 | - | - | - | - | - |
| Superior temporal g | Immediate | - | 807 | - | 318 | - | 33 | 94 |
| | Delayed | 44 | 699 | - | 41 | - | 71 | - |
| TP$_{S}$ | Immediate | - | 86 | - | - | - | - | - |
| | Delayed | - | 142 | - | - | - | - | - |
| Middle temporal g | Immediate | 30 | 851 | - | 83 | 6 | 165 | 148 |
| | Delayed | 82 | 101 | 69 | - | 56 | - | - |
| TP$_{M}$ | Immediate | - | 86 | - | - | - | - | - |
| | Delayed | - | 68 | - | - | - | - | - |
| Inferior temporal g | Immediate | - | 58 | - | - | - | - | - |
| | Delayed | - | - | - | - | - | - | - |
| IC$_{RL}$ | Immediate | - | 45 | - | 44 | - | - | - |
| | Delayed | - | - | - | - | - | - | - |
| CR$_{A}$ | Immediate | - | 31 | - | - | - | - | - |
| | Delayed | - | - | - | - | - | - | - |
| CR$_{S}$ | Immediate | - | 113 | - | - | - | 38 | - |
| | Delayed | - | 40 | - | - | - | - | - |
| CR$_{P}$ | Immediate | 25 | 45 | - | 31 | 27 | - | - |
| | Delayed | 46 | 35 | - | - | - | - | - |
| TR$_{P}$ | Immediate | - | 31 | - | 63 | - | 27 | - |
| | Delayed | 51 | - | - | - | - | - | - |
| SS | Immediate | - | 38 | - | 45 | - | - | - |
| | Delayed | 26 | - | - | - | - | - | - |
| External capsule | Immediate | - | 80 | - | - | - | - | - |
| | Delayed | - | - | - | - | - | - | - |
| SLF | Immediate | - | 101 | - | 94 | 2 | 17 | 88 |
| | Delayed | - | 123 | - | - | - | - | - |
| FO$_{I}$ | Immediate | - | 61 | - | - | - | - | - |
| | Delayed | - | - | - | - | - | - | - |

### B. LHD group

| | | Identity vs. Location vs. Action | | | | | | |
|---|---|---|---|---|---|---|---|---|
| Region | Test phase | I | L | A | I+L | I+A | L+A | I+L+A |
| Insula | Immediate | - | - | 93 | - | - | - | - |
| | Delayed | - | - | 101 | - | - | - | - |
| Parahippocampal g | Immediate | - | - | 107 | - | - | - | - |
| | Delayed | - | - | - | - | - | - | 109 |
| Calcarine | Immediate | - | - | 333 | - | - | - | - |

| | | | | | | | | |
|---|---|---|---|---|---|---|---|---|
| | Delayed | - | - | - | - | - | - | 333 |
| Lingual | Immediate | - | - | 726 | - | - | - | - |
| | Delayed | - | - | - | - | - | - | 729 |
| Inferior occipital g | Immediate | - | - | 92 | - | - | - | - |
| | Delayed | - | - | - | - | - | - | 92 |
| Fusiform g | Immediate | - | - | 203 | - | - | - | - |
| | Delayed | - | - | - | - | - | - | 205 |
| $IC_{AL}$ | Immediate | 34 | - | - | - | - | - | - |
| | Delayed | 26 | - | - | - | - | - | - |
| $IC_{PL}$ | Immediate | 27 | - | - | - | - | - | - |
| | Delayed | - | - | - | - | - | - | - |
| $CR_{A}$ | Immediate | - | - | 36 | - | - | - | - |
| | Delayed | - | - | 65 | - | - | - | - |

Voxel clusters in which the existence of damage significantly affected the following measures of visual memory, using the coding I = identity, L = location, A = action, for the following functions: identity only, location only, action only, identity plus location, identity plus action, location plus action, and identity plus location plus action. Brain regions defined by the AAL and WM atlases with clusters of ≥ 25 'significant' voxels correspond to a z score of 2.6, ($p \leq .005$) are presented. RHD/LHD = right/left hemisphere damage; g = gyrus; $TP_{S/M}$ = superior/middle part of the temporal pole; $CR_{S/A/P}$ = superior/anterior/posterior part of the corona radiata; $FO_{I}$ = inferior portion of the fronto-occipital fasciculus; $IFG_{OP}$ = opercular part of the inferior frontal gyrus; $IC_{AL/PL/RL}$ = anterior/posterior/retrolenticular limb of the internal capsule; SLF = superior longitudinal fasciculus; $TR_{P}$ = posterior thalamic radiation; SS = sagittal stratum.
